# Single nuclei RNAseq stratifies multiple sclerosis patients into distinct white matter glial responses

**DOI:** 10.1101/2022.04.06.487263

**Authors:** Will Macnair, Daniela Calini, Eneritz Agirre, Julien Bryois, Sarah Jäkel, Petra Kukanja, Nadine Stokar, Virginie Ott, Lynette C. Foo, Ludovic Collin, Sven Schippling, Eduard Urich, Erik Nutma, Manuel Marzin, Sandra Amor, Roberta Magliozzi, Elyas Heidari, Mark Robinson, Charles ffrench-Constant, Gonçalo Castelo-Branco, Anna Williams, Dheeraj Malhotra

**Affiliations:** Roche Pharma Research and Early Development, Neuroscience and Rare Diseases, Roche Innovation Center, Basel, Switzerland; Laboratory of Molecular Neurobiology, Department of Medical Biochemistry and Biophysics, Karolinska Institutet, 17177 Stockholm, Sweden; Institute for Stroke and Dementia Research (ISD), Klinikum der Universität München, Ludwig-Maximilians Universität LMU, Germany; Munich Cluster of Systems Neurology (SyNergy), Munich, Germany; Roche Pharmaceutical Sciences, Pathology department, Roche Innovation Center, Basel, Switzerland; Department of Neurobiology and Aging, Biomedical Primate Research Centre, Rijswijk, The Netherlands; Department of Pathology, Amsterdam UMC-Location VUmc, Amsterdam, The Netherlands; Institute of Anatomy, Rostock University Medical Center, 18057 Rostock, Germany; Department of Biosciences, Medicine and Movement, University of Verona, Verona, Italy; Deutsches Krebsforschungszentrum, Im Neuenheimer Feld 280, 69120 Heidelberg; Department of Molecular Life Sciences, University of Zurich, Winterthurerstrasse 190, 8057, Zurich, Switzerland; Faculty of Medicine and Health Sciences, University of East Anglia, Norwich Research Park, Norwich NR4 7TJ UK; Centre for Regenerative Medicine, Institute for Regeneration and Repair, MS Society Edinburgh Centre for MS Research, The University of Edinburgh, Edinburgh BioQuarter, 5 Little France Drive, Edinburgh EH16 4UU, UK; MS Research Unit, Biogen, Cambridge, MA, 02142, USA

## Abstract

The lack of understanding of the cellular and molecular basis of clinical and genetic heterogeneity in progressive multiple sclerosis (MS) has hindered the search for new effective therapies. Here, to address this gap, we analysed 632,000 single nuclei RNAseq profiles of 156 brain tissue samples, comprising white matter (WM) lesions, normal appearing WM, grey matter (GM) lesions and normal appearing GM from 54 MS patients and 26 controls. We observed the expected changes in overall neuronal and glial numbers previously described within the classical lesion subtypes. We found highly cell type-specific gene expression changes in MS tissue, with distinct differences between GM and WM areas, confirming different pathologies. However, surprisingly, we did not observe distinct gene expression signatures for the classical different WM lesion types, rather a continuum of change. This indicates that classical lesion characterization better reflects changes in cell abundance than changes in cell type gene expression, and indicates a global disease effect. Furthermore, the major biological determinants of variability in gene expression in MS WM samples relate to individual patient effects, rather than to lesion types or other metadata. We identify four subgroups of MS patients with distinct WM glial gene expression signatures and patterns of oligodendrocyte stress and/or maturation, suggestive of engagement of different pathological processes, with an additional more variable regenerative astrocyte signature. The discovery of these patterns, which were also found in an independent MS patient cohort, provides a framework to use molecular biomarkers to stratify patients for optimal therapeutic approaches for progressive MS, significantly advances our mechanistic understanding of progressive MS, and highlights the need for precision-medicine approaches to address heterogeneity among MS patients.

## Main

Although we have highly effective therapies for the early inflammatory relapsing-remitting phase of multiple sclerosis (MS), we lack such therapies for the neurodegenerative progressive phase. Therapeutic strategies that have been tested in clinical trials include enhancing neuroprotection directly and enhancing remyelination resulting in indirect neuroprotection by restoring metabolic support and saltatory conduction to the demyelinated axon^1^. However, in spite of promising preclinical data, such trials have not so far resulted in improvement in clinical disability, even though subgroup analysis has shown some promise (e.g. MS-SMART^2^, Opicinumab^3^, Bexarotene^4,5^, Clemastine^6^). This translational mismatch may result from the diversity of disease response in people with MS: within both primary and secondary progressive MS (PPMS and SPMS) clinical subtypes, there is a clear heterogeneity of clinical course, with some people with MS never reaching the progressive disability phase, while others rapidly become disabled. This diverse disease course is difficult to predict at disease onset. Speculating that a heterogeneous neurodegenerative and/or neuroregenerative response to MS pathology between patients underlies these differing disease outcomes, we and others have, in previous work, identified cellular heterogeneity in MS using single nucleus transcriptomics, albeit in a limited number of patients and few pathological MS lesion types^7–9^. However, these studies had insufficient samples to characterize inter-patient heterogeneity of demyelinated lesions and intra-patient heterogeneity. To address this critical gap, we performed a single nucleus RNA sequencing (snRNAseq) study on the most extensive cohort of MS patients to date (Fig. 1a), including both white matter (WM) and grey matter (GM) areas. Our goals were firstly to identify the basis of heterogeneity by comparing cellular compositions and cell type-specific gene expression signatures across WM and GM MS lesion types. Secondly, we sought to identify pathologically relevant ways of stratifying patients on the basis of these responses, so as to better find and test potential therapies for progressive MS.

**Figure 1:**
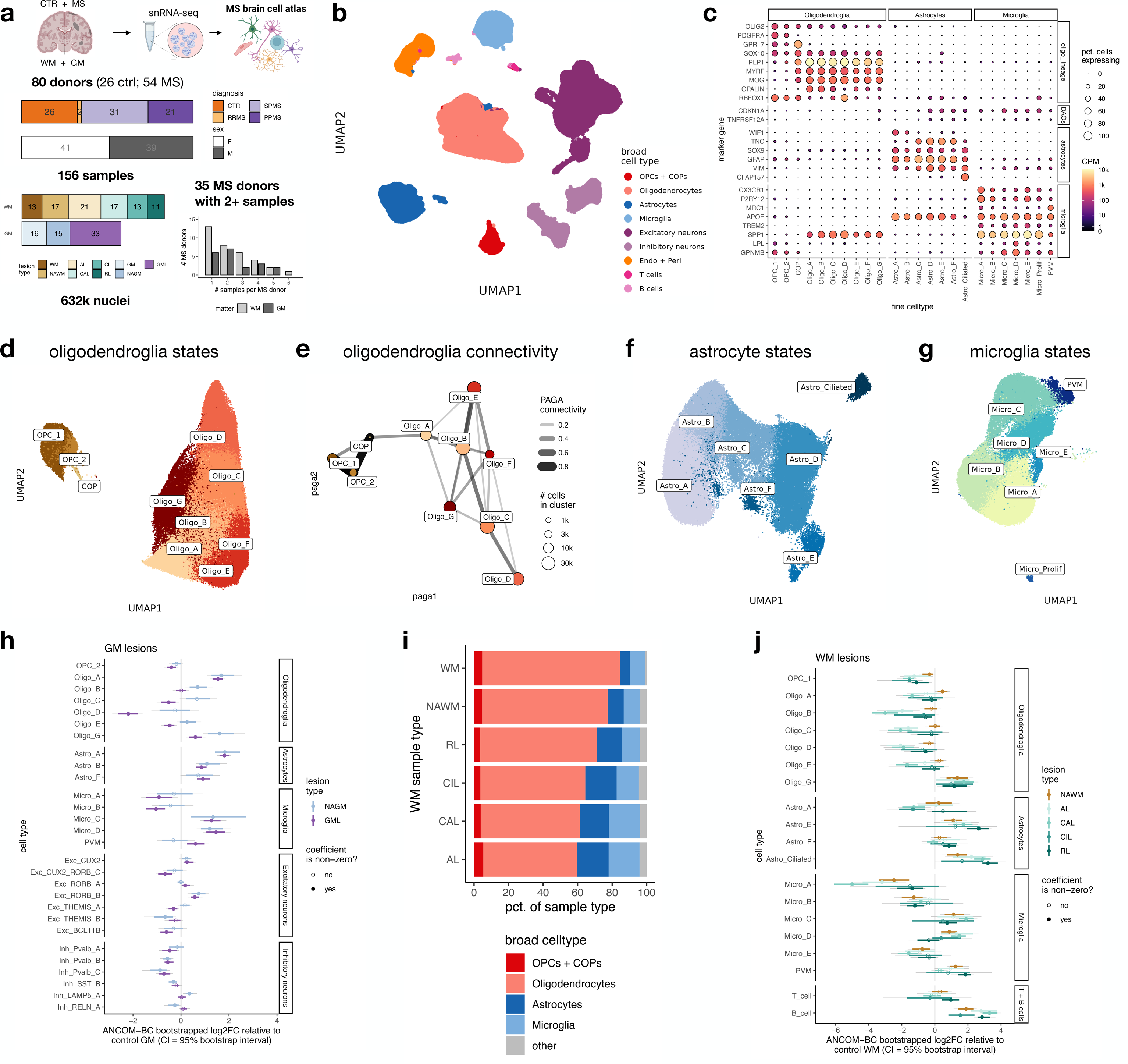
Single cell dissection of cellular heterogeneity in multiple sclerosis lesions. **a** Overview of the donor and sample characteristics (CTRL=control, RRMS=relapsing-remitting MS, SPMS=secondary progressive MS, PPMS=primary progressive MS). **b** UMAP plot of cell types. (Endo +Peri = endothelial cells and pericytes). **c** Dot plot showing canonical markers of broad glial types and their subclusters. (DAOs=disease-associated oligodendrocytes). **d** UMAP plot restricted to oligodendroglia showing subclusters. **e** PAGA applied to oligodendrocyte and OPC / COP subclusters across all samples (edges with weights below 0.2 not shown). **f** UMAP plot restricted to astrocytes showing subclusters. **g** UMAP plot restricted to microglia showing subclusters. **h** Differential abundance of GM MS samples against control GM, as calculated by bootstrapped ANCOM-BC (see Methods and Supplementary Note). Model accounts for sample layer position, by using formula *count ∼ lesion_type + sex + age_scale + pmi_cat + layer_PC1 + layer_PC2 + layer_PC3 + layer_PC4* (where *age_scale* is age at death, normalized to have SD = 0.5^56^). Line at 0 corresponds to no difference between healthy and MS. Points correspond to median log2FC effect estimated by ANCOM-BC; coloured range is 80% bootstrapped confidence interval, grey range is 95% CI. Points are filled when the 95% CI excludes zero; otherwise empty. **i** Pseudobulk analysis showing percentage (pct) of broad cell types in each sample group. **j** Differential abundance of WM MS samples against control WM, as calculated by bootstrapped ANCOM-BC as in **h**. Model fitted is *count ∼ lesion_type + sex + age_scale + pmi_cat*. Abbreviations: WM=white matter, GM=grey matter, NAWM=normal appearing white matter, NAGM=normal appearing grey matter, AL=active demyelinated lesion, CAL=chronic active demyelinated lesion, CIL=chronic inactive demyelinated lesion, RL=remyelinated lesion, GML=grey matter demyelinated lesion.

### Diverse neural cell subtypes observed in brain WM and GM in MS and controls

We profiled 173 WM and GM samples, including (pre QC) >950,000 nuclei from 55 MS cases and 30 controls. Our cohort was similar for age, gender and post-mortem interval between MS and controls (Extended Data File 1, Extended Data Fig. 1a,b, Extended Data Table 1). After randomization of samples during library preparation and sequencing to minimise batch effects, followed by doublet removal, cell and sample QC, including using CellBender^10^ to reduce ambient RNA (Methods), we obtained 632,375 single-nucleus transcriptomes from 156 QC-passed samples, including 506,594 nuclei from MS patients and 125,781 nuclei from controls, profiled at a median depth of 3,810 nuclei/sample, 3,508 reads/nucleus and 1,826 genes/nucleus (Extended Data Figs. 1c-d, Extended Data File 1). These comprised 62 WM lesions (21 active, 17 chronic active, 13 chronic inactive and 11 remyelinated; respectively AL, CAL, CIL, RL), 17 adjacent normal-appearing white matter (NAWM) regions from MS patients, and 13 cortical hemisphere WM regions from non-neurological controls. In addition, we profiled 33 subpial cortical GM demyelinated lesions (GML), 15 adjacent normal appearing grey matter (NAGM) regions from MS patients and 16 cortical GM tissues from controls, all defined as per classical neuropathology^11^, thereby creating a comprehensive atlas of single-nuclei MS transcriptomes (Fig. 1a). We performed pre-processing, integration and clustering via two distinct pipelines (Methods; integration performed with Harmony^12,13^ and Conos^14^), and the clusters identified showed high agreement (Extended Data Fig. 2a, Extended Data File 1). All clusters had acceptable QC metrics, and no cluster was composed of nuclei captured only from individual patients, samples, lesion types or technical covariates, indicating that data integration was successful (Extended Data Fig 2b, Extended Data Fig 2c). This captured all major cell types of the human cortical GM and WM (Fig. 1b) identified using canonical markers (Fig.1c, Extended Data Fig. 2d), derived from both MS and control donor samples (Extended Data Fig. 2e). There was regional and disease-related heterogeneity and we found 59 distinct batch-corrected fine cell type clusters (Methods), including 14 subtypes of cortical excitatory neurons (across layers 2-6), 12 of inhibitory neurons, 11 of oligodendroglia, 7 of astrocyte, 7 of microglia / macrophages, 7 blood vessel-related cells (including 4 endothelial cell and 1 pericyte clusters), and B cell and T cell subpopulations (Extended Data Fig. 2d, Extended Data File 3); we also identified 9 small clusters with mixed lineages which were potentially doublets, and were therefore not considered further (Extended Data Fig. 2d, Extended Data File 3).

We interrogated the fine cell type clusters in turn. Based on expression of previously-described genes characterising oligodendroglia^7,15,16^, we identified 2 oligodendrocyte precursor cell (OPC, *PDGFRA+*, *CSPG4*+, *PTPRZ1*+), 1 committed oligodendrocyte precursor (COP, *GPR17+, BCAS1+*) and 7 oligodendrocyte populations (Fig.1c,d, Extended Data Fig. 3a, Extended Data File 3). (Example of GPR17 validation for COPs shown in Extended Data Fig3b). Analysis of connectivity with PAGA^17^ found a putative main trajectory from OPCs to COPs, followed by Oligo_A, then Oligo_B to C and finally Oligo_D. Markers of this pathway were suggestive of classical oligodendrocyte differentiation, leading from Oligo_A immature markers (e.g. *PLP+*), through B and C, to Oligo_D oligodendrocytes with most myelin protein transcripts (e.g. *MOG+* along with *RBFOX1* and *KLK6*) (Fig. 1c,e, Extended Data Fig. 3a, Extended Data File 3). However, there is, in both MS WM and control WM, an additional branch point to the large cluster Oligo_E, expressing immature markers as well as transcripts relating to cell morphology, cholesterol synthesis and active metabolism (e.g. *FCHSD2*, *ABCG1*^18^*, SFRP1*) (Extended Data Fig. 3a, Extended Data File 3). We also found 2 additional branches leading to either Oligo_F or G, both of which are disease-associated (DA) (see below), expressing transcripts related to the interferon response (e.g. *IRF9*) (Extended Data Fig. 3a). Oligo_G expresses transcripts related to heat shock protein and chaperone protein folding responses (e.g. *HSP90AA1*) and *CDKN1A* and *TNFRSF12A*, similar to the DA2 clusters described recently in mouse^19^ (Fig.1c). Oligo_F expresses transcripts related to DNA damage and injury (e.g. *TOP2A*) (Extended Data Fig. 3a).

Astrocytes (Fig. 1f) divided into grey matter (GM) and WM types, expressing *WIF1, ETV5* (GM, Astro_A-B) and *TNC, ID3* (WM, Astro_C-E), similarly to mouse^20^ (Fig. 1c, Extended Data Fig. 3c). GM astrocytes are more involved in synapse function (e.g. *CHRDL1*) and phagocytosis (e.g. *MERTK*), whereas WM astrocytes are more involved with blood brain barrier (BBB) function and water transport (e.g. with higher expression of *MFSD2A* and *AQP4*) (Extended Data Fig. 3c, Extended Data File 3). Astro_D-E are more reactive, expressing higher levels of *GFAP, VIM, and NES* (Fig.1c). A clear cluster of ciliated astrocytes was present (*CFAP157+, DNAH9+*) (Fig. 1c,f, Extended Data Fig. 3c).

Similarly to signatures identified in recent microglial datasets^21^, we observed microglial subclusters expressing transcripts associated with homeostasis, in addition to more reactive subclusters (Fig.1 c,g). Micro_A and Micro_B appear to be homeostatic, showing relatively high expression of *P2RY12* and *CXCR1*, while Micro_C-E had a more reactive phenotype, expressing higher levels of MHC class II molecule transcripts (e.g. *HLA-DPB1*), transcripts such as *TREM2* and *APOE*, but also *PLIN3* (lipid accumulation) (Fig. 1c, Extended Data Fig. 3d, Extended Data File 3). Some microglia expressing these markers have been described in other neurodegenerative diseases in mouse and human, sometimes termed disease-associated microglia (DAM) or microglia inflamed in MS (MIMS)^21^ and Micro_D and E most closely map onto these phenotypes. Perivascular macrophages (PVM) / border associated macrophages were also detected, expressing high levels of transcripts such as *MRC1* and *LYVE1* (Fig. 1c,g, Extended Data Fig. 3d).

Neuronal heterogeneity reflected the subtypes of excitatory and inhibitory neurons found in cortical layers in human GM, including for excitatory neurons, *CUX2*+ neurons from layer 2, *RORB*+ neurons from layer 3 and 5, and *TLE4*+ neurons in the lower layers (5/6). Inhibitory GABAergic neurons subdivided by location e.g. *RELN*+ neurons (layer 1), and by neurotransmitter e.g. *PVALB+, SST+, VIP+* neurons (Extended Data Fig. 2d).

As a result, our dataset, which uses snRNA-seq to study the largest cohort of MS patients and lesions to date, describes a wide range of homeostatic and disease-associated cell states. We further examined these in terms of their composition and transcriptional changes in both lesions and patients.

### Distinct compositional differences in WM and GM lesions

Having defined our cell type subclusters, we next investigated the compositional differences between MS and control samples. We focused first on GM samples, as we and others have shown that there is a loss of PVALB+ and SST+ inhibitory neurons in MS GM^22,23^ as well as some types of excitatory neurons^8,24^. We reproduced these findings (Fig. 1h), with effects similar between GM lesions (GML) and normal appearing GM (NAGM), as before, but more pronounced in GML. Consistent with astrogliosis, GM Astrocyte clusters Astro_A, Astro_B and Astro_F are increased in MS in GM, both in NAGM and GML (Fig. 1h). As expected, oligodendrocyte clusters Oligo_C and Oligo_D (expressing most myelin protein transcripts) are reduced in GML compared to NAGM and control. However, disease-associated oligodendrocytes (Oligo_G) are increased. Immature oligodendrocytes (e.g. Oligo_A) are increased in abundance in GML and NAGM, and there is increased Oligo_B and Oligo_C abundance in NAGM (but not GML), consistent with the described regenerative response to demyelination in GML, generated in surrounding NAGM tissue and which is more successful than in WM^25^ (Fig. 1h)..

Turning to WM, classical pathology descriptions divide WM MS demyelinated lesions by the pattern of immune infiltrate into active (AL), chronic active (CAL), and chronic inactive (CIL), with additional remyelinated lesions (RL) (definitions in Methods, Extended Data Fig. 4). Analysis of sample compositions at the broad cell type level confirmed the changes in overall neuronal and glial numbers previously described within the classical lesion subtypes: the expected reduction in oligodendrocytes in MS most pronounced in demyelinated lesions, increase in microglia/macrophages especially in AL and CAL (which is what defines these lesions), and an increase in astrocytes in MS, compared to NAWM and control WM (Fig. 1i). However, more detailed differential abundance analysis at the cell type subcluster level reveals additional differences in subcluster patterns (Fig. 1j). While most oligodendrocyte subtypes were reduced in MS WM, there was an increase in immature oligodendrocyte type Oligo_A confined to NAWM, (unlike in GM, reiterating the different environments of GM and WM for oligodendrocyte regeneration), and an increase in Oligo_G (disease-associated oligodendrocytes). There was also a change in microglia from a more homeostatic phenotype (Micro_A,B) to a more reactive type (Micro_C,D) a change of astrocytes from a more homeostatic phenotype (Astro_A) to a more reactive phenotype (Astro_E,F) and an increased in ciliated astrocytes (Astro_ciliated) (Fig. 1j). These differences were generally present in most MS samples compared to controls, but with NAWM samples being more similar to remyelinated lesions and controls, and AL and CAL being similar but with the biggest differences relative to non-MS samples. Repeat analysis with the orthogonal technique Milo^26^ showed broadly similar results (Extended Data Fig. 5a,b).

### GM and WM lesions have distinct cell type-specific transcriptional responses

The cell composition analysis indicates that there is considerable variation for specific glial cell types in WM and GM lesion samples (Fig. 1h-j and Fig. 2a,b). We then investigated whether there was variation between donors and notably we also found significant differences in many glial types (12 out of 23 glia types; FDR < 0.05 for donor effect) in WM samples (Fig. 2a), and some (5 out of 17; FDR < 0.05 for donor effect) in GM samples (Fig.2b). These results highlight the relevance of disease-associated glial cell states to neuropathology in MS. To understand this further, we next investigated the differential gene expression (DEG) between cells in different lesion environments, taking in account donor variation. We identified gene expression changes for each broad cell type between WM lesions and control WM tissue, and between GM lesions and control GM tissue using a mixed model (glmmTMB^27^) fit to pseudobulk data, including age, sex and post-mortem interval (PMI) as possible confounding variables, and donor ID as a random component (Methods, Supplementary Note). Indeed, the fitted models for all broad cell types showed strong donor effects for many genes (Extended Data Fig. 6a,b). Nevertheless, we identified 5,106 DEGs in WM, and 4,824 DEGs in GM across all major cell types (Fig. 2c, Extended Data File 4 and https://malhotralab.shinyapps.io/MS_broad/).

**Figure 2:**
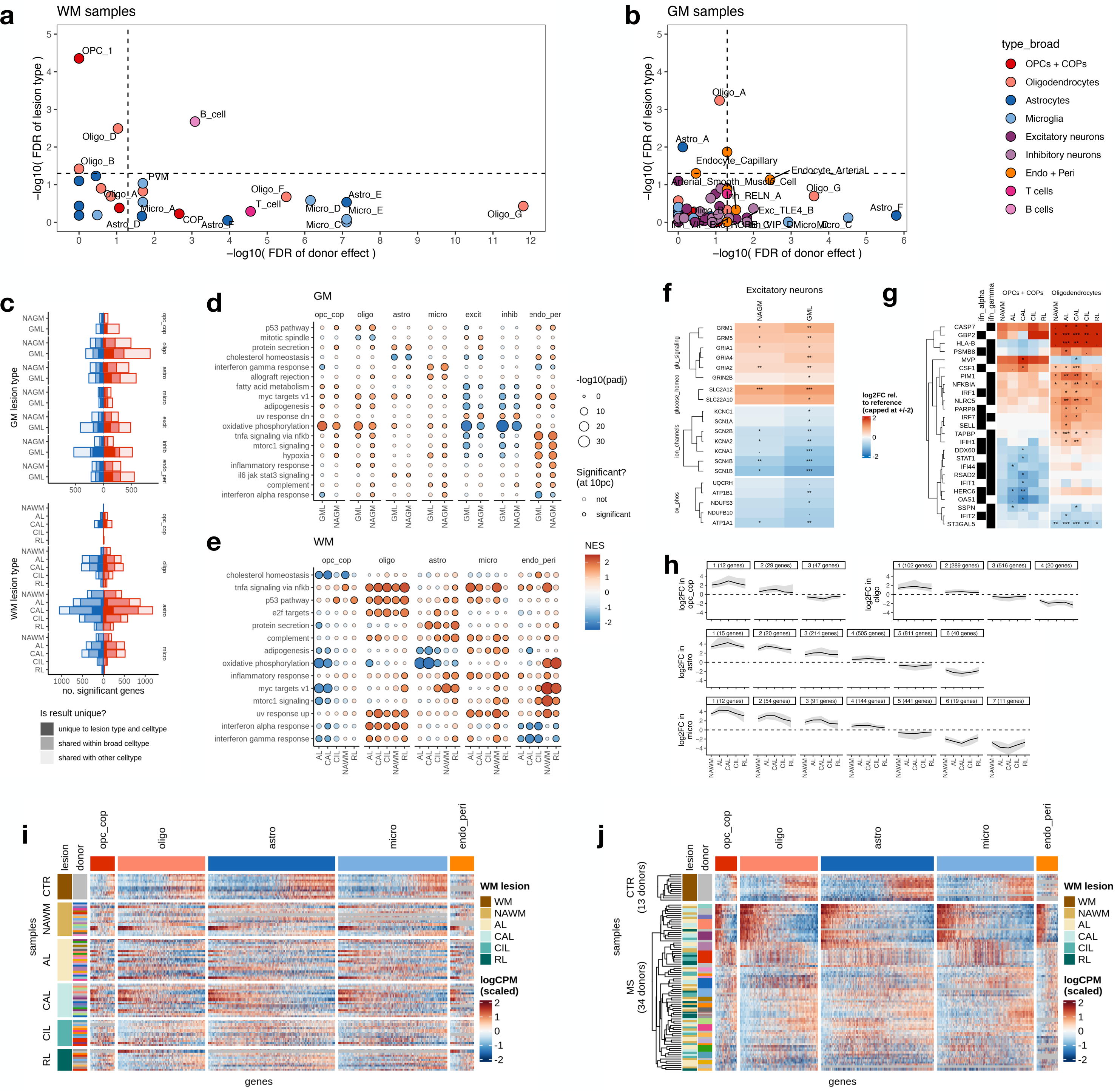
Patterns and determinants of cell type-specific gene expression profiles in WM and GM lesions. a,b. Contribution to variability in cell type abundances explained by lesion plus patient in WM (**a**), and GM (**b**). In **b**, three neuronal layer PCs are included as confounders. FDR is Benjamini Hochberg-corrected p-values from likelihood ratio tests of nested models. Axes show evidence that lesion type (y axis) or donor effect (x axis) significantly improve the fit of a model explaining cell type proportion of a sample. **c** Barcharts showing number of significant differentially expressed (DE) genes for WM and GM tissue and cell type, only DE for this cell type, or also DE in another cell type. **d,e** Dotplots of top GO terms plotted for broad cell types and tissue (GM **d**, WM **e**). Key is shared. **f** DE genes related to glutamate signalling, glucose homeostasis, ion channels and oxidative phosphorylation in excitatory neurons in GM lesions (Mean log2 CPM). **g** Differential expression of interferon alpha and gamma genes in oligodendroglial cells in WM lesions (Mean log2 CPM). **h** WM gene expression fold change profiles over lesion types for each broad cell type showing continuous patterns (opc_cop = OPCs + COPs; oligo = oligodendrocytes; astro = astrocytes; micro = microglia). Restricted to genes where at least one lesion type has FDR <15%, hierarchical clustering with cut distance set to log(4), and clusters with fewer than 10 genes not shown. Figure in brackets shows number of genes in the cluster. **i** Pseudobulk expression heatmap of genes showing either MS or donor variability in broad cell types in WM, samples ordered by lesion type. **j** Pseudobulk expression heatmap as in **i**, row order on basis of hierarchical clustering, showing differences between MS donors not explained by lesion type, sex, or type of MS, but all samples from one donor cluster together, suggesting a strong donor effect. **i,j** Column order determined by the first principal component of the gene matrix for each cell type. Abbreviations: WM=white matter, GM=grey matter, NAWM=normal appearing white matter, NAGM=normal appearing grey matter, AL=active demyelinated lesion, CAL=chronic active demyelinated lesion, CIL=chronic inactive demyelinated lesion, RL=remyelinated lesion, GML=grey matter demyelinated lesion, opc_cop = OPCs and COPs; oligo = oligodendrocytes; astro = astrocytes; micro = microglia.

In MS WM and GM, all cell types showed strong changes in gene expression, more marked in demyelinated lesions than normal appearing matter (Fig. 2c), with gene ontology analysis (Fig. 2d,e) indicating several altered pathways. In GM, both excitatory and inhibitory neurons showed more DEGs in GML compared to NAGM, predominantly in selectively vulnerable neuronal cells (*PVALB*+ interneurons and upper/mid layer excitatory neurons) (Extended Data File 3 and https://malhotralab.shinyapps.io/MS_fine/). Focusing on MS excitatory neurons (residing in layer II/IV), there was upregulation of genes related to glutamate signalling (e.g. *GRIA1,2,4, GRIN2B, GRM1,5*), glucose or cation homeostasis (*SLC2A12, SLC22A10*) with concurrent down-regulation of specific ion channels (*SCN1A, SCN1B, SCN2B, SCN4B, KCNA1, KCNA2, KCNC1*) and oxidative phosphorylation (OXPHOS) genes (*ATP1A1, ATP1B1, NDUFB10, NDUFS3, UQCRH*) (Fig. 2f). This suggests that glutamate excitotoxicity in excitatory neurons in combination with fewer inhibitory neurons (Fig. 1h) leads to increased excitatory and decreased inhibitory tone which may be critical in MS GM pathology, and therefore may be a promising therapeutic target in progressive MS.

In WM lesions, consistent with our prior observations^16,28^, expression of genes involved in interferon alpha and gamma responses varied across MS lesions and often showed opposite patterns in OPCs compared to oligodendrocytes (Fig. 2g). Genes involved in inflammation-related pathways were also enhanced in astrocytes, microglia and oligodendrocytes in WM lesions (Fig. 2e). However, we found no patterns of gene expression predictive of lesion type. Within each glial broad type, the majority of DEGs were shared across lesions (Fig. 2e,g). For some such transcripts, we observed ‘u’/’n’-shaped profiles of transcriptional changes along the pathological category of the lesion, with NAWM lesions showing small fold changes relative to control WM, increasing to the largest fold changes in AL and CAL, then decreasing again in CIL and RL (Fig. 2h, Methods). This indicated that, rather than distinct transcriptional differences across MS lesions, there is a continuum of changes, most likely reflecting global neuropathology.

### Donor effects drive cellular and transcriptional heterogeneity in MS brains

The cell-type abundance and gene expression comparisons show distinct changes associated with MS, that are not consistent with distinct signatures for the neuropathologically-defined lesion categories, but rather with a continuum of cellular (Fig. 1h,j) and transcriptional pathology (Fig. 2d-h). What then is driving the cell type and gene expression changes in our dataset? Based on the significant inter-individual variation in cell type composition noted above, we hypothesized that different sub-groups of patients, rather than different lesion types, might share transcriptional pathology. This hypothesis could not be explored in prior bulk RNAseq studies of WM lesions but the unique strength of our study design, with multiple different lesion types from the same patient and the power of single nuclei resolution, enables it to be tested.

For each sample, and in each broad cell type, we analysed the gene expression pattern for genes which either showed significant disease effect, or were highly variable for each sample. As expected from our earlier results, there was no pattern of up or downregulation of genes that correlated with sample neuropathological category or lesion type, in either WML (Fig. 2i) or GML Extended Data Fig. 7a). However, in both WM and GM samples, hierarchical clustering of the same data showed a clear expression pattern which correlated with the donor ID of the samples (Fig. 2j, Extended Data Fig. 7b). This provided strong evidence for cross-cell type transcriptional similarity within patients. Within an individual patient, the gene expression profiles were remarkably similar across multiple lesion types and normal appearing matter, while subgroups of different patients showed distinct transcriptional profiles (Fig. 2i,j, Extended Data Fig. 7a,b). We concluded that although MS lesions clearly differed by cellular composition, at the gene expression level, cells within both WM and GM lesions appeared more affected by donor identity rather than the lesion environment.

### Coordinated multicellular gene expression programs define patient subgroups

This demonstration that patients with long-standing MS differ markedly in the transcriptional signatures of their glia, whatever their lesion classification, but which fall into apparent subgroups, is important in the context of the variable responses to experimental neuroprotective therapies for example targeting remyelination. To explore these apparent subgroups more, and to understand the underlying cellular and molecular mechanisms, we adapted MOFA+, a computational method originally developed for identifying low-dimensional representations of variation across multiple data modalities^29^ to identify similar donor-associated transcriptional patterns across multiple cell types (modalities) simultaneously^29,30^. For each cell type, we selected genes with evidence of a MS effect and/or a donor effect (Extended Data Fig. 6a,b) to capture both consistent MS pathology and patient-patient variability. This resulted in gene sets largely distinct for each cell type, with some common genes (Extended Data Fig. 8a,b). MOFA+ identified 5 factors each in both GM (GM_F1-5) and WM (WM_F1-5) samples that explained at least 5% of variability for some cell type, with each factor describing one axis of variation in MS. Where the factor explains variance in multiple cell types simultaneously, this represents a cross-cellular response to MS; where the factor explains variance mostly in one cell type, this program is most influential in that cell type.

In GM samples, for all factors except factor GM_F1 and GM_F3, there was considerable overlap between the distributions of control and MS samples (Extended Data Fig. 9a). Factor GM_F1 gene expression could robustly distinguish MS GM pathology (NAGM and GML) from healthy control GM, with MS diagnosis explaining 71% of variability in factor GM_F1 (Extended Data Fig. 9b). Factor GM_F1 explains equivalent variability in gene expression across glia (Extended Data Fig. 9b). Factor GM_F3 distinguishes GML from control, is predominantly neuronal, and high factor GM_F3 is characterized by downregulation of genes related to oxidative phosphorylation, the electron transport chain and protein folding (Extended Data Fig. 9c, Extended Data File 5) indicating altered metabolism and mitochondrial function, as described already in MS brain samples^31,32^.

MOFA+ WM factor scores (WM_F1-5) clearly distinguished MS patients from controls, and stratified MS patients into distinct subgroups (Fig. 3a). Very similar factors were found using the orthogonal method scITD^33^ (Extended Data Fig. 9d, Methods). Four subgroups of MS patients had a distinct pattern of high/low factor scores across factors WM_F1-4, being either factor WM_F1, WM_F2, WM_F3 or WM_F4 high; scores for factor WM_F5 showed more variability across donors (Fig. 3b). These apparent subgroups were not explained by lesion type (Fig. 4a), or any available known metadata including sex, type of MS, age, post mortem delay, brain bank origin (Extended Data Fig. 9e) or differences in technical quality (Extended Data Fig. 9f). To infer potential mechanisms, we examined the genes driving these factors and found that, broadly, factors WM_F1-4 described glial responses to damage and WM_F5 a regenerative response. More specifically, high factor WM_F1 scores were characterised by a pan-glial upregulation of genes involved with protein folding, chaperone proteins and ubiquitination e.g. *HSPB1, HSPA4L, HSP90AA, BAG3, SERPINH1* (as found in other neurodegenerative diseases^34,35^), and reduced microglial expression of *CX3CR1, P2RY12* and *P2RY13* (homeostatic markers), suggesting an adaptive cellular response to stress (Fig. 3c, Extended Data Fig. 9g, Extended Data File 5). High factor WM_F2 genes were characterised by cross-glial upregulation of genes in the integrated stress response, DNA damage, growth arrest and apoptosis pathways, including *GADD45A*, *GADD45B*^36^ and *NAMPT* (Fig. 3d). Factor WM_F3, which affects oligodendroglia most strongly, was characterised by upregulation of extracellular matrix (ECM) genes, including *COL19A1, COL22A1, TNC* and *ITGB4* (Fig. 3e), described to inhibit oligodendrocyte maturation and reduce myelination^37,38^. High Factor WM-F3 was also associated with downregulation of *CRYAB* (alpha-crystallin B) thought protective in demyelination^39^ and of interest due to its protein homology to Epstein Barr virus nuclear antigen 1 with its association with MS diagnosis^40^. Factor WM_F4 also affected oligodendroglia most strongly, but was characterised by upregulation of MHC class 1 molecules (*HLA-B* and *HLA-C*), previously described as upregulated in oligodendroglia in MS lesions and in preclinical mouse models of EAE targeting them for destruction^16,28^, as well as the immune-related gene *ARHGAP24* described before as expressed in oligodendrocytes^41^. In addition, WM_F4 was associated with genes whose upregulation is associated with efforts to increase oligodendrogenesis, oligodendrocyte differentiation and myelination (e.g. SFRP1^42^, and *ANGPT2* similar to *ANGPTL2*^43^), (Fig. 3f). Finally, factor WM_F5, affecting only astrocytes, was characterised by a strong upregulation of genes relating to primary cilia, including *DNAH11+6, SPAG17, ZBBX, and CFAP54* (Fig. 3g). Ciliated astrocytes are thought to be pro-regenerative^44^, although may become longer and dysfunctional in disease^45–47^.

**Figure 3:**
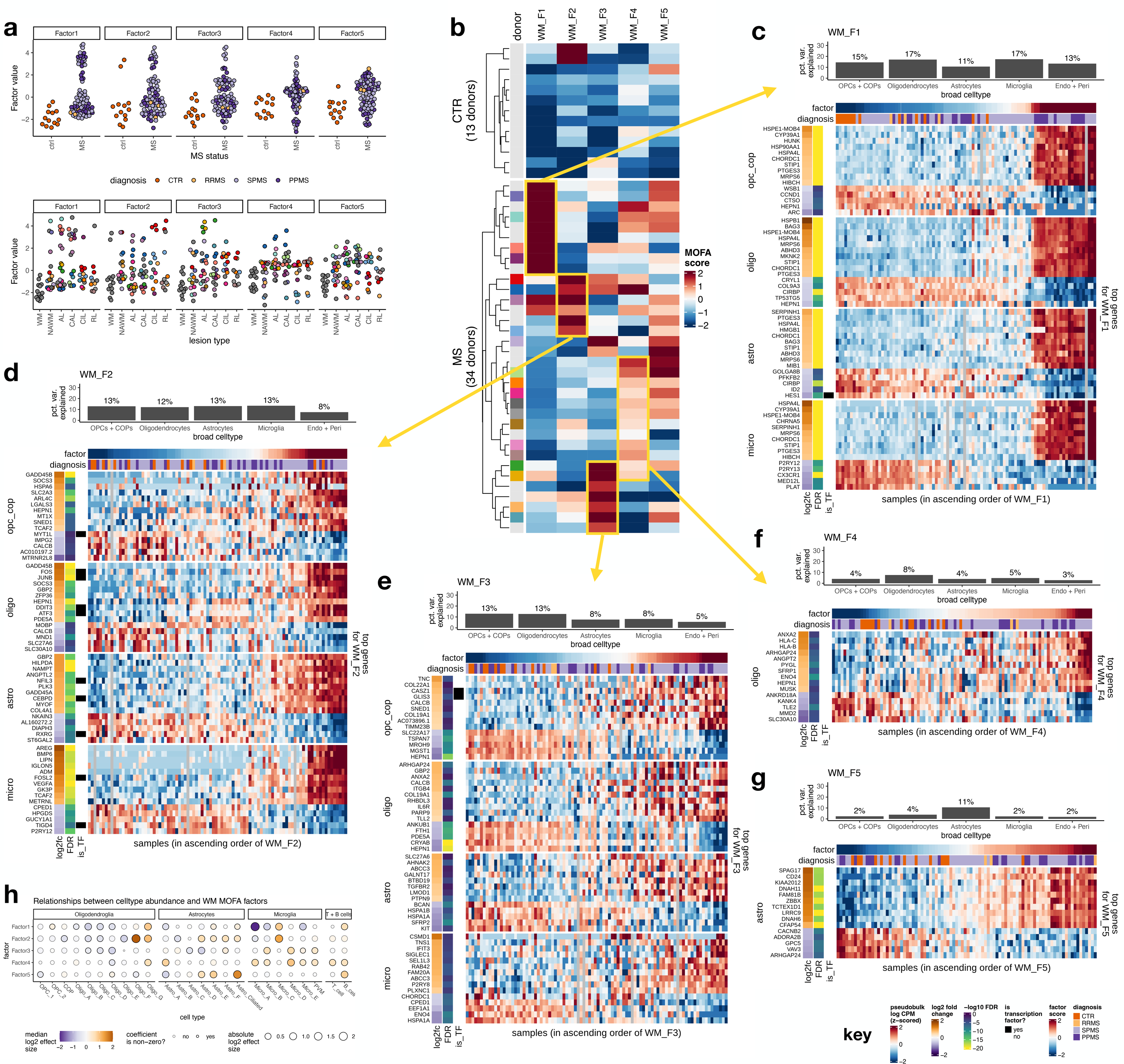
Patient stratification by WM factor values. **a** MOFA+ WM factor values (WM_F1-5) for control and MS samples, showing no correlation with WM lesion type. **b** Heat map of MOFA+ factors (columns) with signs changed to positively correlate with MS status, with donors in rows. Yellow boxes point to **c-g** showing genes driving each factor including bar plot showing percentage (pct) variance explained by the factor in each broad cell type, and heatmap of 15 genes with largest absolute Factor weights for each cell type (10 increased, 5 decreased) where >= 10% variance is explained by Factor. Columns are samples, ordered in increasing order of Factor score (top bar). Rows are genes, split by broad cell type. Heatmap colours are log CPM of pseudobulk expression, z-scored within each row; grey indicates insufficient cells of this cell type in the sample. Column annotations show log2fc, FDR and whether the gene is a transcription factor. **c** WM Factor1, **d** WM Factor 2, **e** WM Factor 3, **f** WM Factor 4, and **g** WM Factor 5. Key bottom right. **h** Dotpot showing relationship between cell type abundance of glial and immune cell subclusters and WM MOFA factors.

**Figure 4:**
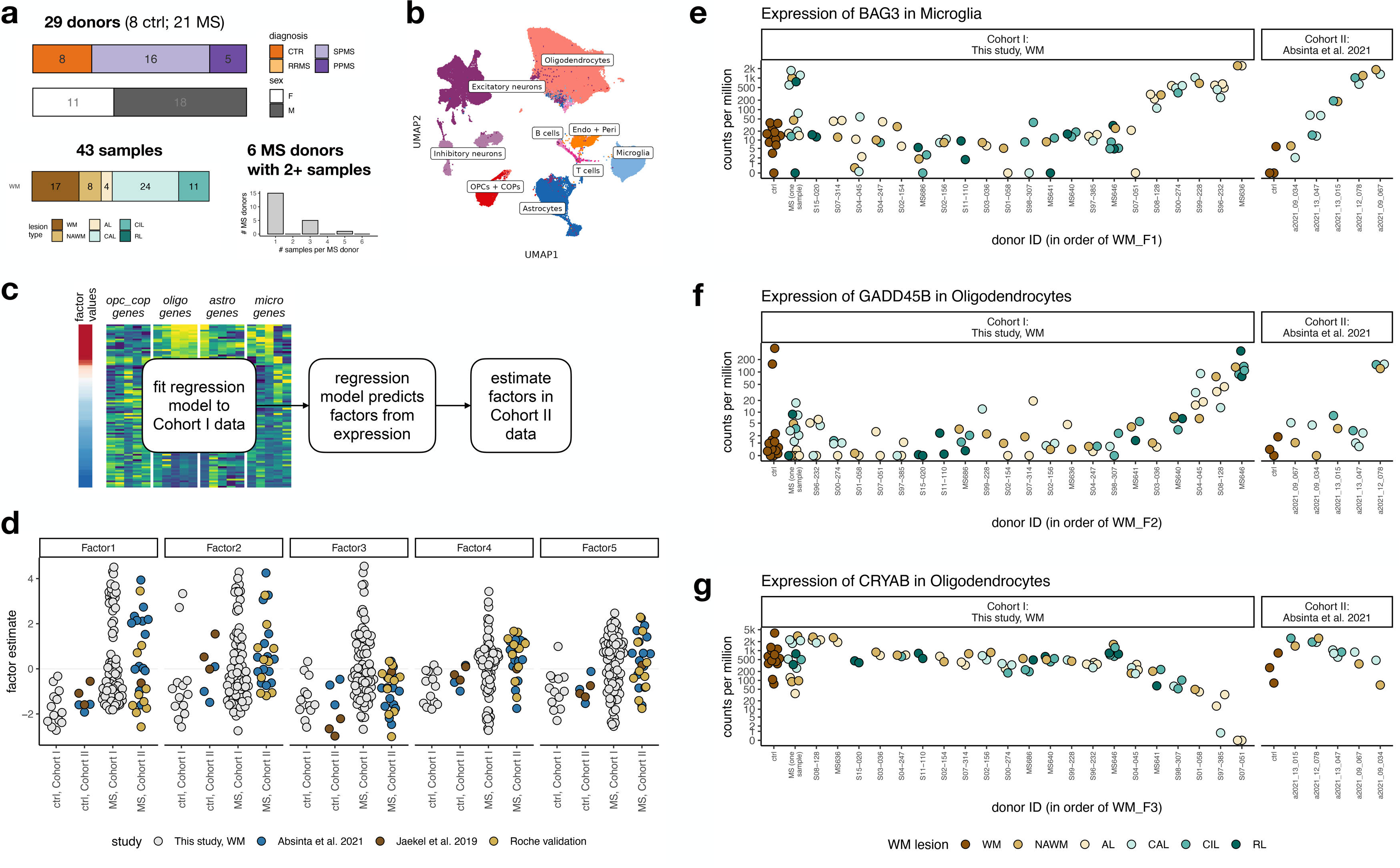
Second cohort dataset for validation. **a** Overview of Cohort II donor and sample characteristics (CTR=control, RRMS=relapsing-remitting MS, SPMS=secondary progressive MS, PPMS=primary progressive MS). **b** UMAP plot of cell types of Cohort I and II combined to show similarity. (Endo + Peri = endothelial cells and pericytes). **c** Cartoon of how the regression model is applied to data for factor estimates. **d** Factor estimates for cohort II compared to cohort I factor values coloured by dataset. **e-g** Gene expression of selected genes related to WM factors in selected broad cell types, in controls, single samples per donor for datasets, and multiple samples per donor for Cohort I and Cohort II, ordered by WM factor level expression, **e** *BAG3* (WM_ F1), **f** *GADD45B* (WM_F2), **g** *CRYAB* (WM_F3).

To further investigate whether these WM factors could describe different degenerative and regenerative pathological responses, we compared glial cellular compositional changes in samples and WM factor values (Fig. 3h). High factors WM_F1-3 were associated with fewer oligodendrocyte types Oligo_B-D, consistent with general loss of oligodendrocytes and/or conversion to other more disease-related types. High factor WM_2 was associated with increases in Oligo_F and G (high in stress genes), reactive astrocytes (Astro_D-F), and reactive microglia (Micro_C), whereas high factor WM_F1 was associated with a marked reduction in the homeostatic microglia Micro_A, and to a lesser extent Micro_B and E. Oligo_G was increased in factors WM_1, 2 and 4. In addition, increased WM_F4 (affecting oligodendroglia most strongly) showed a reduction in Oligo_D, the subtype making most myelin transcripts, with no reduction in the immature subtypes. As Factor 4 is associated with genes related to increased oligodendrocyte differentiation and myelination, this may suggest that the regenerative response is mounted but blocked and maturation fails. The increase in microglial subtypes Micro_A and B (with homeostatic markers) with increasing WM_F4, may also suggest a attempted regenerative response, since oligodendroglial crosstalk with specific microglial populations is essential in effective regeneration^48,49^. High factor WM_F5 (affecting astrocytes most strongly) was markedly associated with ciliated astrocytes (Astro-Ciliated), with their proposed beneficial effect^44^. Thus, factor-based analysis correlated with differential cell type composition and allowed us to stratify the majority of MS donors by WM pathology phenotype: Type 1 (Stressed - chaperone response), Type 2 (Stressed - DNA damage), Type 3 (inhibitory ECM response), Type 4 (immune and attempted regenerative oligodendrocyte response) in combination with a more variable expression of a regenerative astrocyte response. Therefore, this stratification correlates with differential cell type composition.

### Validation of proposed patient stratification in other MS cohorts

To investigate whether the patient stratification identified by our MOFA+ approach was unique to our MS dataset (Cohort I) or could also stratify other MS patients, we analyzed an independent snRNA-Seq cohort (Cohort II) of 43 WM samples from 21 MS donors and 8 control donors (Fig. 4a). Using the same workflow as before (methods), after QC, we were able to include 10 new WM MS samples (named here as Cohort IIa), and WM samples from published datasets (Cohort IIb, including 14 MS samples and 3 controls from the Absinta et al., 2021 dataset^9^ and 1 MS donor and 3 controls from our previous Jaekel et al., 2019 dataset^7^). The Absinta et al., 2021 dataset^9^ contained 5 MS donors each with 2-4 samples whereas the rest were single samples from single donors. Datasets containing mixed white and grey matter samples could not be includeds (e.g. ^8^) as we aimed to verify that donors could be stratified according to our WM Factors and that the donors accounted for more of the variability in cell type-specific gene expression than the WM lesions.

The integration of the new Cohort II with the Cohort I showed the same broad cell clusters (Fig. 4b). We then implemented a regression model that allows us to estimate factors values acquired from a snRNA-Seq cohort to any new snRNA-Seq cohort dataset (which could be MS or any other neurological disease) (Fig. 4c and Methods). We used this regression model to estimate Cohort I WM factor values in Cohort II (Fig. 4c) and found that the validation samples showed very similar distributions of WM factor values as in the discovery data, with the exception of WM-F3, which we hypothesize relates to the low sample size (Fig. 4d). In addition, visualisation of an example of a gene driving these factors (here WM_F3: *CRYAB*) across multiple samples of the Cohort II dataset indicated similar expression across different lesion types and distinct expression across MS individuals, adding evidence to the global donor effect rather than lesion effect (Fig. 4e). Thus, the pathological phenotypes in distinct subsets of patients were also identified in independent MS cohorts, suggesting that these may be useful in stratifying patients more widely.

## Discussion

This snRNAseq study on the most extensive cohort of MS patients to date, shows that GM and WM biology in MS are fundamentally different at a molecular and cellular level. While GM changes relate to presence of a demyelinated lesion and patient ID, WM MS biology is more complex. Although there are cellular compositional differences present in each of the classical MS WM lesion categories as we would expect, the gene expression patterns of these cells are surprisingly agnostic to the lesion environment and instead are associated primarily with individual patient effects. These global patient effects allow us to take the first step towards the stratification of progressive MS patients by their molecular and cellular pathology, only made possible by the large number of samples captured both within individuals and from different individuals. Recent work has explored the trade-offs between read depth and number of individuals^50^, finding greater read depth useful for characterising lowly expressed genes and rare cell types, but our study suggests that there is also much to be discovered by increasing the breadth of patient samples.

Our WM results point to four fundamentally different neurodegenerative pathological phenotypes in MS, each specific for a subgroup of patients and shared by all WM lesions and NAWM in a single patient: First, where there is a cross-glial stress response with increased protein folding/chaperone/ubiquitination pathways (Type 1 Stressed - chaperone response). Second, where there is an alternative cross-glial DNA damage stress response (Type 2 Stressed - DNA damage). Third, where there is an increased inhibitory ECM response to oligodendroglial differentiation (Type 3 - inhibitory ECM response). Fourth, where there is an immune oligodendrocyte response and a failure in the final stage of oligodendrocyte maturation (Type 4 - immune response). Superimposed on these is a regenerative astrocyte response. None of these phenotypes group with any available patient metadata including age, sex, type of progressive MS, previous medications, post mortem delay or sample quality measures (Extended Data Fig. 9e,f). These pathological phenotypes are reminiscent of the previous work of Lucchinetti et al. who tried to type different patient pathological responses^51^ but without the benefit of current high resolution techniques.

A key prediction of our results is that each patient group expressing one of WM Factors 1-4 will respond best to different neuroprotective/regenerative therapies. This may help explain the apparent poor response of neuroprotective/pro-regenerative therapies in progressive MS trials – beneficial effects may have been missed due to the inability to perform appropriate (Factor based) subgroup analysis. Any positive response in these trials may be ‘diluted out’ by patient heterogeneity, and effective therapies for one subgroup may be lost. We propose that pro-remyelinating drugs acting through increasing oligodendrocyte maturation may be most effective in the patient subgroup Types 3 and 4 where oligodendrocytes stall in differentiation, and not in those subgroups where the need is to reduce cellular stress. It is also conceivable that Siponimod, now approved for selected SPMS patients, may have a more marked effect in the Type 2 (stressed - DNA damage) subgroup, through its proposed role in NRF2 signalling and antioxidant pathways^52^.

To stratify MS patients to give future clinical trials the greatest power to reveal effective therapies, we now need to link these post mortem phenotypes to biomarkers measurable in the living, ideally in serum or CSF, or perhaps even as targets for Positron Emission Tomography (PET) ligands. High protein levels of CSF Parvalbumin correlated with loss of PVALB+ inhibitory cortical neurons and MS disease severity^22^, consistent with our GM findings, but we need accessible biomarkers of our WM phenotypes. This will allow us to determine whether the phenotypes are stable in the same patients over time^53^ and to interrogate clinical trial datasets for effect or indeed lack of effect within post hoc stratified patient groups. In this study, we provide a resource of cell type-specific genes, identified by MOFA factors, whose expression clearly distinguishes WM and GM pathology, and subgroup phenotype to aid such future biomarker efforts. This is an essential step change for designing effective precision medicine therapeutic strategies for progressive MS - a critical unmet need.

## Limitations of the study

A peripheral readout for CNS pathology in MS would be optimal in clinical settings for monitoring of the degenerative phase of the disease with stratification for prognosis and therapies. Our dataset, the largest snRNAseq of MS samples yet, including white and grey matter, and including multiple samples from the same donors, constitutes a first important step in this direction, providing a molecular blueprint of MS neuropathological responses. We anticipate that in the near future, with additional large MS cohorts where snRNA-Seq CNS profiles can be integrated with blood/CSF datasets, this will become possible.

For practical reasons, our study uses snRNAseq from post-mortem archival tissues, with the limitation that this only evaluates primarily pre-mRNA nuclear transcripts. We undersampled rare immune cell populations e.g. activated CD8 T cells, monocytes, dendritic cells, B cells and microglia inflamed in MS (MIMS)^9^ which are mostly enriched at the edges of chronically inflamed WM lesions, perhaps as we focussed on the entire lesion, but the sharing of gene expression programs across all lesions in oligodendrocytes, astrocytes and microglia suggests the generalizability of our results. In addition, we cannot comment on subpial GM lesions adjacent to compartmentalised inflammatory meningeal infiltrates^54^, as these were not in our dataset. We tried to match our MS donors and controls as well as possible, but differences (in banks, PMI etc) remain and differences will exist that we are unaware of. However, no results associated with any available patient metadata. Efforts to process clinical summaries from archival tissue from neurological patients with neurological disorders are now on the way^55^. The integration of such studies with datasets as ours will be required in the future to link snRNA-Seq based stratification with clinical criterias. Although we use a larger cohort of individuals than often used in single cell-omic studies, with all available and suitable public data for validation, even larger numbers will be required to validate our findings of patient groupings based on factor analysis at a transcriptional level, especially the high in factor WM-F3 group. Increasing the scope of our analysis with other modalities, for instance at the proteomics, lipidomics and epigenomics level in future studies will help in the further characterization and stratification of MS patients.

## Methods

### Overview of workflow

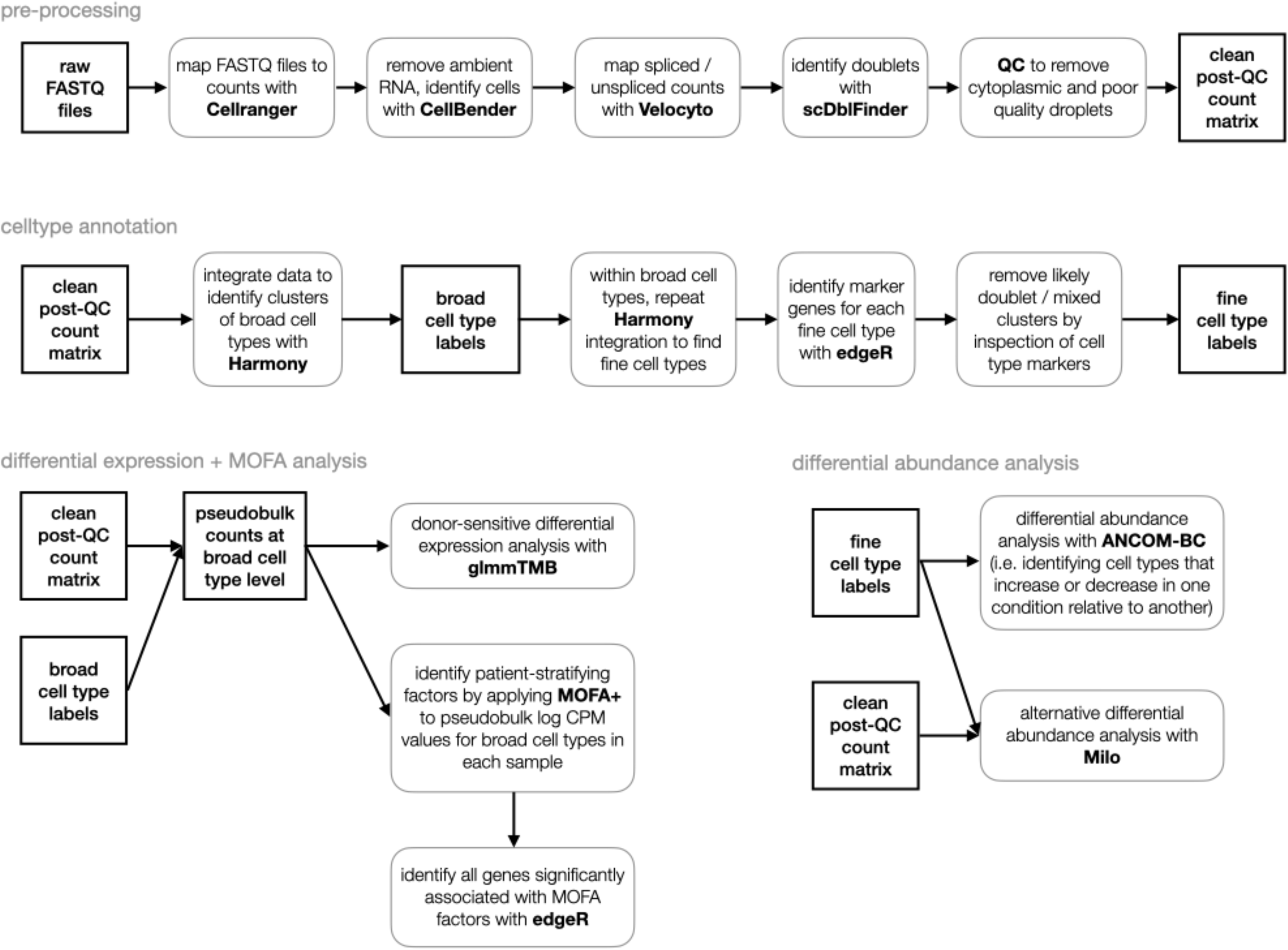

### Sample preparation and single nuclei RNA sequencing

#### Brain tissue samples, ethical compliance and clinical information

Human tissue samples were obtained from the Netherlands Brain Bank (NBB), the MS UK tissue bank (UKTB) and the Edinburgh Brain Bank (EBB) via donor schemes with full ethical approval from respective brain banks (METc/2009/148 from Medical Ethical Committee of the Amsterdam UMC, MREC/02/2/39 from UK Ethics Committee), and individual material transfer agreements between Roche and ABB, UKTB and EBB. We have complied with all relevant ethical regulations regarding the use of human postmortem tissue samples. In the discovery dataset, we examined a total of 156 (127 MS and 29 controls) snap frozen brain tissue blocks obtained at autopsies from 54 MS patients and 26 controls. MS patients and controls were similarly matched for age and sex. Samples were from frontal, parietal or temporal regions, with cortical GM or underlying WM. For detailed donor information see Extended Data File 1. As some brain banks only collect disease samples rather than controls, we inevitably have some differences in sample source with case-control status and different collection practices also impact age and PMI for the GM samples (Extended Data Table 1). However, we have had to be pragmatic with these precious resources. For example, analyses of these variables in our differential expression and cell composition analyses found very few genes or cell types associated with age. The details of the relevant genes are included in Extended Data File 4. We have added a statement to this effect in the limitations section.

#### Brain tissue characterization

Snap frozen tissue blocks from donors with GM lesions were provided by UKTB to Roche. Subpial GM lesions were determined using MBP and PLP staining by neuropathologists at Roche and confirmed by independent experts (Anna Williams, Roberta Magliozzi). Pathological staging of WM lesions from EBB and ABB donor samples was done at the respective brain banks. In the WM, de- and remyelinated lesions were identified by Luxol Fast Blue (LFB) staining and demyelinated lesions were grouped into active, chronic active and chronic inactive lesions with Oil red O staining to determine microglial activity^57^. Remyelinated lesions were defined as showing light staining on LFB, and presence of only a few non-activated microglia/macrophages which did not contain ingested MOG, PLP or MBP. Brain tissue specimens from the respective WM regions were shipped on dry ice to Roche and directly processed.

#### Nuclei isolation and single nuclei RNA sequencing

Nuclei were isolated from fresh-frozen 10μm sections, using Nuclei Pure Prep Nuclei Isolation Kit (Sigma Aldrich) with the following modifications. The regions of interest were macro-dissected with a scalpel blade, lysed in Nuclei Pure Lysis Solution with 0.1% Triton X, 1mM DTT and 0.4U/ul SUPERase-In™ RNase Inhibitor (ThermoFisher Scientific) freshly added before use, and homogenized with the help first of a 23G and then of a 29G syringe. Cold 1.8M Sucrose Cushion Solution, prepared immediately before use with the addition of 1mM DTT and 0.4U/ul RNase Inhibitor, was added to the suspensions before they were filtered through a 30μm strainer. The lysates were then carefully layered on top of 1.8M Sucrose Cushion Solution. Samples were centrifuged for 45min at 16000xg at 4°C. Pellets were re-suspended in Nuclei Storage Buffer with 0.4U/ul RNase Inhibitor, transferred in new Eppendorf tubes and centrifuged for 5min at 500xg at 4°C. Pellets were again re-suspended in Nuclei Storage Buffer with 0.4U/ul RNase Inhibitor, and centrifuged for 5 minutes at 500xg at 4°C. Finally, purified nuclei were re-suspended in Nuclei Storage Buffer with 0.4U/ul RNase Inhibitor, stained with trypan blue and counted using an automated cell counter (Countess II, Life technologies). A total of 12,000 estimated nuclei from each sample was loaded on the 10x Single Cell Next GEM G Chip. cDNA libraries have been prepared using the Chromium Single Cell 3’ Library and Gel Bead v3.3 kit according to the manufacturer’s instructions. cDNA libraries were sequenced using Illumina NovaSeq 6000 System and NovaSeq 6000 S2 Reagent Kit v1.5 (100 cycles), aiming at a sequencing depth of minimum 30K reads/nucleus.

### Sample swap checks via genotyping

We genotyped all samples included in this study using the GSAv3 illumina CHIP. Genotypes were imputed using the Haplotype Reference Consortium (HRC) reference panel (version r1.1)^58^ and lifted over to GRCh38. Genotype processing and quality control was performed using Plink v1.9^59^. SNPs with imputation score <0.4 or with missingness greater than 5% were excluded. We used MBV^60^ to identify sample swaps. Briefly, MBV takes as input a VCF file containing the genotype data of the samples, as well as bam files containing the mapped single nuclei sequencing reads. MBV then reports the proportion of heterozygous and homozygous genotypes (for each individual in the VCF file) for which both alleles are captured by the sequencing reads in all bam files. Correct samples can then be identified as they should have a high proportion of concordant heterozygous and homozygous sites between the genotype data and the mapped sequencing reads. We identified and corrected 23 sample swaps; three further samples were excluded because they could not be matched to a genotype.

### snRNAseq data processing and quality control

All samples were processed with CellRanger (v3.1.0), using the GRCh38 reference human genome and the ensembl Homo_sapiens GRCh38.96 reference annotation (modified to count intronic reads, and including genes with gene biotype protein_coding, lincRNA, antisense, and IG_* and TR_* genes). Ambient RNA contamination was removed via processing all samples with CellBender (v0.2.2), using the *raw_feature_bc_matrix.h5* output file from CellRanger as input, with *--total-droplets-included* set to 25000, *--expected-cells* set to the “Estimated Number of Cells” given in the *CellRanger metrics_summary.csv* output file, and other parameters set to defaults. Barcodes called as cells by CellBender, with the corresponding cleaned count matrices, were used for gene expression quantifications. We used *velocyto* (v0.17.17) on the *CellRanger* output to quantify intronic and exonic reads.

We identified doublets using *scDblFinder* (v1.12.0), applied to each sample separately (*multiSampleMode = “split”*), with all other parameters default^61^. We used the score threshold determined from the data by *scDblFinder*.

After removing doublets, we did quality control, primarily on the basis of percentage of exonic reads. We first removed nuclei with insufficient data to be worth including, requiring nuclei to have a minimum of 300 expressed genes, 500 UMI counts, and a maximum of 50% mitochondrial reads. We also excluded samples where an excessive proportion of input barcodes were called as cells by *CellBender*. Specifically, for each sample we calculated the proportion of droplets given >50% probability of being a cell by *CellBender*, applied the logit transform, calculated the median, and excluded any samples whose proportion was more than 3 MADs distant from the median. The result of this was to exclude 7 GM samples with >95% droplets called as cells, one GM sample with <3% droplets called as cells, and no WM samples. We then excluded any nuclei with greater than 75% exonic reads, or greater than 20% mitochondrial reads. After applying these filters, we then excluded any samples with <500 nuclei remaining. This resulted in 64 GM and 92 WM samples passing QC, comprising 632k nuclei across 156 samples.

### Data integration and clustering

#### Data integration

Data integration was done with *Harmony* (v0.1.1), as implemented within the Seurat package. Mitochondrial genes (those starting with MT-) were excluded. The counts data were loaded into a Seurat object, then we applied the functions NormalizeData, FindVariableFeatures, ScaleData, RunPCA (with n_dims = 50), followed by RunHarmony; with the exception of n_dims for RunPCA, all parameters were set to default. To identify clusters at the broad celltype level, we ran FindNeighbours then FindClusters, with resolution set to 1.

Broad cell types were assigned to each cluster on the basis of known marker genes: *PLP1, MAG, MOG, OPALIN* (Oligodendrocytes), *PDGFRA, BCAN* (OPCs / COPs), *FGFR3, GFAP, SLC14A1, AQP4* (Astrocytes), *P2RY12, SPP1, CSF1R, IRF8* (Microglia), *SLC17A7, FEZF2, RORB* (Excitatory neurons), *GAD1, ADARB2, LHX6* (Inhibitory neurons), *CLDN5, FLT1, EPAS1* (Endothelial cells), *EPS8, LAMA2* (Pericytes), *IGHG1* (B cells) and *IL7R* (T cells). The log normalised expression of each gene was calculated for each cluster, and these values scaled to 0 to 1 over all clusters. For each cluster, the broad cell type with the highest scaled expression averaged over the known marker genes was selected as the label.

#### Subclustering

Integration with Harmony identified cell types annotated with broad cell type labels. To identify subclusters corresponding to cell states, we repeated the integration process within the broad cell types. It is not advisable to integrate samples with very small numbers of cells. To avoid this, we first grouped the broad cell types together into the following six combinations of cell types, with the number of PCs used given in brackets: OPCs + COPS and oligodendrocytes (30 PCs); astrocytes (20 PCs); microglia (20 PCs); excitatory neurons (30 PCs); inhibitory neurons (30 PCs); and endothelial cells, pericytes, T cells and B cells (20 PCs).

For each of these broad cell type groupings, even after grouping together, some samples had very low numbers of cells. We therefore excluded any samples with fewer than 100 cells before performing integration with Harmony. Harmony was performed as above, with resolution 0.2 for all broad celltype groups, except OPCs + COPs and oligodendrocytes where we used resolution 0.5. Any clusters with fewer than 200 cells in total were excluded.

To label the cells in the samples with fewer than 100 cells, we trained a classifier (XGBoost^62^) on up to 1000 randomly selected cells from each cluster, using a second set of an equal number of randomly selected cells as a validation set. This classifier was applied to all unlabelled cells, and all cells that the classifier labelled with at least 50% probability were retained.

This resulted in 68 distinct batch-corrected fine cell type clusters, comprising: 11 of oligodendroglia; 7 of astrocyte; 7 of microglia / perivascular macrophages; 14 of cortical excitatory neurons (across layers 2-6); 12 of inhibitory neurons; 7 blood vessel-related cells (including 4 endothelial cell and 1 pericyte clusters); B cell and T cell subpopulations; and 9 clusters with mixed lineages. These mixed clusters could potentially be doublets that had not been identified by *scDblFinder*, and were therefore excluded from further analysis; this resulted in 59 QC-passed clusters.

#### Independent data processing, integration and clustering

We additionally performed an entirely independent processing pipeline (distinct methods for: handling ambient RNA contamination; QC; integration; and clustering). We found high concordance between the clusters identified by both methods (Extended Data Fig. 2a, Extended Data File 2).

### Marker gene identification and cell type annotation

To identify marker genes within each of the 6 broad cell type groupings, we used *edgeR* applied to pseudobulk counts of each subcluster in each sample. This avoids the inflated FDR values due to pseudoreplication that are common to methods such as FindMarkers in Seurat^12^. To do this, we first constructed a matrix of pseudobulk values, where each row corresponds to a gene, and each column corresponds to the sum of all the cells of one cluster in one sample. We ran *edgeR* using *calcNormFactors* with the method “TMMwsp” applied to library sizes across all clusters simultaneously (the default here is “TMM”, however “TMMwsp” is designed to better take zeros into account, and is therefore more appropriate when some samples may have small library sizes). The cluster variable was used to construct the design matrix for estimating dispersions via *estimateDisp*. We then ran *glmQLFit* and *glmTreat* on each cluster, with the formula *∼ is_cluster*, where *is_cluster = cluster == sel_cl*, to identify genes that are differentially expressed in that cluster relative to all other clusters. We defined marker genes as genes differentially expressed in a cluster, not labelled as either lincRNA, pseudogene or antisense, with positive logFC, and >= 1 CPM mean expression in that cluster. We then ran FGSEA on the marker genes identified for each cluster, using logFC as the ranking variable.

### Differential abundance of cell types in MS lesions and control samples - *ANCOM-BC*

We first checked for samples with sample sizes that were much smaller than for other samples, to exclude samples where abundances might be very noisy. We excluded samples with log sample sizes 2*MAD (median absolute deviation) less than the median log sample size, separately for WM and GM; this excluded zero WM samples and two GM samples. We also checked for samples with unusual proportions of neuronal cells relative to other WM or GM samples. White matter samples with neuronal proportion at least 2*MAD (median absolute deviation) higher than the median neuronal proportion for all WM samples were excluded; grey matter samples with at least 2*MAD neuronal proportion fewer than the GM median neuronal proportion were excluded. This excluded 18 out of 94 WM samples and 1 out of 71 GM samples.

To test whether abundances of fine cell types across samples were affected by lesion type and donor ID (**Fig. 2i,j**), we used likelihood ratio tests applied to models including lesion type and donor id. Briefly, we fit a series of nested models for each fine cell type: full (*counts ∼ lesion_type + sex + age_scale + pmi_cat + (1 | donor_id)*), fixed (*counts ∼ lesion_type + sex + age_scale + pmi_cat*), covariates only (*counts ∼ sex + age_scale + pmi_cat*) and null (*counts ∼ 1*). We used the R package *glmmTMB* (v1.1.2.2)^27^ to fit a negative binomial distribution to the raw counts for each cell type (see Supplementary Note for explanation of use of raw counts rather than proportions). We used the function *anova* to perform likelihood ratio tests of the following nested sequence of models: full; fixed; covariates only; null. This gives a p-value indicating whether the more complex model improved the fit more than would be expected by chance. We adjusted the p-values across these tests using the Benjamini-Hochberg procedure, across all cell types and models together.

To test for differential abundance in fine cell type due to lesion type, we used ANCOM-BC version 1.3.2^63^. The likelihood ratio test analysis above indicated that donor ID needed to be taken into account, however this version of ANCOM-BC does not allow random effects. To factor out donor effects, we therefore did a bootstrapped analysis of abundance: each bootstrap took one random sample per donor, and ran ANCOM-BC on each bootstrapped sample (e.g. in WM, there were 76 samples across 42 donors, so each bootstrap comprised 42 samples). We summarised the results of 20k bootstraps by taking the median, 80% and 95% confidence intervals of the inferred coefficients for each fine cell type (20k samples is sufficient to properly estimate tail probabilities for 95% CIs; cf ^64^).

To test differential abundance in WM samples, we additionally excluded all neuronal cell types, as these should not be present in WM. We fit ANCOM-BC with the formula *∼ lesion_type + sex + age_scale + pmi_cat*, where *age_scale* is patient age, scaled to have SD = 0.5 across all patients in the dataset^56^, and *pmi_cat* is post-mortem interval, split into three categories (under 1 hour, between 1 and 12 hours, and more than 12 hours).

To test differential abundance in GM samples, we first fit the data with a similar formula: *∼ lesion_type + sex + age_scale + pmi_cat2* (here, *lesion_type* includes ctrGM, NAGM and GML; *pmi_cat2* has only two categories to reflect the values observed in GM, between 1 and 12 hours, and more than 12 hours).

Using this formula this produced results that conflicted with known biology, identifying multiple neurons as increasing in abundance in GM lesions relative to GM controls. Analysing differences between neuronal proportions between ctrGM, NAGM and GML, we observed that GML samples were enriched in L1/L2/L3 neurons, and those from NAGM samples were enriched in L5/L6 neurons (ctrGM samples had intermediate proportions) (Extended Data Fig. 10a). This indicated that GML samples were taken from more superficial cortical layers in the brain, and the matched NAGM also contained deeper layers (although the experimenters had made efforts to take all samples from the same depth).

To identify layer effects for each sample, we calculated principal components (PCs) reflecting neuronal layer distributions in normal tissue. We applied PCA to the centred log ratios of the neuronal cell types in the control GM samples. We then identified principal components that could be relevant to layers (by filtering on both the absolute Spearman rank correlation between the PC loading and the layer numbers of neurons known to be layer-specific, thresholding at minimum 0.2 correlation) and which explained at least 1% of variance. This identified seven PCs that could contain layer information (Extended Data Fig. 10b). This analysis was performed in control GM samples only; we then calculated CLRs for all samples, and projected them into the selected PCs, using the loadings derived from the control samples.

We then repeated the bootstrapped ANCOM-BC analysis, including layer PCs as covariates to factor out layer effects. We used the formula *∼ lesion_type + sex + age_scale + pmi_cat2 + layerPC1 + … + layerPCn*, i.e. we repeated the analysis using the first *n* layer PCs identified above, including from 1 up to 7 PCs. We found that including the first 3 layer PCs gave results that fitted well with expected biology, i.e. that almost no neuronal types were found to increase in abundance in either NAGM or GML relative to control GM, and PVALB+ neurons decreased in abundance (Extended Data Fig. 10c). We included 3 layer PCs, however there is little difference in the results for including between 3 and 7 layer PCs.

### Differential abundance of cell types in MS lesions and control samples - *miloR*

*miloR* is an R package that identifies neighbourhoods within a nearest neighbours graph that are enriched or depleted in cells from a particular condition^26^. We ran Milo on WM and GM samples separately, using a graph constructed on Harmony-corrected principal components.

Briefly, we restricted to relevant celltypes for each tissue (WM: OPCs + COPs, Oligodendrocytes, Astrocytes, Microglia, T cells, B cells; GM: OPCs + COPs, Oligodendrocytes, Astrocytes, Microglia, Excitatory neurons, Inhibitory neurons, T cells, B cells), and selected 2000 highly variable genes evenly across each major cell type (this ensures that the HVGs are not dominated by the celltypes with largest numbers). We then ran Harmony on the selected cells and HVGs using the default parameters.

To account for multiple samples per donor, we did a bootstrapped analysis of abundance, as with ANCOM-BC: each bootstrap took one random sample per donor, and ran *miloR* on each bootstrapped sample. (The bootstrapping was only of the counts of cells, and not of the graph construction.) We performed 2000 bootstrap replicates for each model tested.

To identify the neighbourhoods associated with lesion types, we applied the bootstrapped *miloR* with the following formulae: WM, *∼ lesion_type + sex + age_scale + pmi_cat*; GM, *∼ lesion_type + sex + age_scale + pmi_cat2 + ctrl_PC01 + ctrl_PC03 + ctrl_PC04*. The 3 layer PCs here are to correct for layer effects in the GM tissue, as described in the ANCOM-BC analysis. To identify the neighbourhoods associated with factors, we applied the bootstrapped *MiloR* with the following formulae: WM, *∼ WM_F1 + WM_F2 + WM_F3 + WM_F4 + WM_F5 + sex + age_scale + pmi_cat*; GM, *∼ GM_F1 + GM_F2 + GM_F3 + GM_F4 + GM_F5 + sex + age_scale + pmi_cat2 + ctrl_PC01 + ctrl_PC03 + ctrl_PC04*.

To plot the bootstrap results, we showed the median bootstrap value per neighbourhood. We reported the coefficient for a neighbourhood as being significantly different from zero if the 90% bootstrap interval of the log2FC did not overlap with the interval *[-log2(1.2), log2(1.2)]* (this is the same principle as in the *TREAT* method in *edgeR* ^65^). We used the function *annotateNhoods* in *miloR* to give fine celltype labels to each neighbourhood.

### Differential expression analysis using generalised linear mixed models

To identify genes differentially expressed in MS WM and MS GM samples compared to respective control samples per cell type, we did differential expression analysis on pseudo-bulk data, i.e. analysis at the level of the transcript totals across all cells of a given type in each sample. Pseudobulk approaches are known to offer a good compromise between sensitivity and run time constraints^66,67^ (see Supplementary Note for the details of our analysis using different pseudobulk approaches and identification of strong patient effects). Inspection of gene expression at the donor level indicated that our model would need to include donor effects.

We therefore used glmmTMB^27^ with a negative binomial model, and *donor_id* as a random intercept. To filter out samples with low library sizes or numbers of cells, genes with low counts, and estimate library sizes, we followed the approach set out in *muscat*^66^. Briefly, this comprises: removing samples with fewer than 10 cells; removing pseudobulk samples whose log library size is less than 3 MADs less than the median; removing genes with low expression using the function *filterByExpr* in *edgeR*; calculating TMM-normalized library sizes with the function *calcNormFactors* in *edgeR*^68^.

The formula for WM was *counts ∼ lesion_type + sex + age_scale + pmi_cat + (1 | donor_id)*, where *pmi_cat* is post-mortem interval, split into three categories (under 1 hour, between 1 and 12 hours, and more than 12 hours, and *age_scale* is patient age, normalised to have *SD = 0.5*^56^.

In the GM analysis, we accounted for layer effects by including 3 layer PCs as described in the ANCOM-BC analysis. The formula for GM was therefore *lesion_type + sex + age_scale + pmi_cat2 + ctrl_PC01 + ctrl_PC02 + ctrl_PC03 + ctrl_PC04 + (1 | donor_id);* to reflect values observed in GM samples, *pmi_cat2* has only two categories (between 1 and 12 hours, and more than 12 hours). We included an offset of *log(lib.size) - log(1e6)*, so that the reported coefficients correspond to log counts per million (logCPM). Genes with absolute log2 fold change in expression of at least *log2(1.5)* and an FDR-corrected *P < 0.05* were selected as differentially expressed. FDRs were calculated at the level of combination of cell type and model coefficient.

To quantify the extent of donor effects, for each gene we also used *glmmTMB* to fit three simpler models: with lesion type plus covariates (i.e. fixed effects only) (*counts ∼ lesion_type + sex + age_scale + pmi_cat*), with covariates only (*counts ∼ sex + age_scale + pmi_cat*) and a null model (*counts ∼ 1*). We then used the anova function to perform likelihood ratio tests for this nested sequence of models; we applied a Benjamini-Hochberg correction across all genes and LRTs, separately within each cell type.

### Gene set enrichment analysis of differentially expressed genes

FGSEA^69^ was used to perform statistical enrichment tests of differentially expressed genes in each cell type (broad and fine) from each comparison in WM and GM samples. All genes expressed in a given cluster were used as a background list, and GO-term analysis for enriched biological processes and hallmark genes from MSigDB^70^ was performed. Z-score was used as the ranking variable in FGSEA (calculated from the unadjusted p-value and the sign of the log fold change). FDR correction was calculated within each combination of cell type, model coefficient and pathway collection. Processes with an FDR-corrected *P* < 0.1 were considered and their normalised enrichment scores (NES) plotted as a dotplot using ggplot2^71^ R-based libraries.

### Assessment of cluster connectivity with *PAGA*

To characterise connectivity between clusters, we used PAGA^17^ as implemented in scanpy version 1.8.2^72^. As input, we used the nearest neighbour graph constructed by conos, restricted to just cells with oligodendrocyte or OPC / COP labels. We ran PAGA clustering and layout embedding using fine cell types as the group variable, and otherwise used defaults.

### Patient stratification using *MOFA+*

We used MOFA+ to identify factors explaining the variability across the samples (implemented in the R package MOFA2)^29^. MOFA+ was developed for data with multiple modalities measured from the same samples. In this study, we took the different *cell types* to be the different modalities. This allows us to identify responses that are coherent across samples, across multiple cell types simultaneously, and which may have cell type-specific responses. MOFA+ does this by finding factors that seek to explain the variability in the input data (intuitively similar to PCA) across samples, and which correspond to coordinated tissue-level responses, even though the genes identified for each cell type are distinct. For example, a factor corresponding to remyelination might involve myelination genes in oligodendrocytes, debris-clearance genes in microglia, and metabolic support genes in astrocytes. These factors can then suggest possible ways to stratify patients.

We first identified genes with relevant variation for each cell type, based on the negative binomial models fitted to each gene in each cell type. We identified genes with either an MS effect, or a donor effect (or both). Genes with an *MS effect* were defined as those where at least one lesion type had both an FDR < 1% and an absolute log2 fold change of 1 for WM, or log2(1.5) for GM (we observed lower effect sizes in GM, and therefore used a more relaxed threshold). Genes with a *donor effect* were those where the likelihood ratio test of including the donor effect had an FDR < 1%, and the standard deviation of the donor random intercepts was at least *log(2)* for WM, and *log(1.5)* for GM. These thresholds are arbitrary but we have found the factors identified by MOFA+ to be robust to variations on these thresholds. In WM, this resulted in the selection of: 231 genes for OPCs + COPs, 821 for oligodendrocytes, 1192 for astrocytes, 1023 for microglia, 225 for endothelial cells and pericytes. In GM, we selected: 339 genes for OPCs + COPs, 1039 for oligodendrocytes, 1239 for astrocytes, 412 for microglia, 1159 for excitatory neurons, 924 for inhibitory neurons, 852 for endothelial cells and pericytes.

We then calculated normalised expression for the selected genes in each cell type as input to MOFA+. We first excluded any samples with fewer than 10 cells observed for that cell type (this means that there may be missing data for some cell types in some samples). We also excluded any samples with log library sizes 3*MAD (median absolute deviation) less than the median log library size for that cell type. For the remaining samples, we calculated the *log(CPM + 1)* of the pseudobulk values, calculating CPMs with library sizes via the effectiveLibSizes function in edgeR^68^. To remove any possible layer effects in GM samples, we fit a linear model using the first four layer PCs as covariates, i.e. *logCPM ∼ layerPC1 + layerPC2 + layerPC3 + layerPC4*, and used the residuals of this model as values for input to MOFA+. To ensure each gene contributed equally to the model, we then z-scored all resulting values within each combination of gene and cell type.

For each of WM and GM, we then fit MOFA+ to this data, using 5 factors. As we are interested in an unbiased characterisation of the heterogeneity of the data, we did not use the group variable in MOFA+; otherwise we used the default parameters. In both WM and GM, we found 5 factors which explained at least 5% of variance for some cell type.

As described above, we fit MOFA+ only to a relevant subset of genes for each celltype. To identify associations between the identified factors and all genes, we then used *edgeR* fit to the counts of each gene, with the formula ∼ *WM_F1 + WM_F2 + WM_F3 + WM_F4 + WM_F5 + sex + age_scale + pmi_cat* (and similarly for GM).

To estimate the variance explained by the MOFA factors across all genes, we first obtained variance-stabilised log transforms of the data with the *vst* function in the R package *DESeq2*^73^; notably, this shrinks the high sampling variability of genes with low mean counts. We then fit a linear model to the log-transformed values of each gene with the formula above, calculated *anova*, and recorded the sum of squares for each factor. We restricted to genes with *vst*-transformed variance of at least 0.5, and calculated variance explained as the total sum of squares for each factor, divided by the total variance across all included genes.

To calculate geneset enrichment of the genes in MOFA+ factors, we ranked the genes for a given cell type in descending order of the signed log *edgeR* p-value, and used the function *fgseaMultilevel* from the FGSEA package^69^, using a minimum set size of 5 genes and otherwise the default parameters.

### Validation of MOFA+ factors with *scITD*

scITD is a computational method designed to extract multicellular gene expression programs that vary across donors or samples^33^. We ran scITD on WM pseudobulk data from the same set of cell types used as input to MOFA (i.e. OPCs + COPs, oligodendrocytes, astrocytes, microglia, endothelial cells and pericytes). We used the parameters set out in the method vignette (http://pklab.med.harvard.edu/jonathan/), resulting in inclusion of 5098 genes across all cell types.

### Immunohistochemistry and analysis

FFPE sections (4 μm) were deparaffinized in decreasing concentrations of ethanol, and antigen retrieval was performed in antigen unmasking solution (Vector Laboratories, H-3300) for 10 min in the microwave. Sections were incubated with autofluorescence eliminator reagent (Millipore, 2160) for 1 min and washed with TBS 0.001% Triton-X (wash buffer). Endogenous peroxidases were quenched with 3% H_2_O_2_ for 15 min at room temperature (RT), washed in wash buffer and blocked for 30 min at room temperature with PBS 0.5% Triton-X (TBS-T), 10% HIHS (blocking buffer). Primary antibody incubation was performed overnight at 4 °C in blocking buffer. Sections were washed and incubated for 2hrs at RT with HRP-labelled secondary antibodies. Fluorophore reaction was performed using OPAL 570 and OPAL 650 reaction kits for 10 min at RT (Akoya Biosciences, FP1488001KT and FP1496001KT respectively, 1:500). Sections were counterstained using Hoechst (Thermo Fisher, 62249; 1:1,000), washed and mounted.

The following primary antibodies were used: mouse anti-CNP (Atlas, AMAb91072, 1:1,000), rabbit anti-human GPR17 (Cayman Chemical, 10136, 1:100), MBP (Rat anti MBP aa82-87 BioRad 1:300), PLP (Anti-Myelin Proteolipid Protein Antibody, CT, clone PLPC1 MAB388 Sigma-Aldrich, 1:200), MHCII (Dako 1:50), MOG Z12 (inhouse clone 1:50^74^). The following secondary antibodies were used: Vector Laboratories, rabbit-HRP IgG (MP-7401, Vector laboratories), mouse-HRP IgG (MP-7402, Vector laboratories).

For quantifications of GPR17 cell numbers, sections were co-labelled with GPR17 and CNP which was used to define demyelinated lesions. Sections were scanned using a VectraPolaris slide scanner and processed using Qupath^75^ and Fiji^76^ imaging software. Within individual lesions, several regions of interest were selected randomly. These regions of interest were randomised using the Fiji randomization plugin and quantified completely blinded mixing samples from all conditions, regions and lesions.

### Statistical analysis

No statistical methods were used to predetermine sample sizes, but our sample size is eight times larger than those reported in previous snRNAseq MS publications (Jäkel et al.^7^, Schirmer et al.^8^, Absinta et al.^9^). Statistical analyses and graphical visualisations were performed using open-source R programming software^77^. See dedicated method sections for the details of the snRNA-seq bioinformatic analysis; differentially expressed genes were defined as genes significantly expressed (*P* adjusted for multiple comparisons < 0.05), and showing, on average, >1.5-fold difference between groups of nuclei in each cell type in every DEG comparison. Volcano plots were constructed by plotting the log2(fold change) of lesion type with smallest p-value for each gene in the *x* axis and by plotting standard deviation of random (donor) effects for each gene on the y-axis. Statistical analysis used two tailed parametric or non-parametric t-tests for two groups, and Fisher’s exact test and one-way analysis of variance with corresponding post hoc tests for multiple group comparisons. Data distributions are presented as barplots, dotplots (with individual data points) and heatmaps. Log CPM gene expression values in the dot plots and heat maps were averaged, mean-centred, and *z*-score-scaled. Dot size indicates the percentage of nuclei in the cluster in which the gene was detected; among the nuclei in which the gene was detected, the expression level was mean-centred and scaled. Graphical object in Fig. 1a was created with BioRender.com.

## Data availability

All raw snRNA-seq data (FASTQ files) have been deposited to EGA^78^, as dataset EGAD00001009169. The cleaned annotated counts matrices, sample metadata and cell type annotations for both cohorts, and R and R markdown scripts, are available on Zenodo: <u>DOI</u> <u>10.5281/zenodo.8338963</u>. An interactive web browser to analyse cell type-specific expression levels of genes and transcriptomic changes in MS versus control tissue is available at https://malhotralab.shinyapps.io/MS_broad/ (for broad cell types) and at https://malhotralab.shinyapps.io/MS_fine/ (for fine cell types).

## Code availability

The source code used to analyse the snRNA-seq data in the current study is available online at https://wmacnair.gitlab.io/MS_lesions_snRNAseq.

## Author contributions

DM designed the study; WM, CFC, AW, GCB and DM coordinated all the work and wrote the manuscript; WM designed the computational analysis, performed QC, clustering, differential gene expression, cell type composition, PAGA, MOFA+ and Milo analysis; DC developed and optimised the snRNAseq protocol; DC with assistance from SJ performed the nuclei extraction, library preparation and snRNAseq experiments for 173 samples; DC, PK and SJ contributed to the selection of cluster marker genes for validation; SJ and AW characterised WM samples from EBB; SJ performed and analysed the GPR17 validation experiments; SA, MM and EN characterised WM samples from ABB; NS and VO performed MBP stainings and characterization of GM lesions from UKTB with assistance from DC, AW and RM; LF, LC, SS and EU helped with procuring WM and GM tissue samples from ABB and UKTB, and interpretation of the results; JB and EA mapped raw sequencing reads and quantified spliced/unspliced reads; JB quantified ambient RNA, performed genotyping QC, identified sample swaps, performed MAGMA analysis and developed the ShinyApp; EA performed data integration using Seurat and Harmony; WM and MR developed the SampleQC method, and MR provided input on the DE analysis; All co-authors read, revised and approved the manuscript.

## Acknowledgements

We thank Catharine Fournier Aquino and her team at the Functional Genomics Centre Zurich (FGCZ) for help with illumina sequencing of 10X single nucleus RNA-seq sample libraries; Kelly Bales and Irene Knuesel for their support with initiating this project; Elyas Heidari and Pierre-Luc Germain at the University of Zurich, Switzerland for assistance with gene module and *scDblFinder* analysis; and members of Malhotra lab and Collin lab at Roche for fruitful discussions of the results. AW is funded by the MS Society UK, MRC, and the UKDRI (which is funded by the MRC, Alzheimer’s Society and Alzheimer’s Research UK). G.C.-B.’s research group-Swedish Research Council (2019-01360); the European Union (2020 Research and Innovation Program/European Research Council Consolidator Grant EPIScOPE, 681893), Swedish Brain Foundation (FO2017-0075; FO2018-0162), Ming Wai Lau Centre for Reparative Medicine, Knut and Alice Wallenberg Foundation (grants 2019-0107; 2019-0089), Swedish Society for Medical Research (SSMF, grant JUB2019), the Göran Gustafsson Foundation for Research in Natural Sciences and Medicine, and Karolinska Institutet. E.A. was funded by the European Union, Horizon 2020, Marie-Sklodowska Curie Actions, grant SOLO, number 794689.

