## Extended Data for "Single nuclei RNAseq stratifies multiple sclerosis patients into distinct white matter glial responses"

#### Extended Data Figures

**ED1a** Summary of numbers of cells, samples and donors excluded and retained by QC procedure, split by various metadata labels. Distributions of metadata labels split by control and MS samples: *age_cat* is age at death binned into categories; *yrs_w_ms* is years with MS, binned into categories, and NA for control samples; *pmi_cat* is post mortem interval, binned into categories; *sample_source* is the brain bank of origin; and *seq_pool* is the batch in which the samples were sequenced.

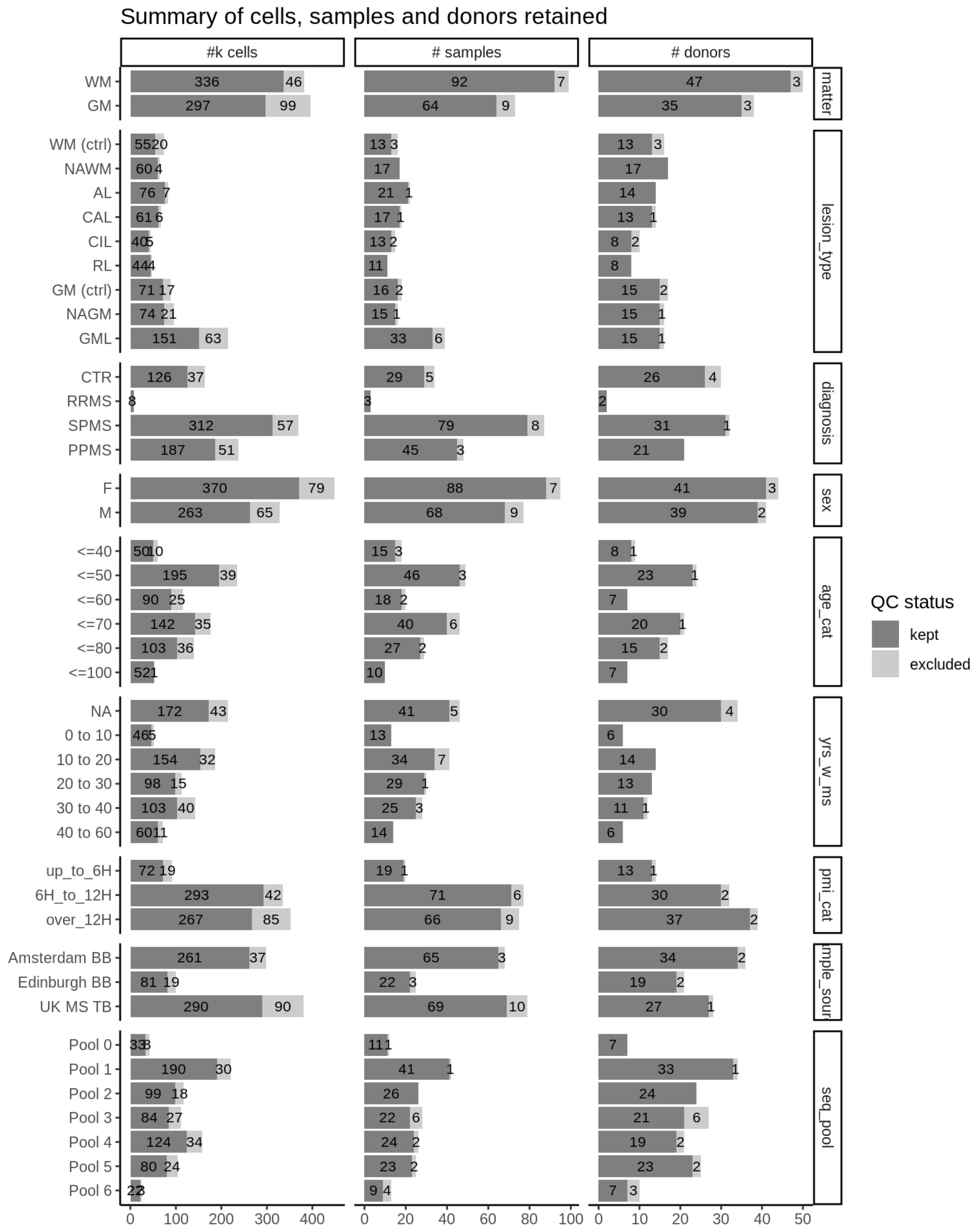

Summary of cells, samples and donors retained
#k cells
### samples
### donors
matter
lesion_type
diagnosis
sex
age_cat
yrs_w_ms
pmi_cat
seq_pool
QC status
kept
excluded

**ED1b** Numbers of WM and GM donors across metadata categories, split by control and MS samples. *pmi_minutes* is post mortem interval in minutes.

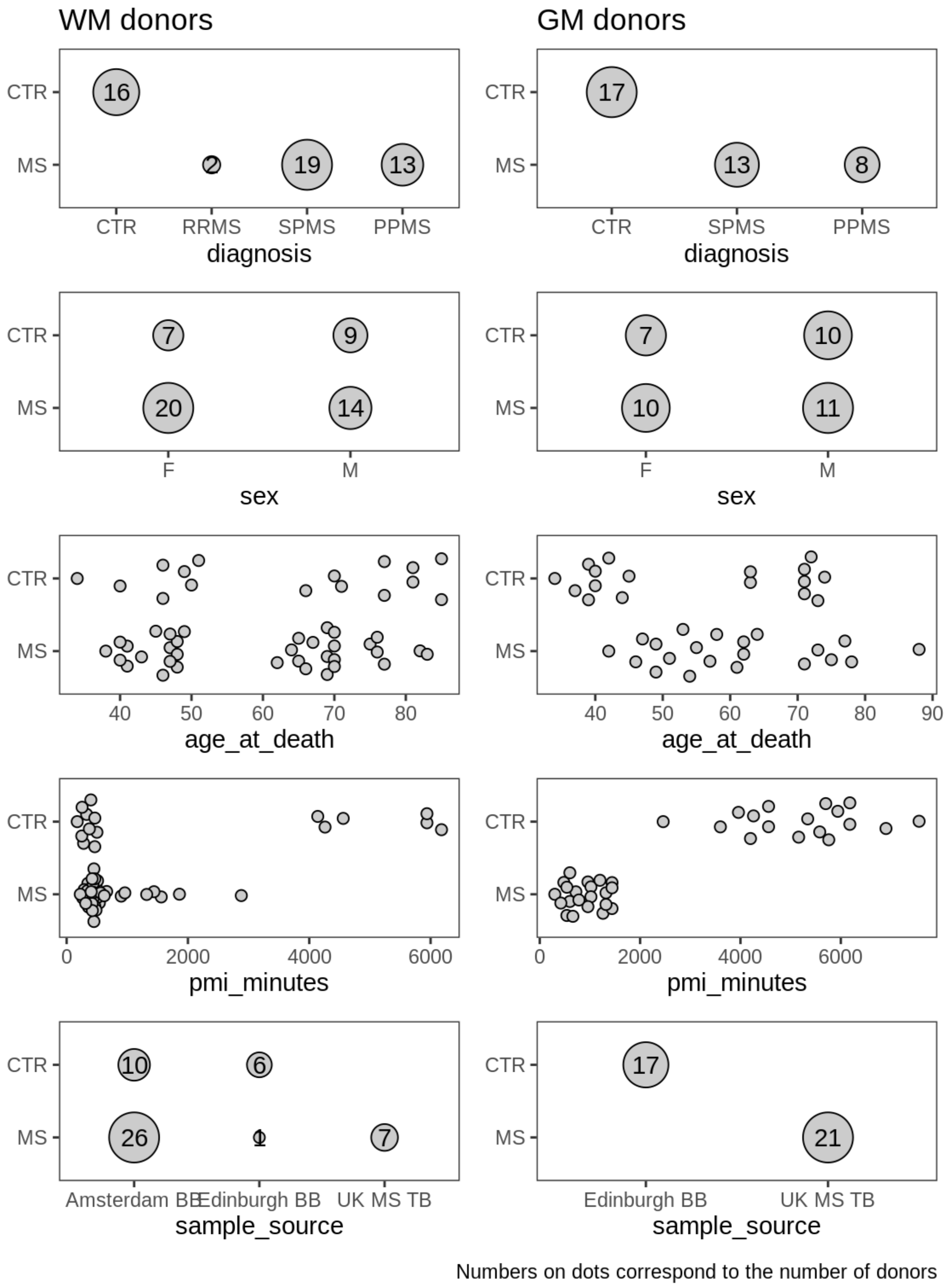

**ED1c** Summary of QC metrics of post-QC WM samples. *donor* is colour for donor ID, with grey values used for donors contributing only one sample; *mito pct* is the proportion of reads in the sample that are mitochondrial; *pct spliced* is the proportion of reads in the sample that are spliced (as opposed to spliced mRNA). Colours in heatmap are the z-scores for each QC metric column, with colours chosen so that red is good and blue is bad (e.g. low library size, or high mitochondrial read percentage).

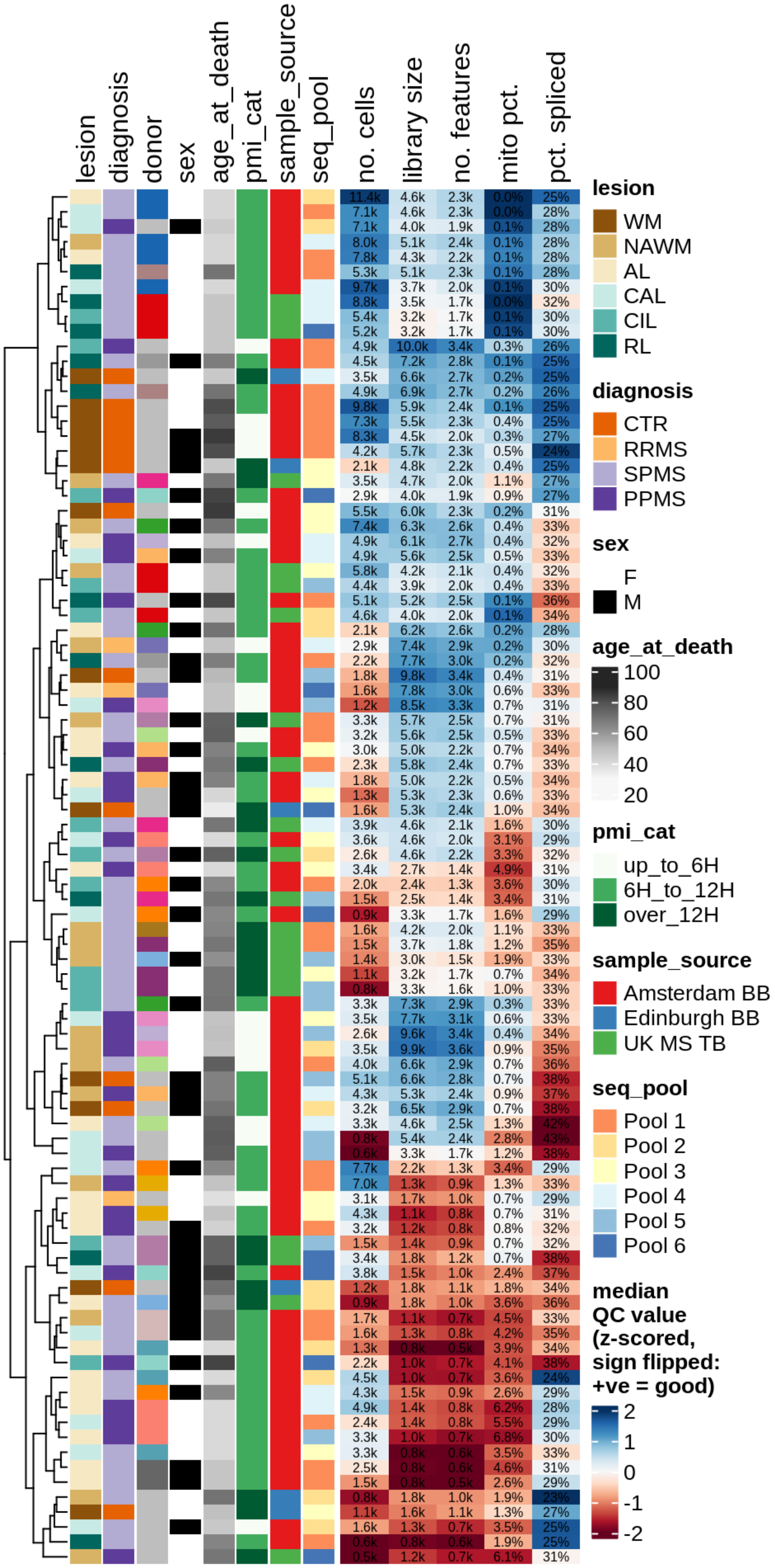

**ED1d** Summary of QC metrics of post-QC GM samples, as for **ED1c**.

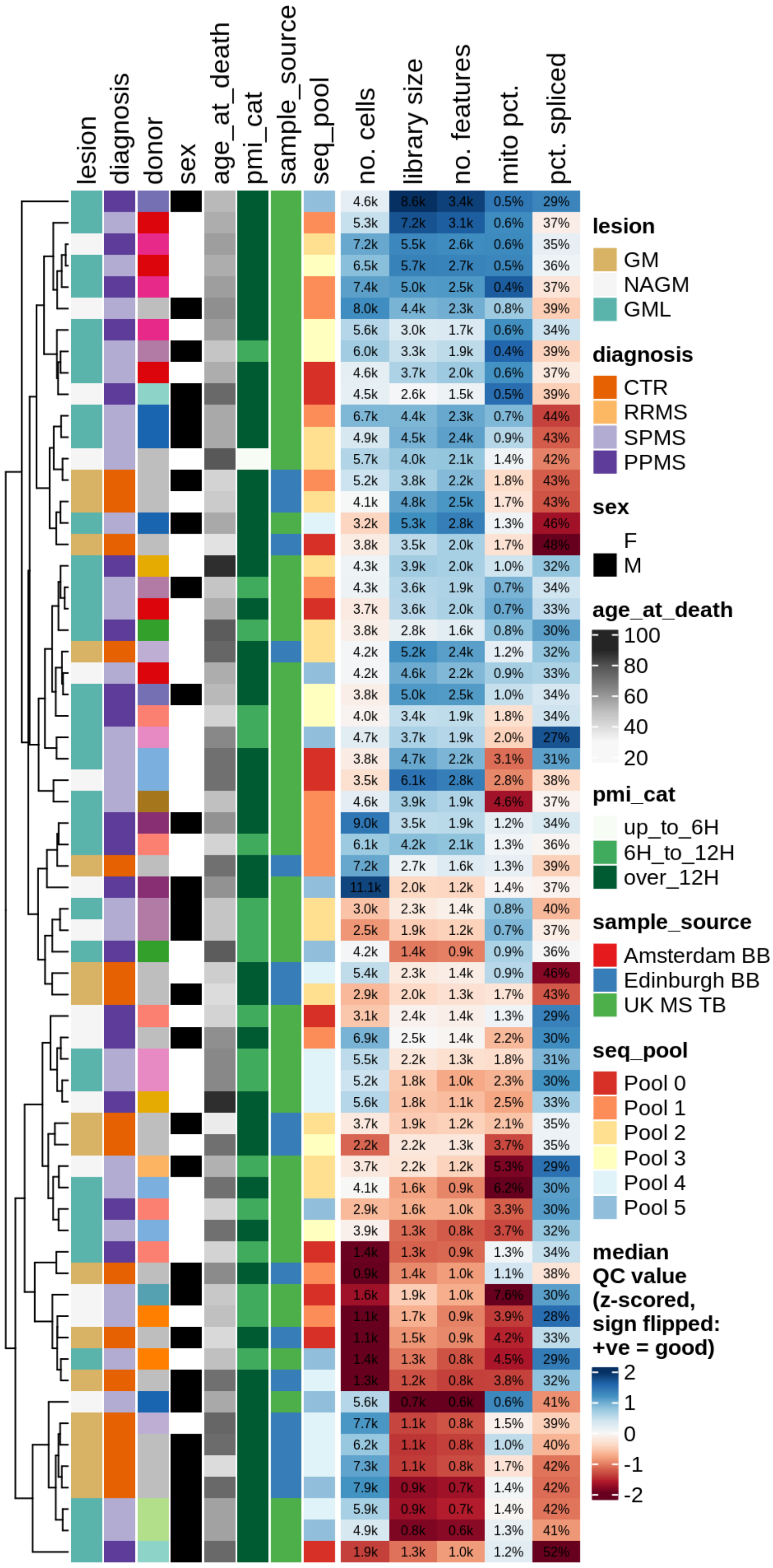

**ED2a** Comparison of fine cell type clusters with orthogonal data integration and clustering analysis presented in Extended Data File 2. Rows are the fine cell types used in this manuscript; columns are the fine cell types via the alternative processing approach. Columns sum to 1. The annotations ("used only", "alternative only") show how many cells were unique to this processing approach's QC filtering.

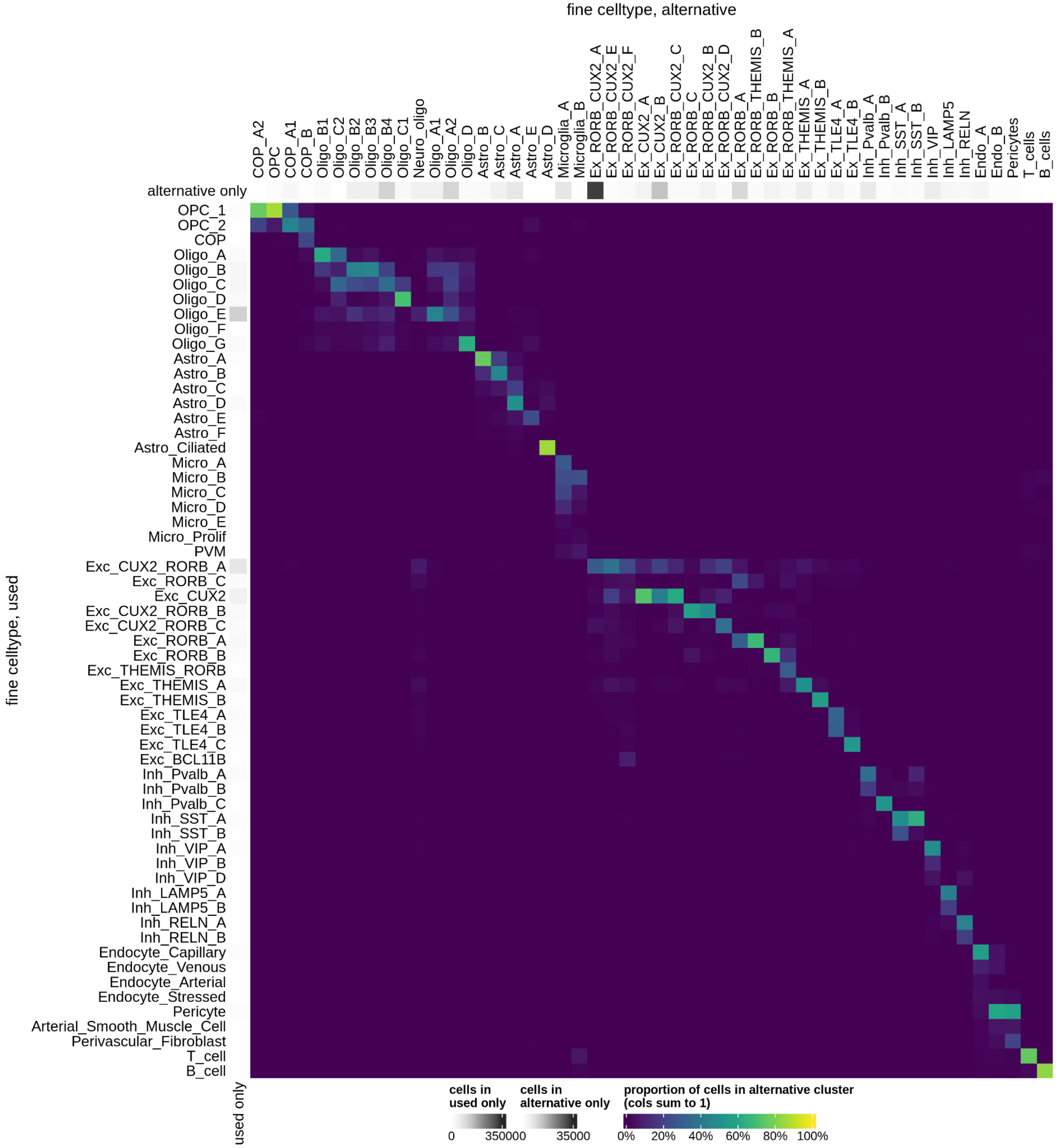

**ED2b** QC metric summaries for fine celltypes. Each point is a sample with >= 10 cells of that type, showing median QC metric value for those cells in that sample (with exception of number of cells).

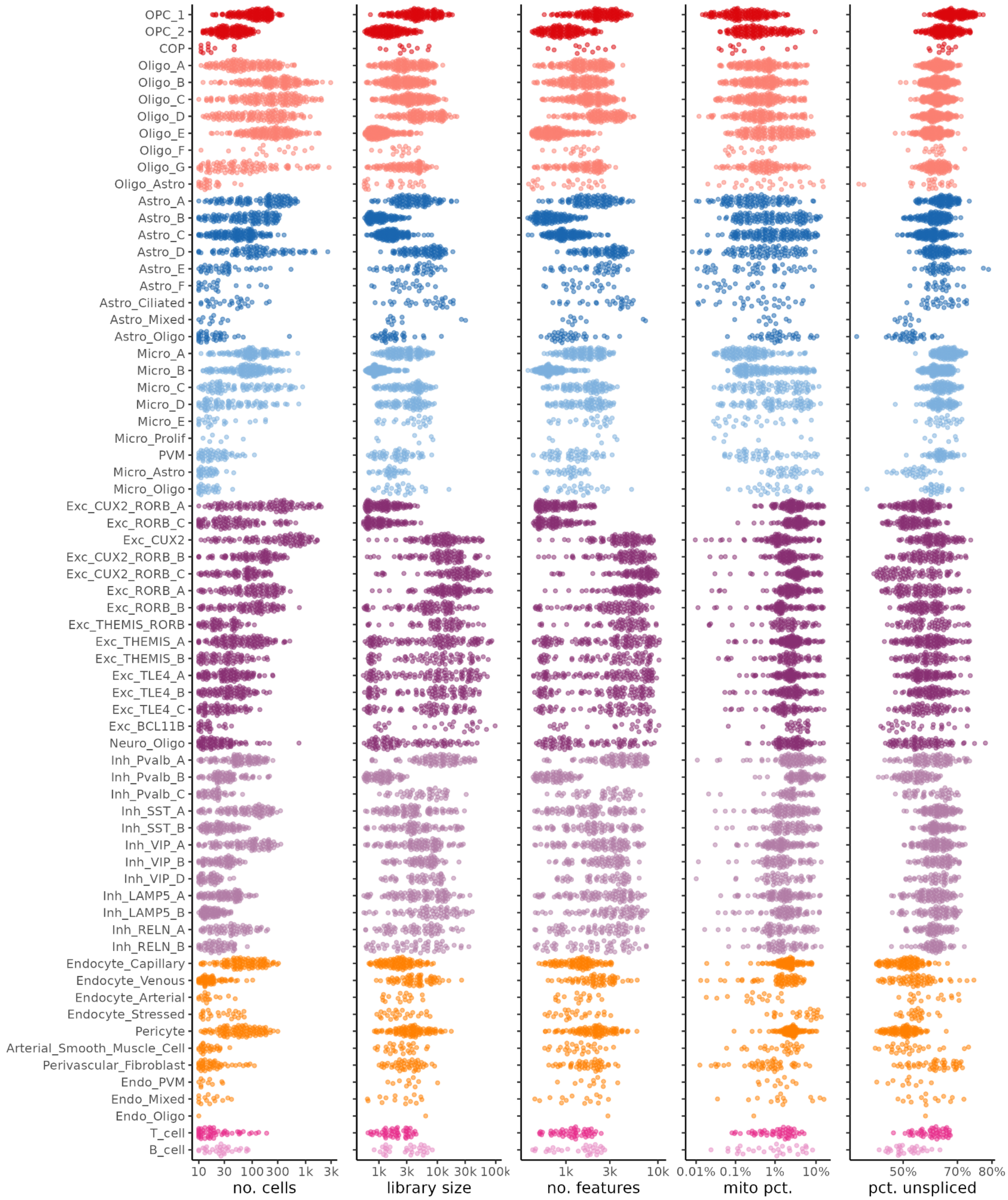

**ED2c** Analysis of cluster mixing across various metadata variables. Entropy (Shannon entropy) calculated by taking total number of cells across each value of metadata variable and converting this to proportions, then calculating the entropy of this proportion vector. For example, if a celltype has 10k cells in GM and 20k cells in WM, then the proportions are 0.33 and 0.67, giving an entropy of 0.92. Entropy reaches the highest possible value when cell counts are equally split across all metadata values.

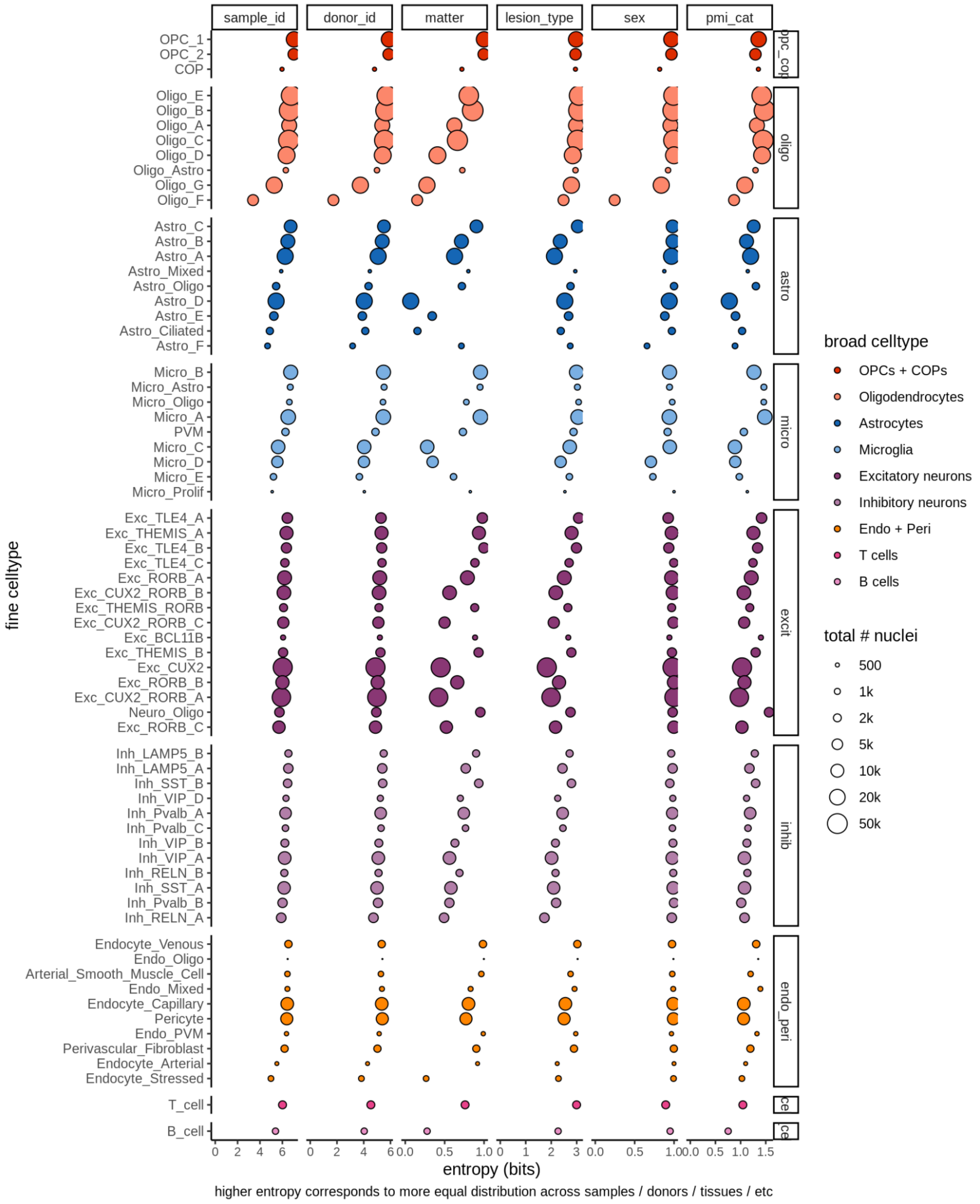

**ED2d** Expression of marker genes selected for broad and fine celltypes. CPM indicates counts per million, i.e. number of counts of gene normalised by total number of pseudobulk counts. Expression and proportions are calculated across all cells and samples.

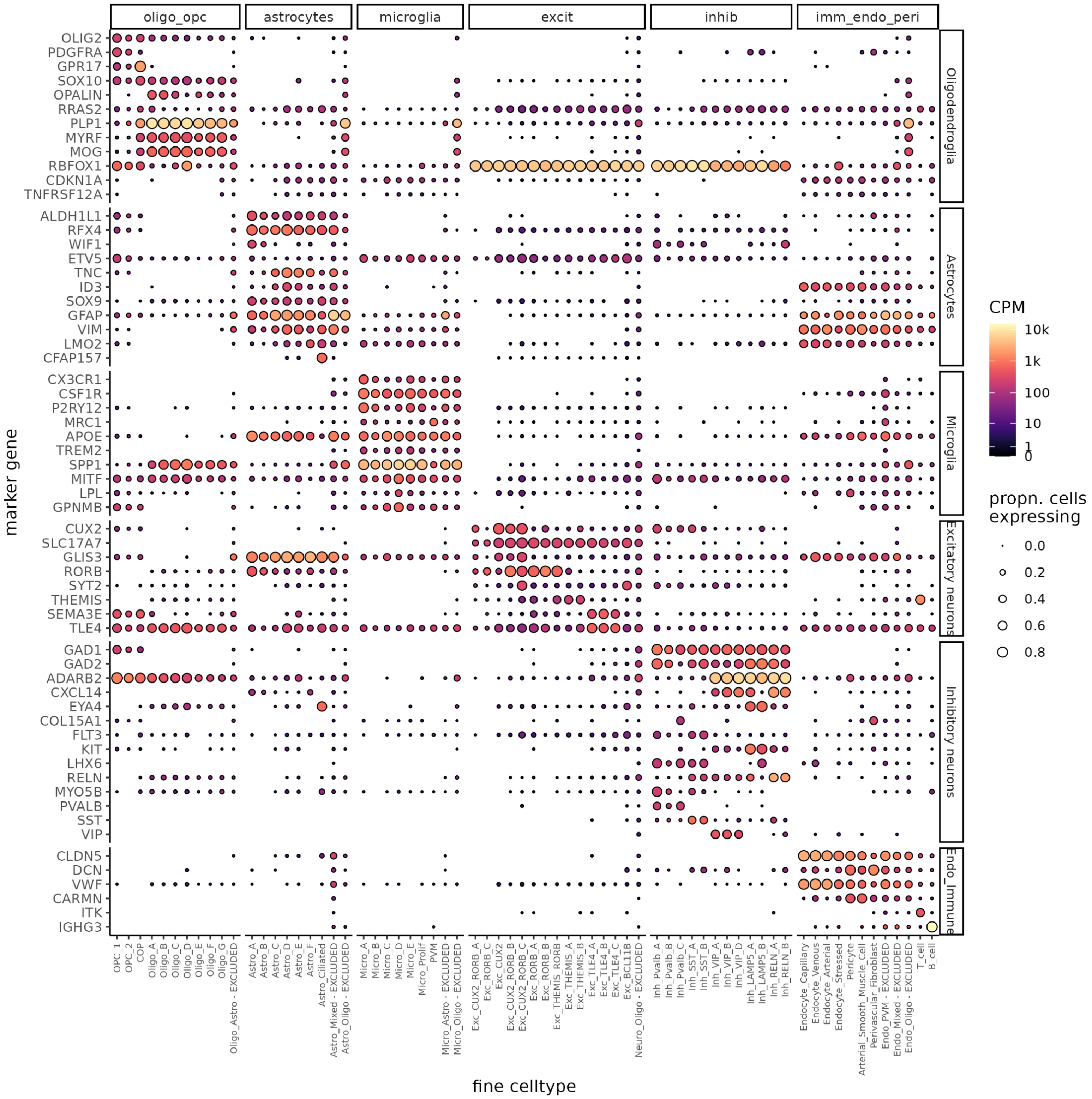

**ED2e** UMAP embedding (as in **Fig. 1b**) annotated with proportion of nuclei in the binned region of UMAP embedding coming from MS as opposed to control samples (left) and WM as opposed to GM samples (right). In both plots, white corresponds to the average proportion across all cells (i.e. 20% of nuclei are from MS samples, and 60% of nuclei are from GM samples). If we used a colour scale with white at 50%, then almost all of the left plot would be a deep brown, which would just indicate that we have more MS than control samples, and give no dynamic range to show the interesting deviations from this average.

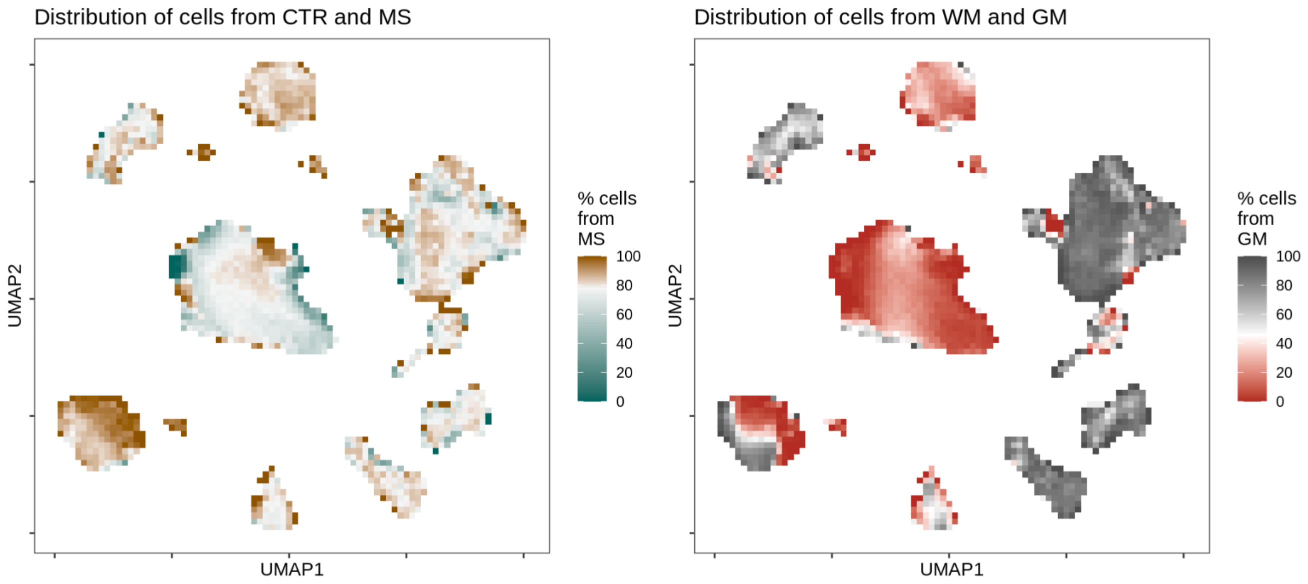

**ED3a** Heatmap of oligodendroglia clusters. OPC=oligodendrocyte precursor cell, COP=committed oligodendrocyte precursor, NFOL=newly formed oligodendrocyte, MFOL=myelin forming oligodendrocyte, MOL=myelinating oligodendrocyte, DA=disease associated, IFN-interferon signalling-related. Gene groupings derived from Falcão et al. and Hilscher et al. [16,79].

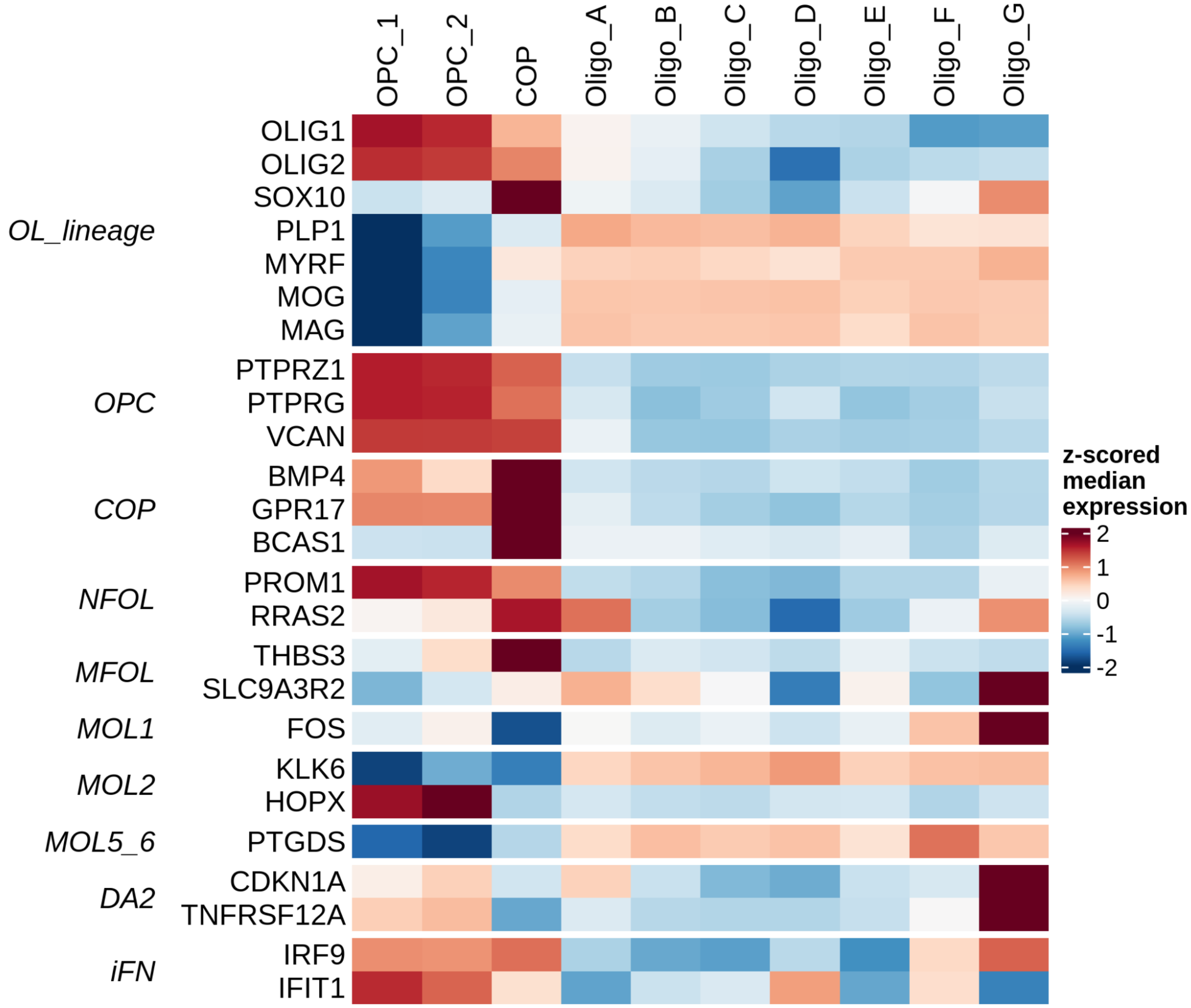

**ED3b Validation of number of GPR17-expressing cells**. Upper panel shows the number of cells with non-zero expression of GPR17 in each snRNAseq sample included in this study. Lower panel shows mean GPR17+ cells per mm2 (total calculated over 10 randomised fields) in an independent cohort of MS WM samples. In both panels: horizontal line denotes median. WM= control white matter, NAWM= normal appearing white matter in MS samples, AL=active demyelinated lesion, CAL=chronic active demyelinated lesions, CIL=chronic inactive demyelinated lesion, RL=remyelinated lesion. Examples of immunofluorescence using anti-GPR17 antibody (red) and DAPI (blue) for cell nuclei on brain white matter. Scale bar=10um.

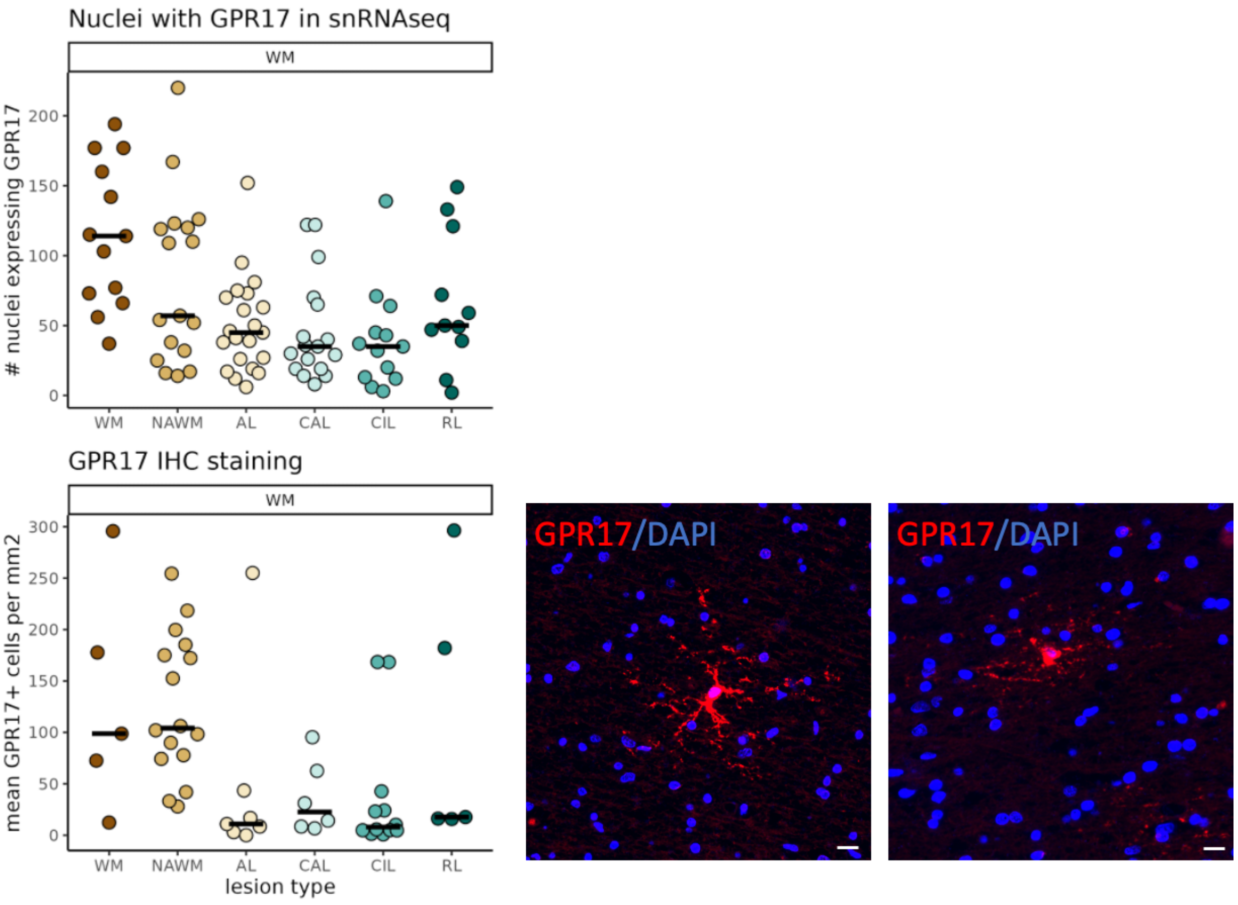

**ED3c** Heatmap of astrocyte clusters. TFs = transcription factors, synapse_fn = related to synaptic function, BBB=blood brain barrier.

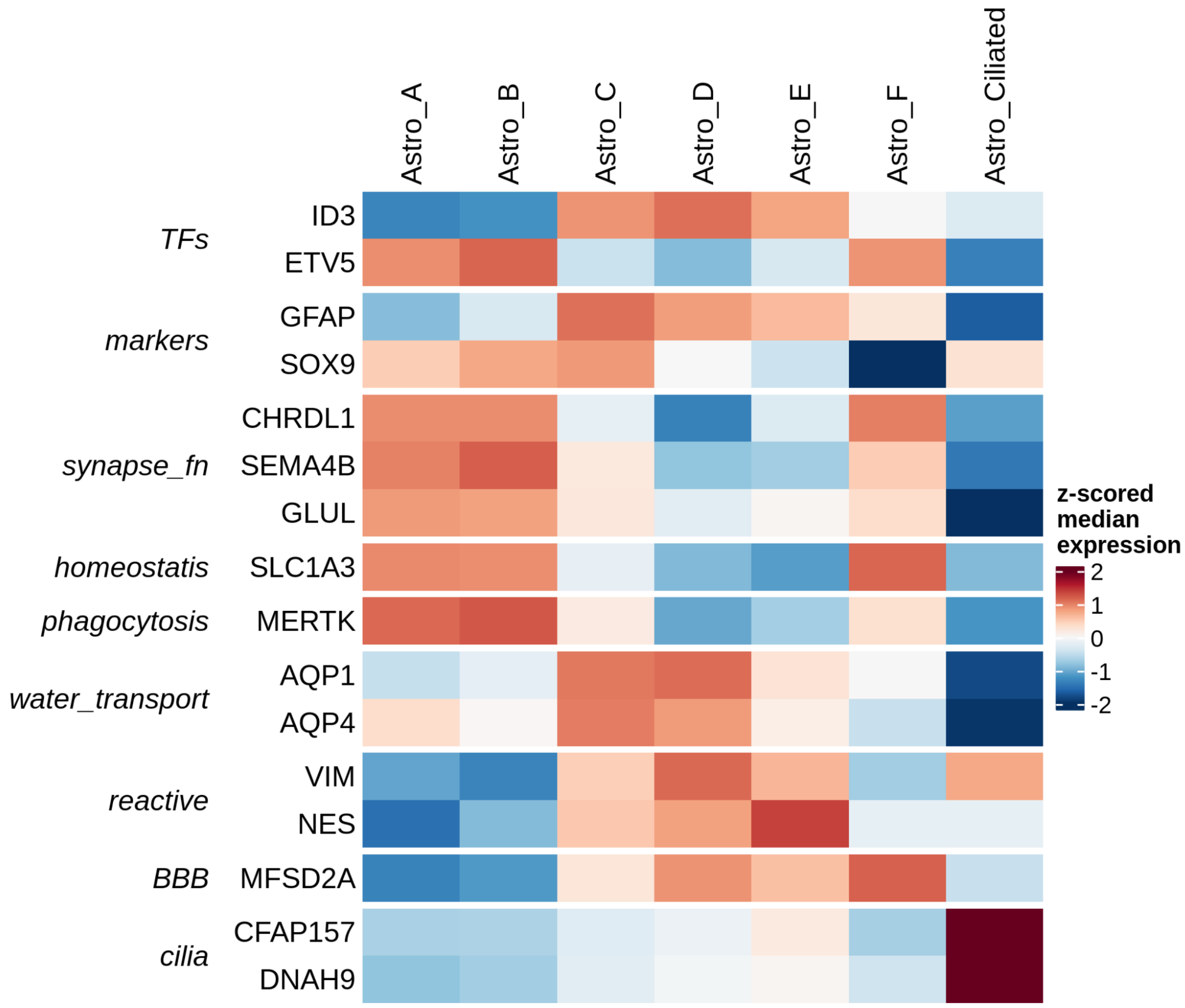

**ED3d** Heatmap of microglia clusters, divided into markers suggesting homeostatic or reactive phenotypes. PVM=perivascular macrophages.

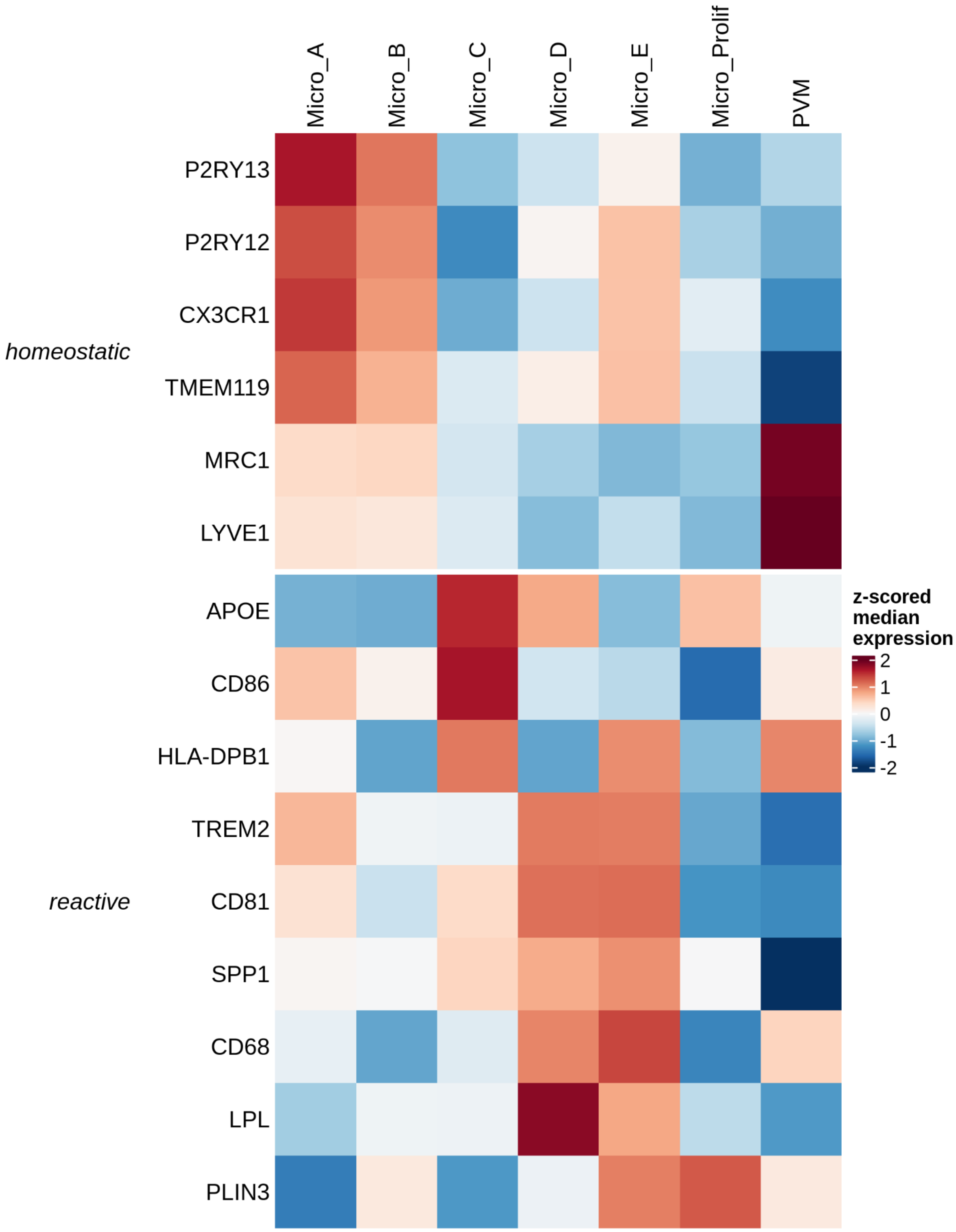

**ED4 Examples of the classical types of MS lesions** as defined by neuropathology on post-mortem MS brain using immunohistochemistry for myelin oligodendrocyte glycoprotein (MOG)(brown - A,C,E,G,I) to show lesions. Inflammatory infiltrate is labelled by antibodies to MHC-class II (brown) in combination with the histological stain luxol fast blue (LFB) identifying myelin (blue - B,D,F,H,J). Example of active (AL), chronic active (CAL), chronic inactive (CIL) and remyelinated (RL) white matter lesions and of one subpial grey matter demyelinated lesion (GML).

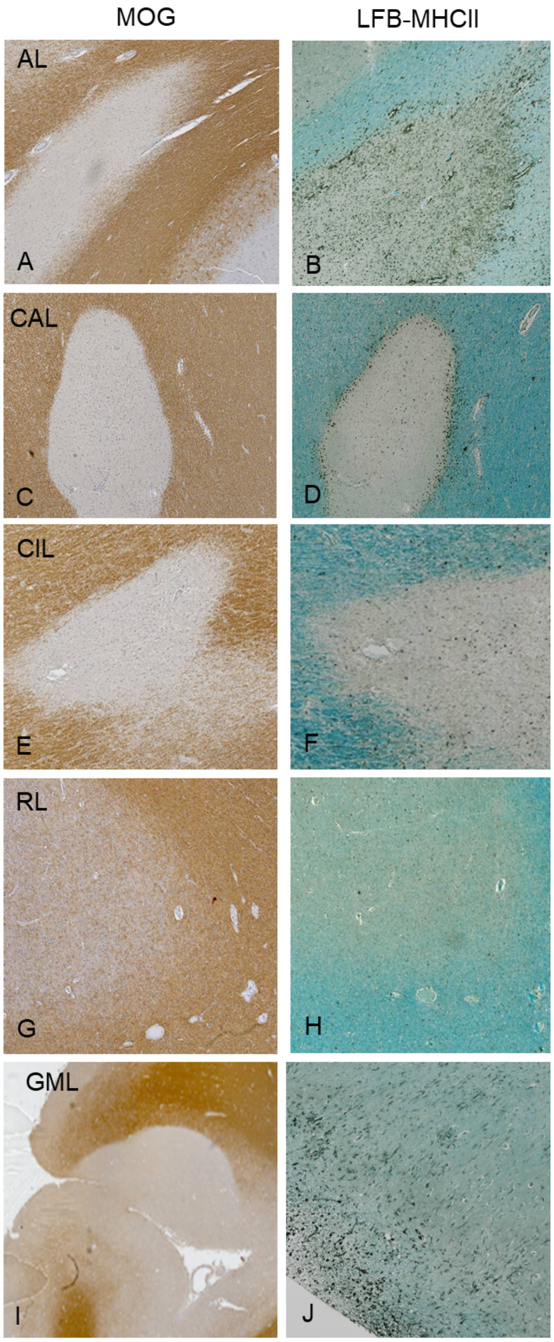

**ED5a** Milo compositional analysis for fine cell types in GM. Control pcs are shown to show the layer effects. Dots correspond to Milo neighbourhoods. Dots are coloured where the bootstrap procedure finds them to be distinct from zero (see Methods). Red - significantly increased, and blue - significantly decreased neighbourhoods in each column.

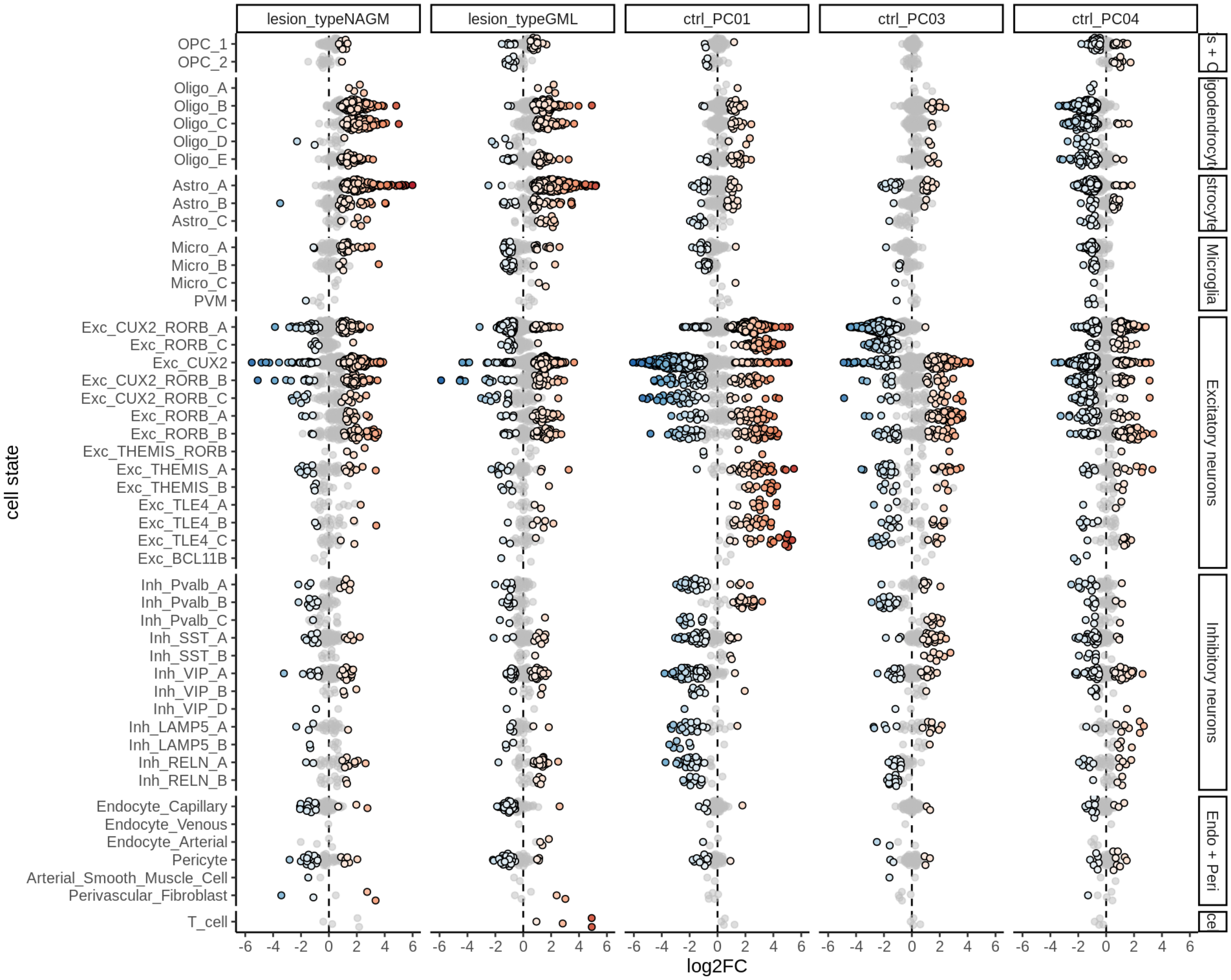

**ED5b** Milo compositional analysis for fine cell types in WM. Dots correspond to Milo neighbourhoods. Dots are coloured where the bootstrap procedure finds them to be distinct from zero (see Methods). Red - significantly increased, and blue - significantly decreased neighbourhoods in each column.

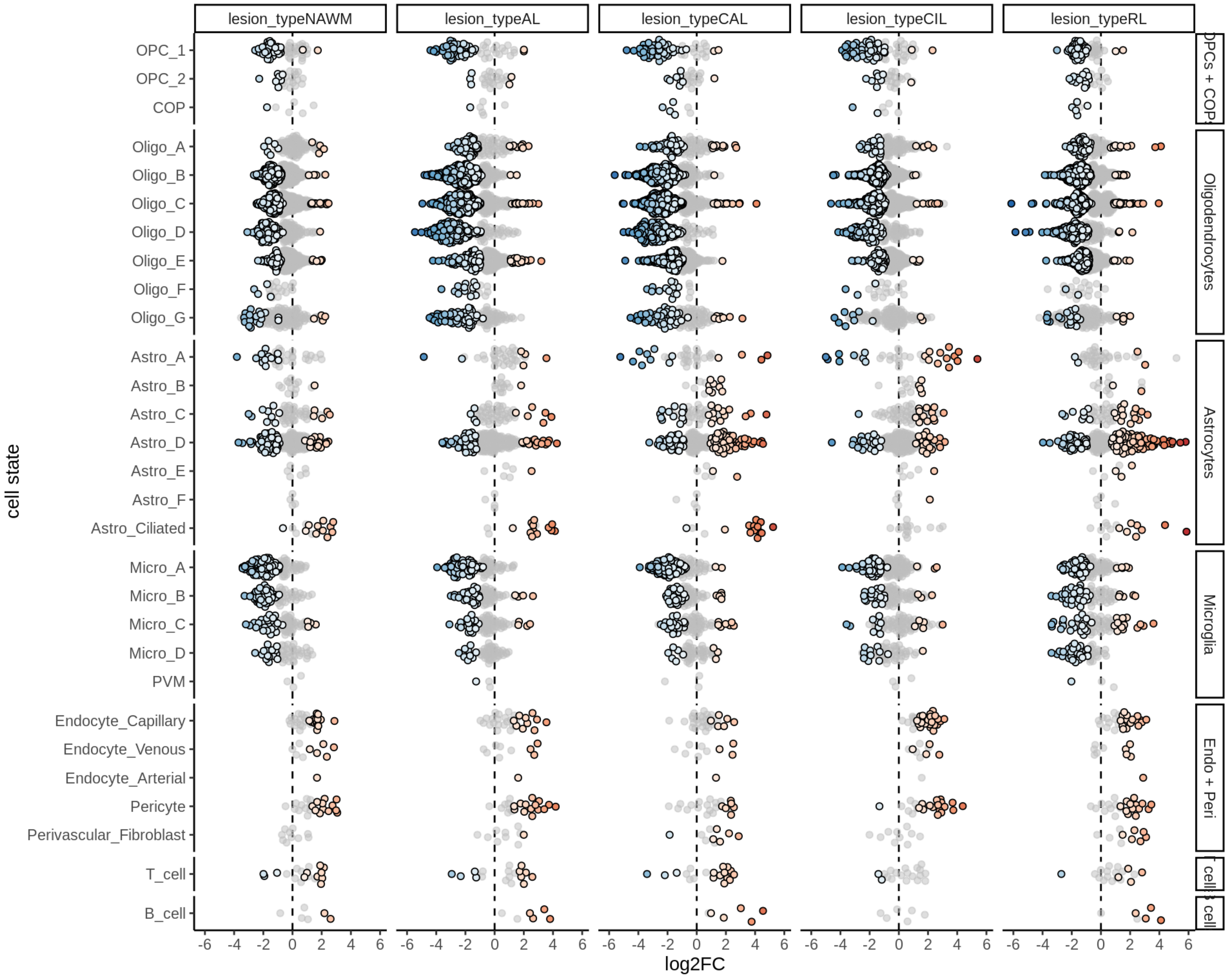

**ED6a** Distribution of model fits for genes for each broad celltype in GM. y-axis shows standard deviation of random (donor) effects for each gene. x-axis shows log2FC of lesion type with smallest p-value for each gene. Horizontal dashed lines show cutoff at *SD = log(1.5)*; vertical dashed lines show cutoff at *abs(log2FC) = log2(1.5)*.

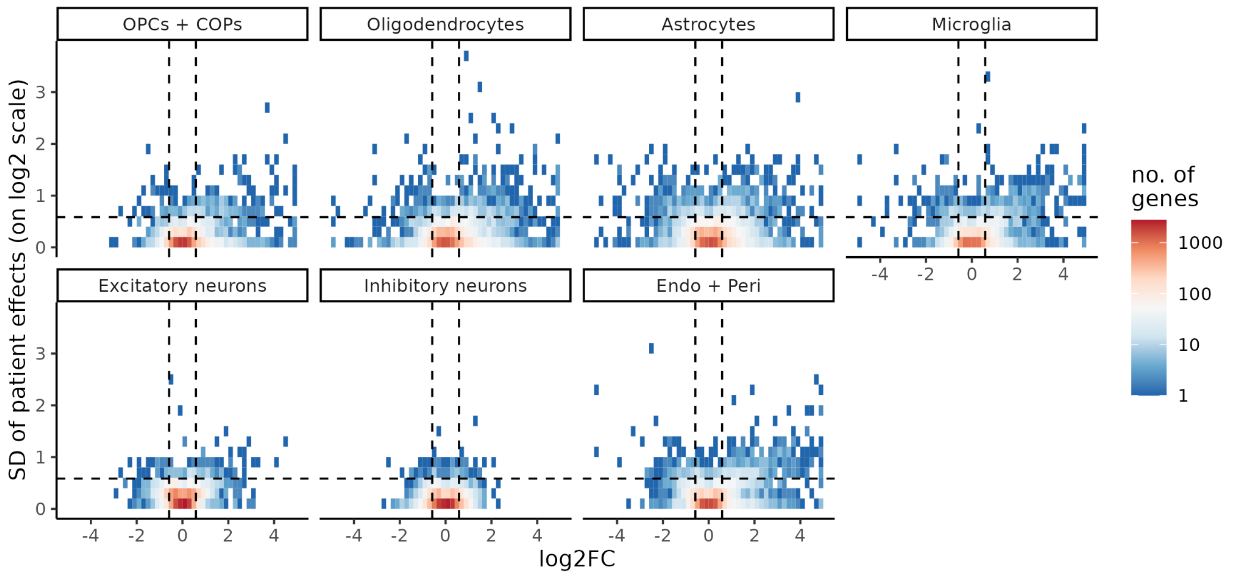


**ED6b** Distribution of model fits for genes for each broad celltype in WM; as for **ED6a**, but thresholded at *SD = log(2)* and *abs(log2FC) = log2(2)*.

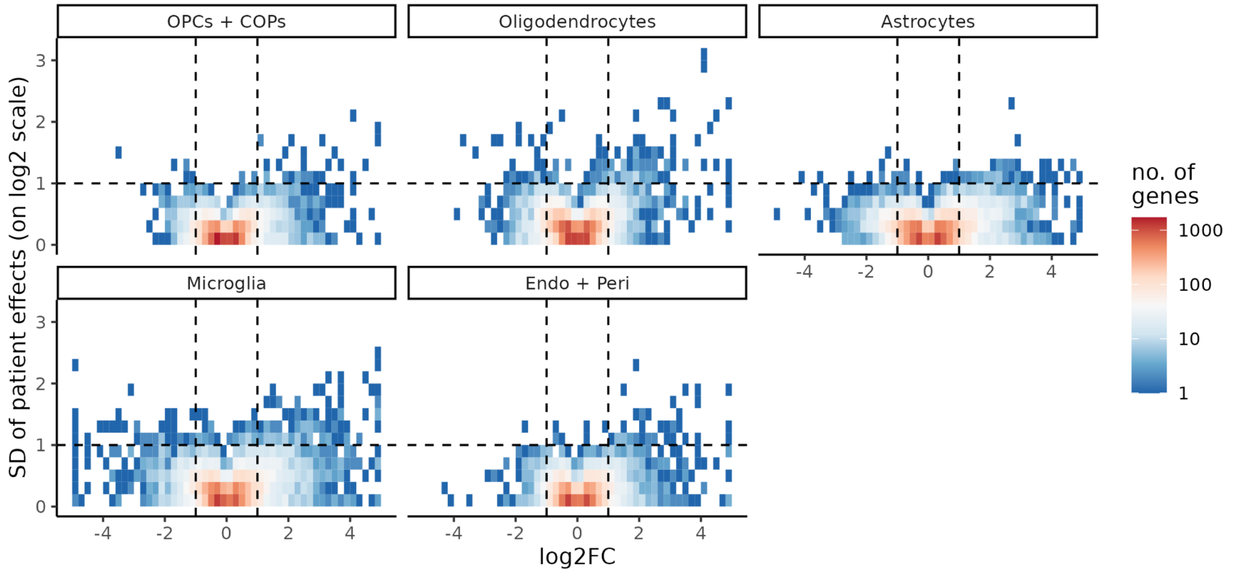

**ED7a** Pseudobulk expression heatmap of genes showing either MS or donor variability in broad cell types in GM, samples ordered by lesion type, showing differences between MS donors not explained by lesion type. Column order determined by the first principal component of the gene matrix for each cell type. Abbreviations: CTR = Control donor grey matter, NAGM = normal appearing grey matter, GML= grey matter demyelinated lesion, opc_cop = OPCs and COPs; oligo = oligodendrocytes; astro = astrocytes; micro = microglia, excit = excitatory neuron, inhib = inhibitory neuron, endo_peri = endothelial cells and pericytes.

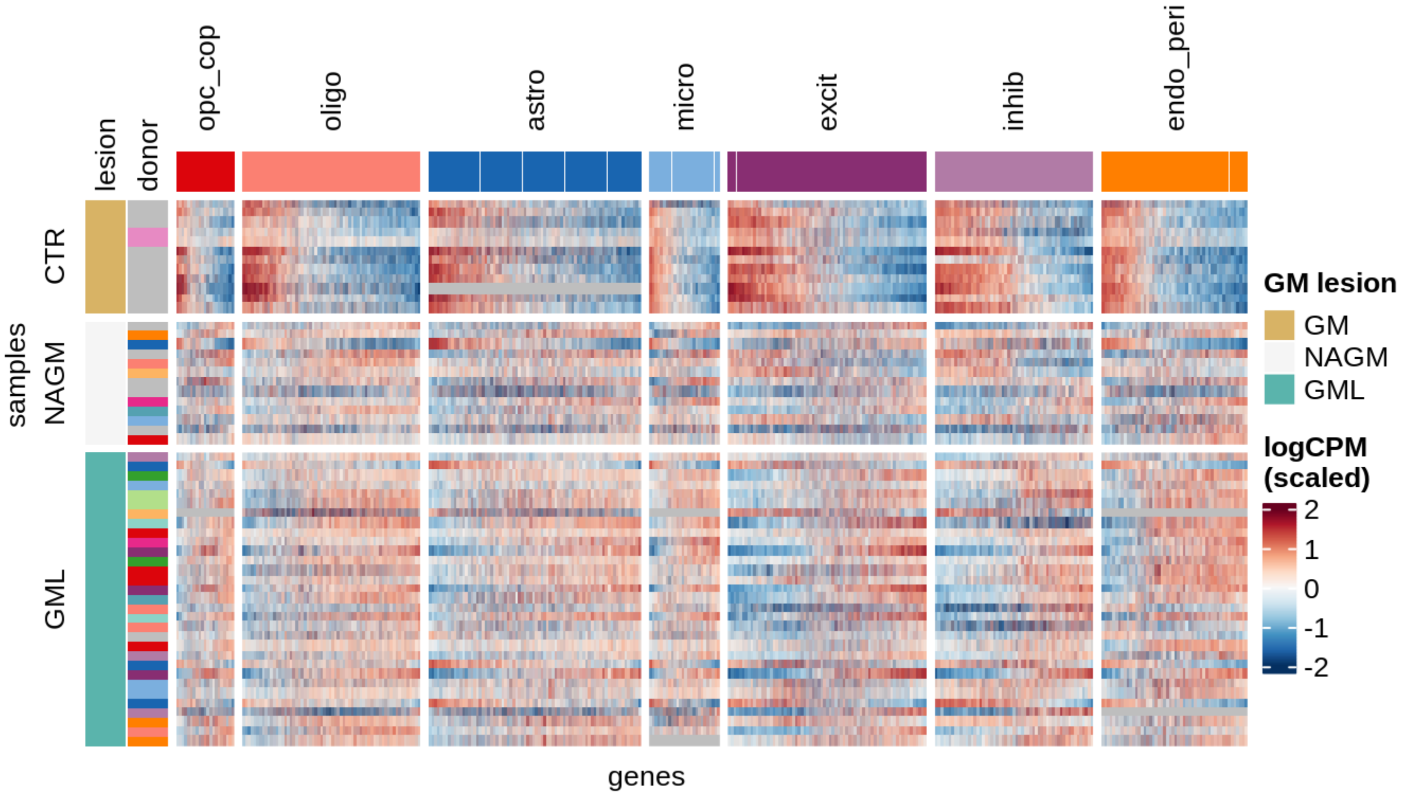

**ED7b** Pseudobulk expression heatmap of genes showing either MS or donor variability in broad cell types in GM, samples ordered on basis of hierarchical clustering, showing that all samples from one MS donor cluster together, suggesting a strong donor effect. Abbreviations: CTR = control donor grey matter, NAGM = normal appearing grey matter, GML = grey matter demyelinated lesion, opc_cop = OPCs and COPs; oligo = oligodendrocytes; astro = astrocytes; micro = microglia, excit = excitatory neuron, inhib = inhibitory neuron, endo_peri = endothelial cells and pericytes.

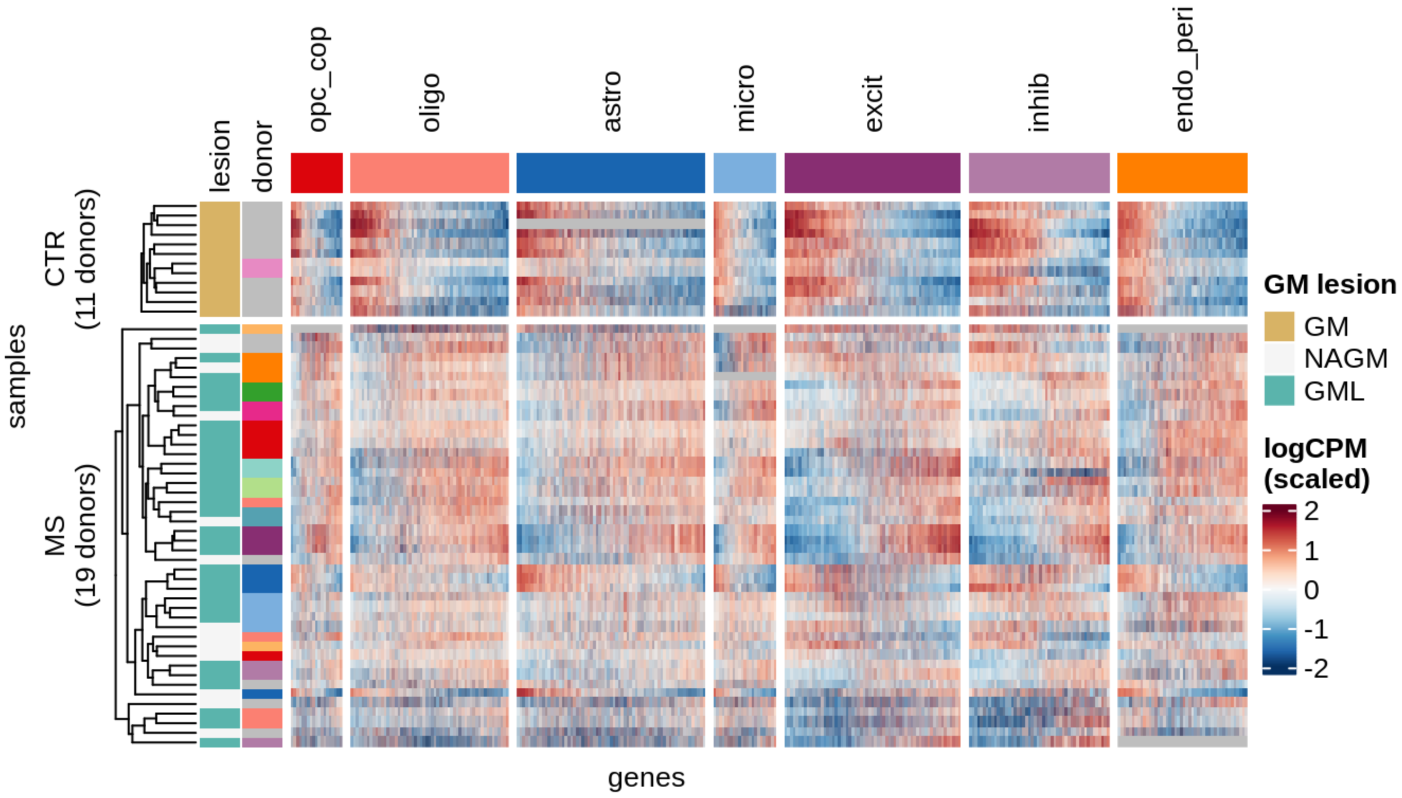

**ED8a** Extent of overlap of genes selected for each broad cell type for GM MOFA+.

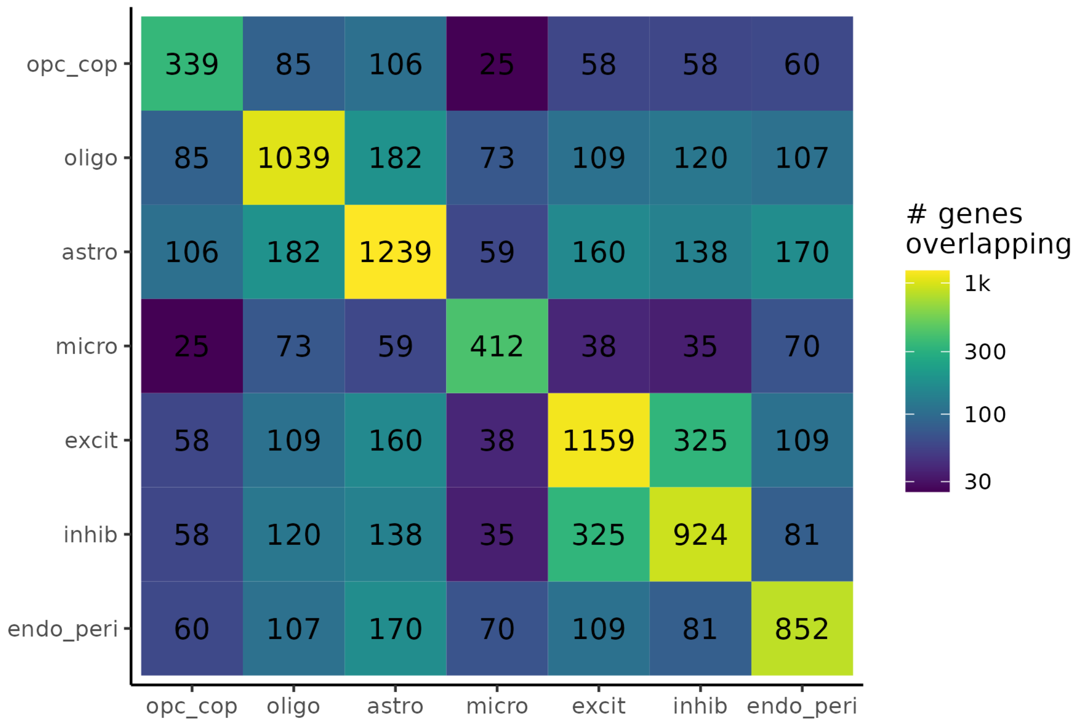


**ED8b** Extent of overlap of genes selected for each broad cell type for WM MOFA+.

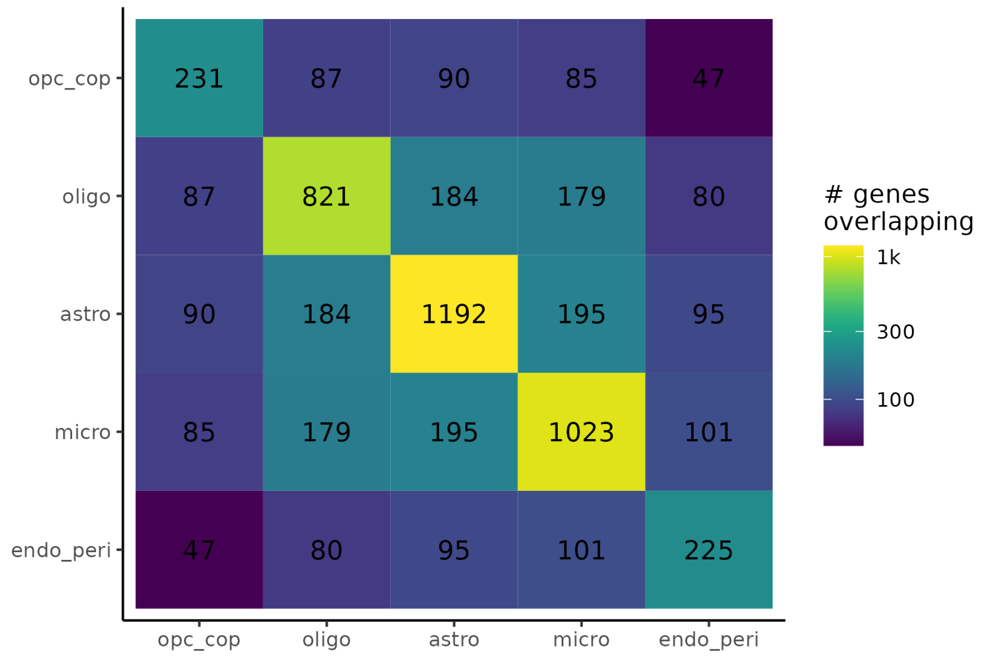

**ED9a** Distributions of MOFA+ factors in GM. In the lower panel, colour denotes donor; grey is used where only one sample was obtained from a donor.

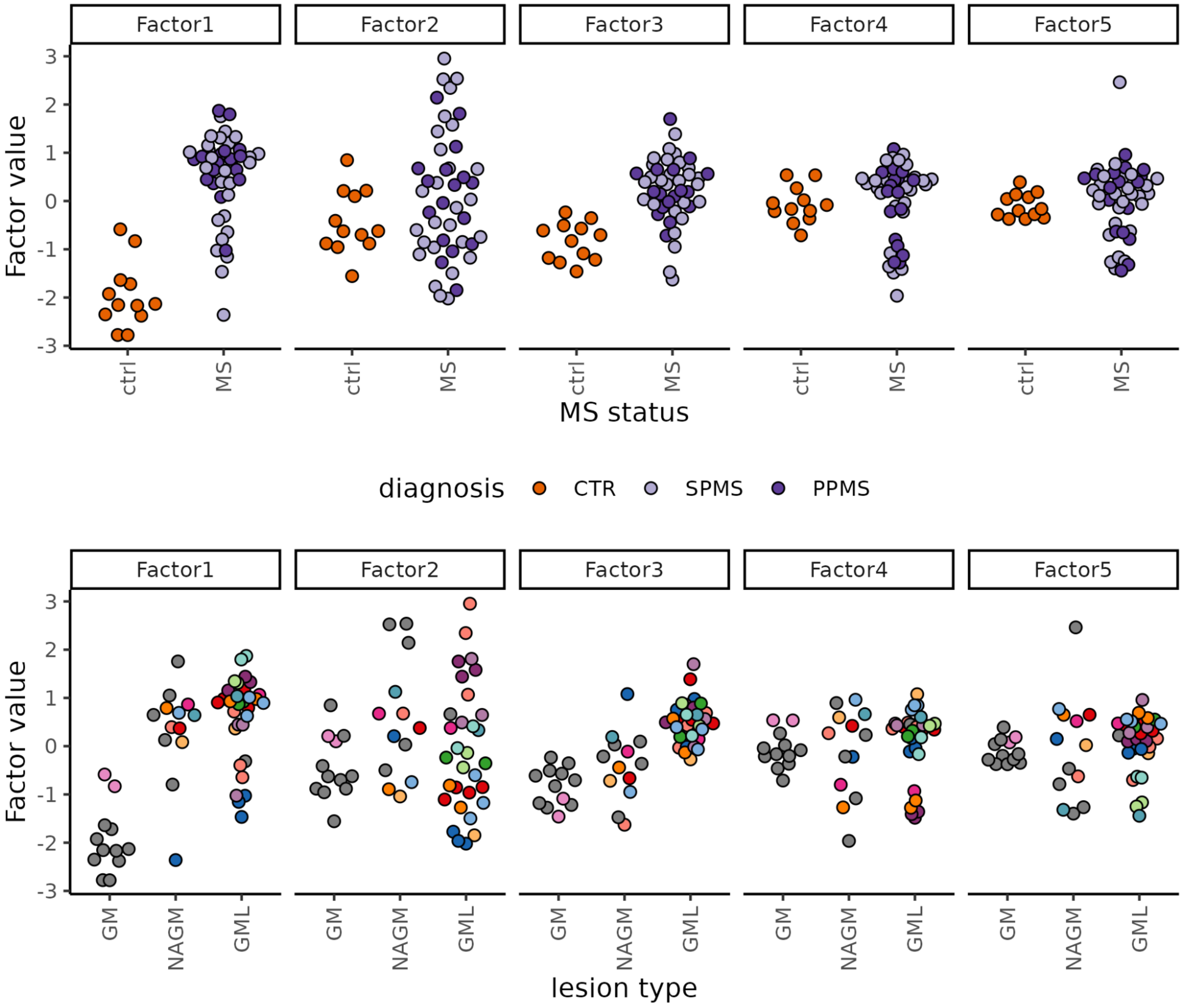

**ED9b** Variance in gene expression explained by MOFA+ factors, and extent to which MOFA+ factors are explained by metadata, in GM. Variance explained in first panel is per celltype (i.e. maximum possible total for each row is 100%). Pseudo-R2 values in second panel are calculated by fitting a mixed model to each factor, using formula *factor_value ~ lesion_type + sex + age_scale + pmi_cat + (1 | donor_id)*, and the *glmmTMB* function in package *glmmTMB*[27]. Pseudo-R2 values are determined by Nakagawa's R2[80], showing proportion of variance explained using fixed components only, and including a donor effect (see Methods).

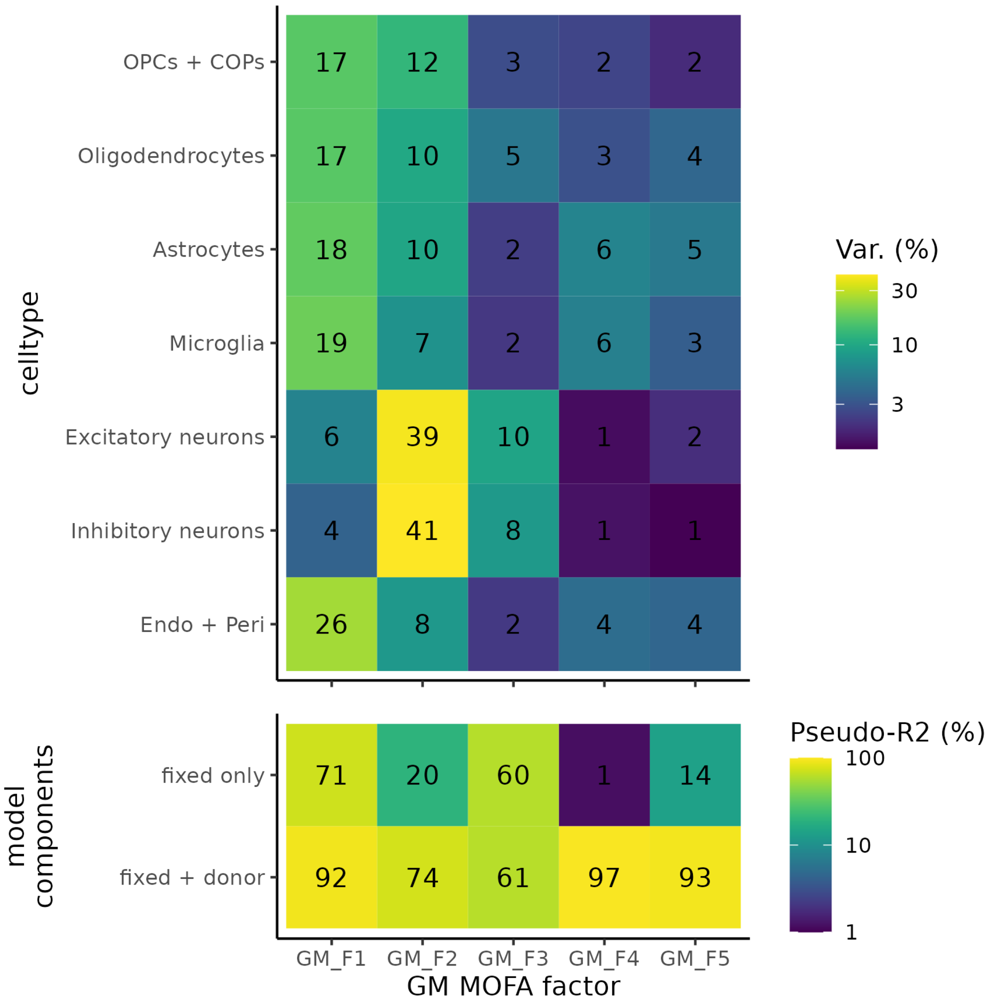

**ED9c** Geneset enrichment analysis of genes associated with GM MOFA factors (see Methods). Panels correspond to GM MOFA factors. Rows are the top 3 GO biological process genesets with the largest absolute nomalized enrichment scores (NES) per combination of factor and broad cell type, subject to FDR < 10%. These overlap, resulting in fewer genesets displayed than the maximum possible (75). NES is a measure of how concentrated the significant genes are in a given set. padj = Benjamin-Hochberg adjusted p-value of enrichment.

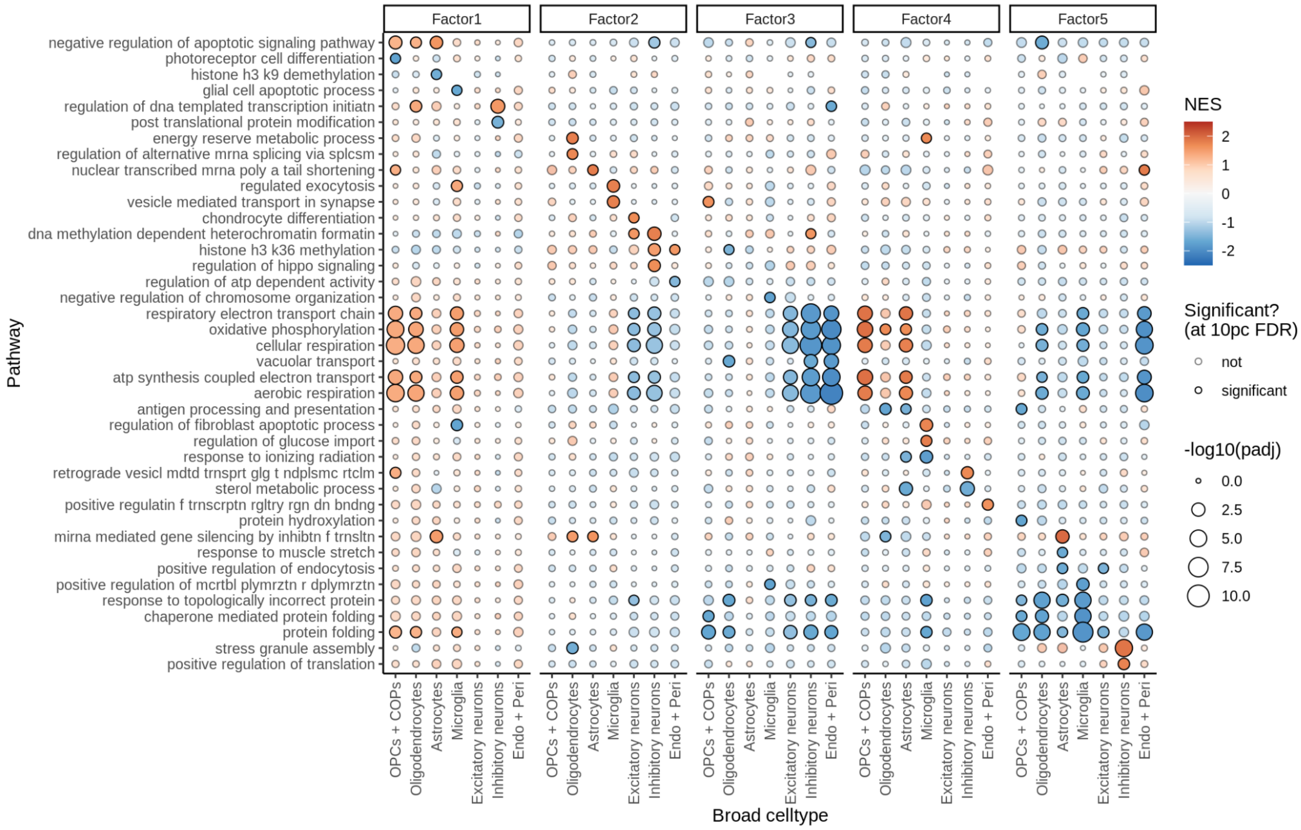

**ED9d** Comparison of WM MOFA factors with factors from scITD (see Methods for calculations). The y-axis corresponds to WM MOFA factor scores, with each row indicating a different MOFA factor. The x-axis corresponds to scITD factor scores, with each column indicating a different scITD factor. For each MOFA factor, we show the Spearman correlation coefficient for the scITD factor with the highest correlation.

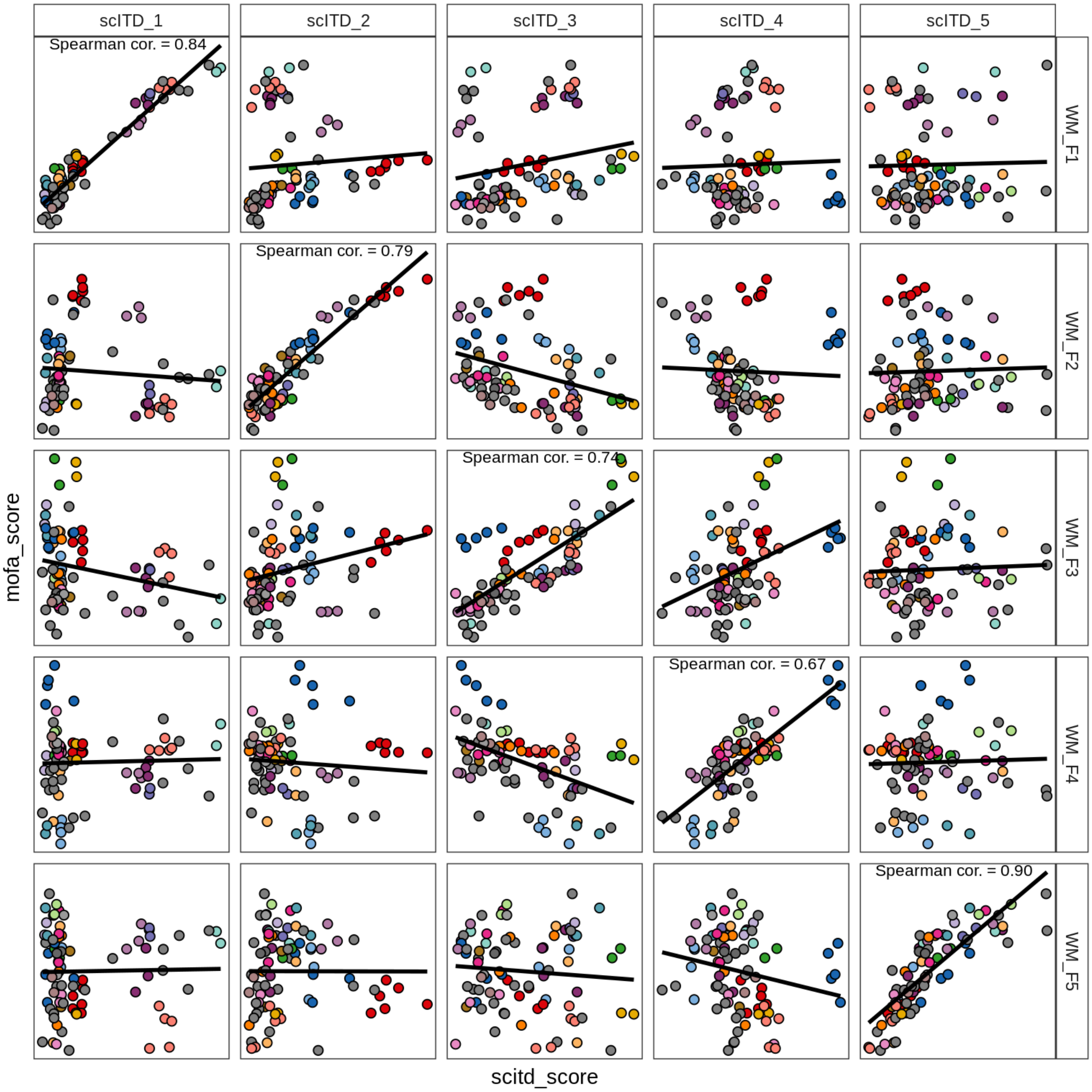

**ED9e** WM MOFA factors annotated by sample metadata. Rows correspond to samples, ordered by hierarchical clustering of the factor scores. Columns correspond to factors. Left hand annotations show lesion type and donor (donor annotation has grey where a donor only has one sample in our dataset). Right hand annotation shows metadata. brn_bank = brain bank, batch = pool of samples sent for sequencing together.

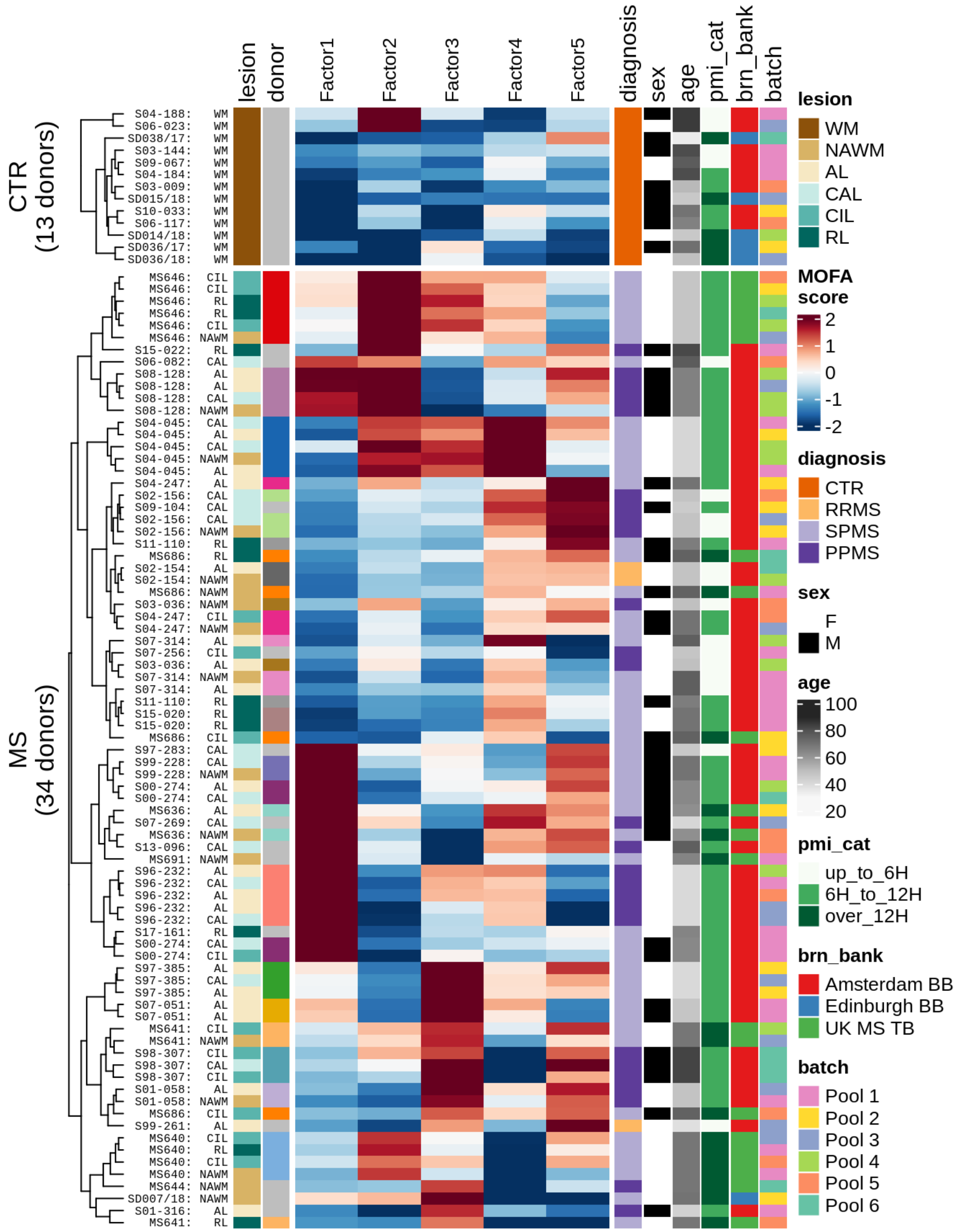

**ED9f** Heatmap of MOFA+ factors in WM (left), annotated with median QC metrics per sample (right). **Left** Rows are samples, annotated by metadata and oligo groupings; donor is colour for donor ID, with grey values used for donors contributing only one sample. Columns are MOFA+ factors, with signs changed to positively correlate with MS status. **Right** Columns are QC metrics: mito pct is the proportion of reads in the sample that are mitochondrial; pct unspliced is the proportion of reads in the sample that are unspliced as opposed to spliced mRNA; neuronal pct is the percentage of cells in the sample that are neurons. Colours in heatmap are the z-scores for each QC metric column, with colours chosen so that dark purple is ‘bad’ (e.g. low library size, or high mitochondrial read percentage).

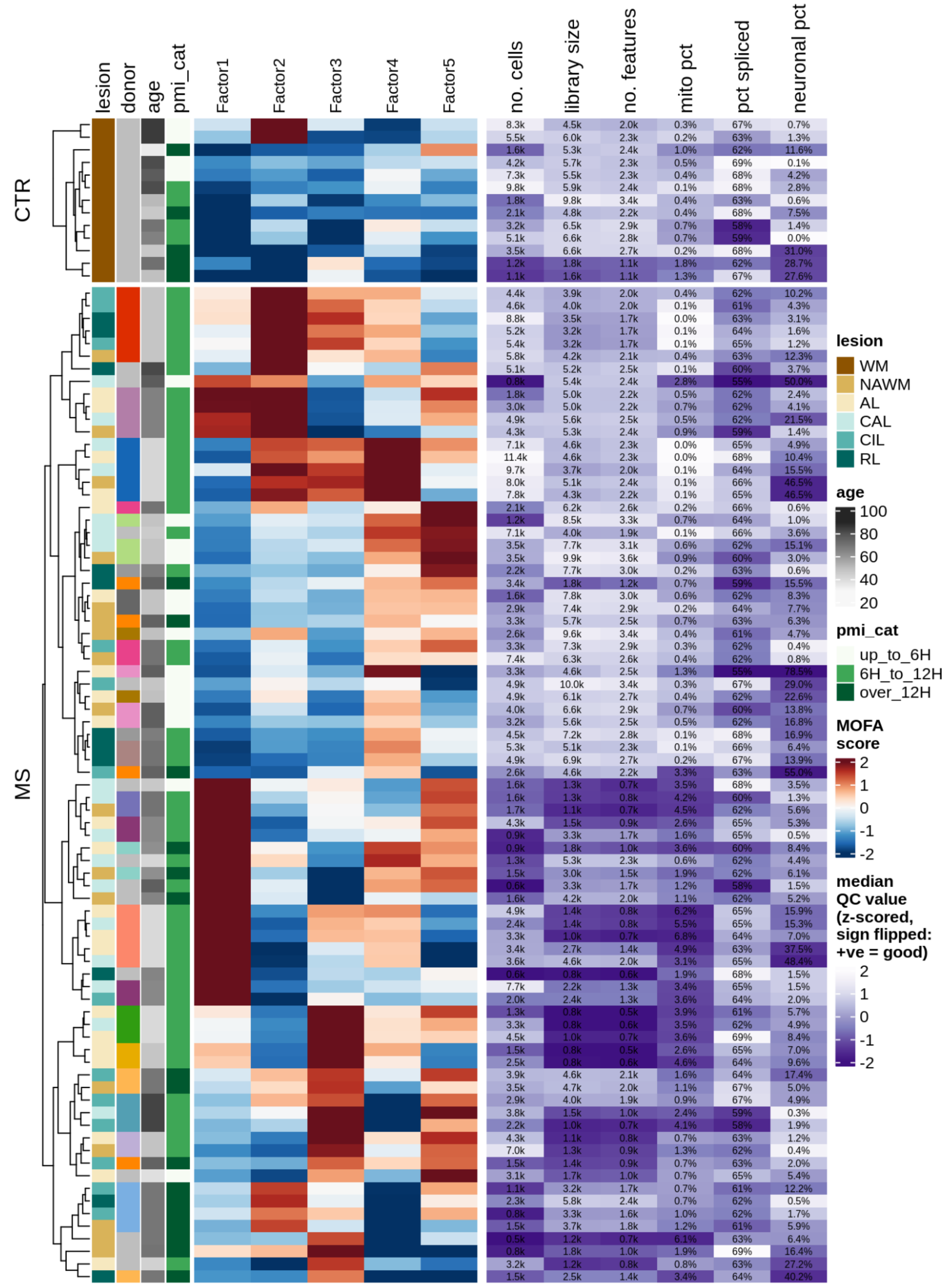

**ED 9g** Geneset enrichment analysis of genes associated with WM MOFA factors (see Methods). Panels correspond to WM MOFA factors. Rows are the top 3 GO biological process genesets with the largest absolute nomalized enrichment scores (NES) per combination of factor and broad cell type, subject to FDR < 10%. These overlap, resulting in fewer genesets displayed than the maximum possible (75). NES is a measure of how concentrated the significant genes are in a given set. padj = Benjamin-Hochberg adjusted p-value of enrichment.

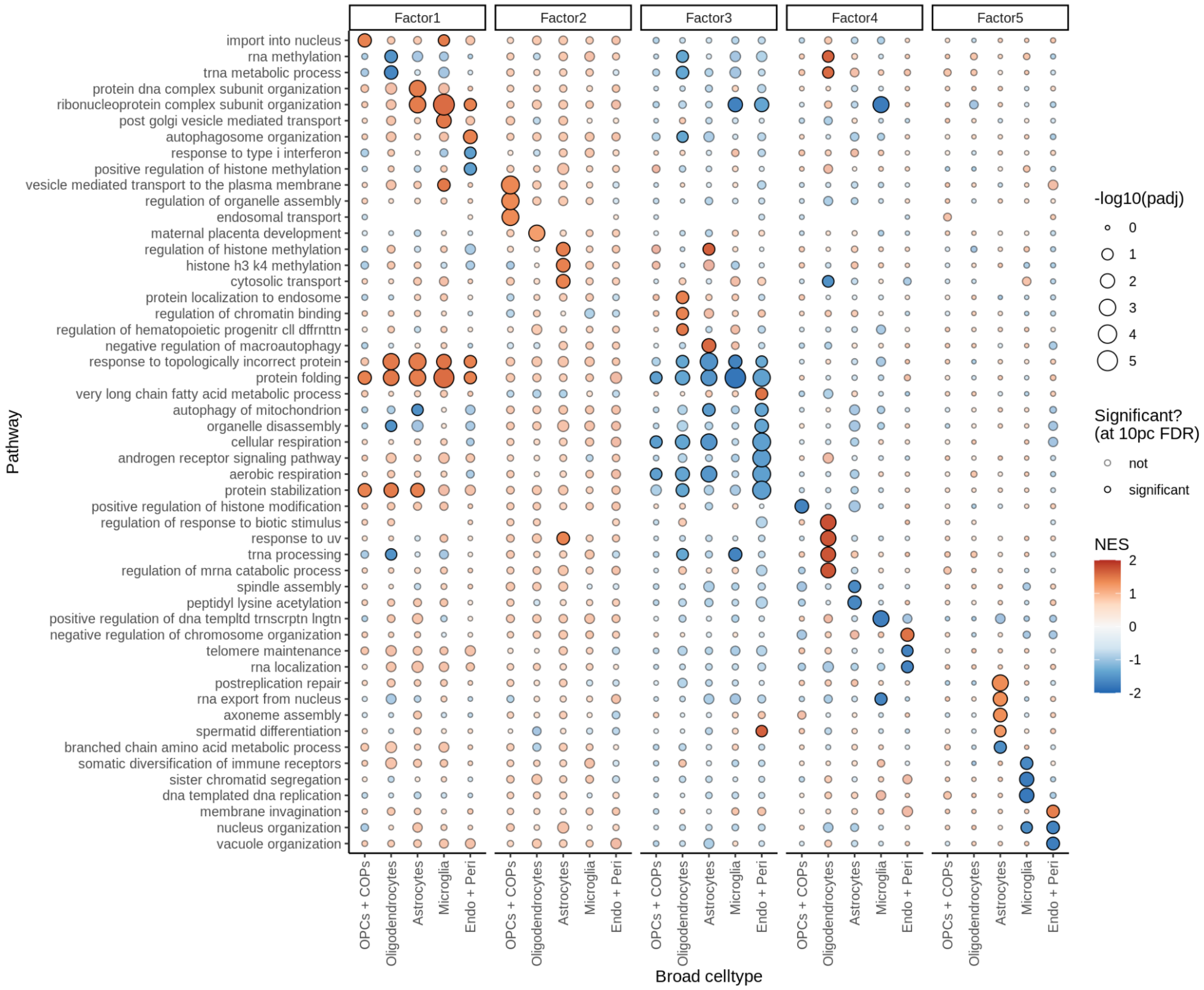

**ED10a** Proportions of neuronal compartment per sample, split by layer-specificity of neurons. L1 and L2/L3 neurons account for relatively low proportions of NAGM samples, while L5 and L6 neurons account for high proportions of NAGM samples; vice versa for GML samples, while control GM lies in the middle. This indicates that, on average, the samples are roughly ordered as follows: NAGM is deeper than control GM, which is deeper than GML.

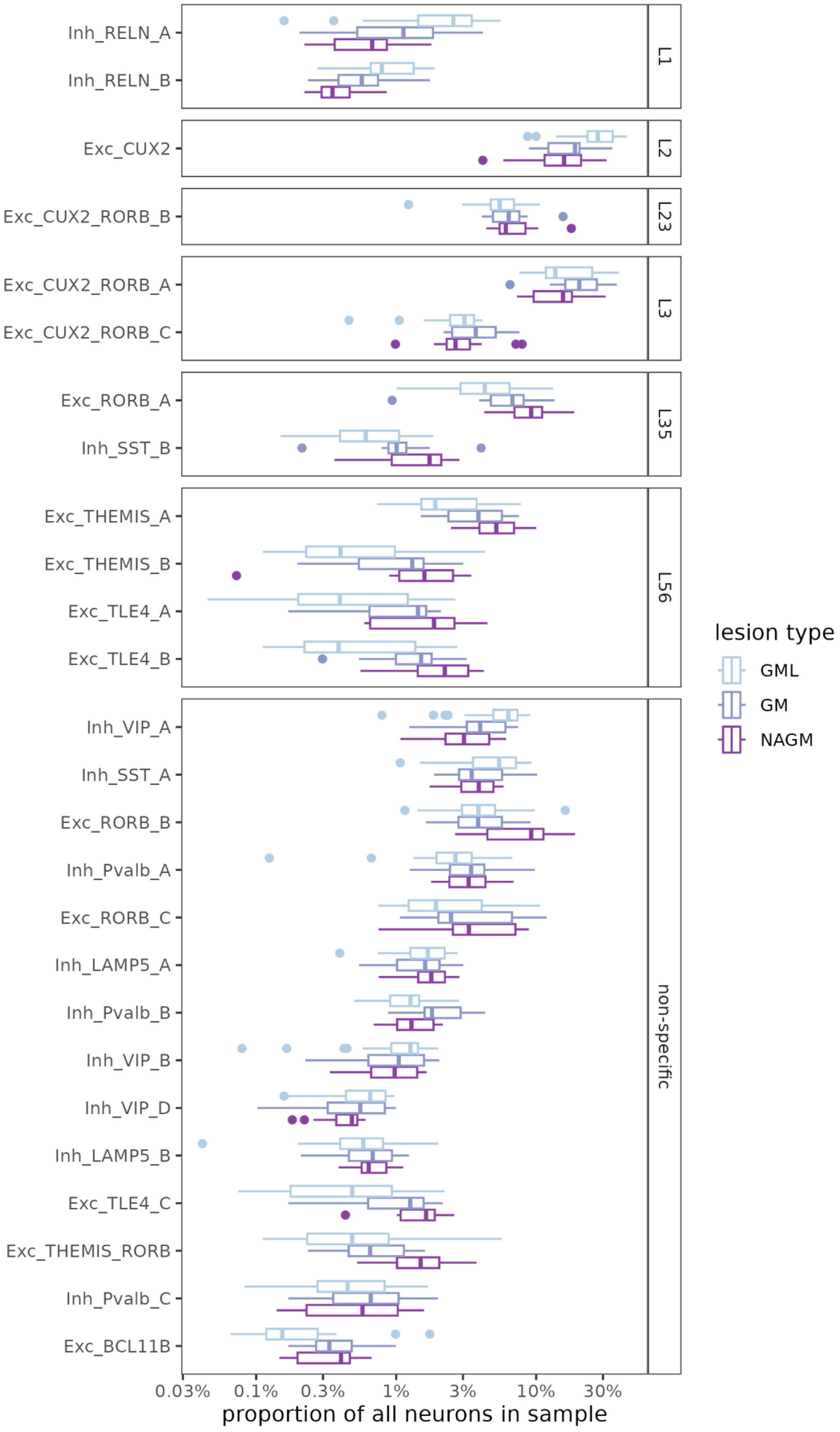

**ED10b** Principal components of GM neuronal layer centred log ratios (CLRs; see Methods), applied to control GM samples only. y-axis shows absolute Spearman correlation between PC loadings and neuronal layer numbers (i.e. L1-L6 corresponds to numbers 1-6); neuronal clusters without an assigned layer number are excluded. x-axis shows the variance explained by each PC (on a log scale). Dashed lines show thresholds at 0.2 Spearman correlation, and 1% variance explained, giving 4 PCs that could be relevant to layers.

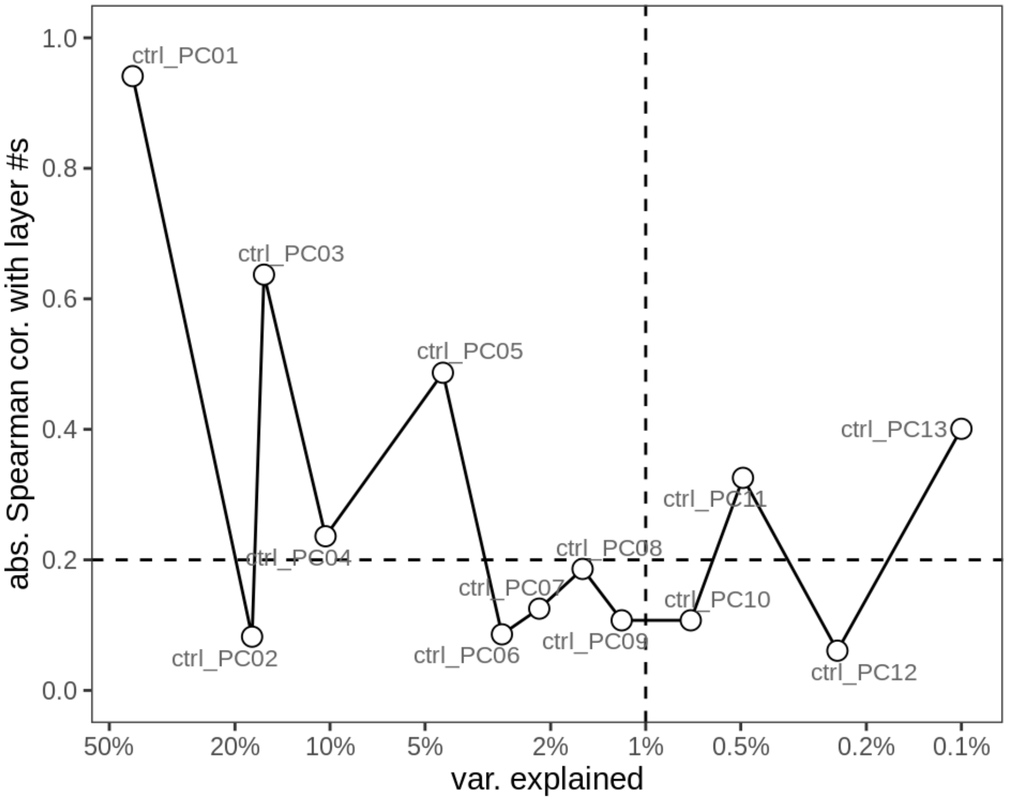


**ED10c (next page)** Bootstrapped ANCOM-BC results for GM including varying numbers of PCs as covariates. Number of PCs used varies from 0 to 8 (see **ED10b** for selection of possible PCs). Point shows median bootstrapped value; coloured lines show 80% bootstrapped confidence intervals; grey lines show 95% bootstrapped confidence intervals. Points are filled when the 95% CI excludes zero; otherwise empty. All intervals based on 20k bootstraps.

Neurons only

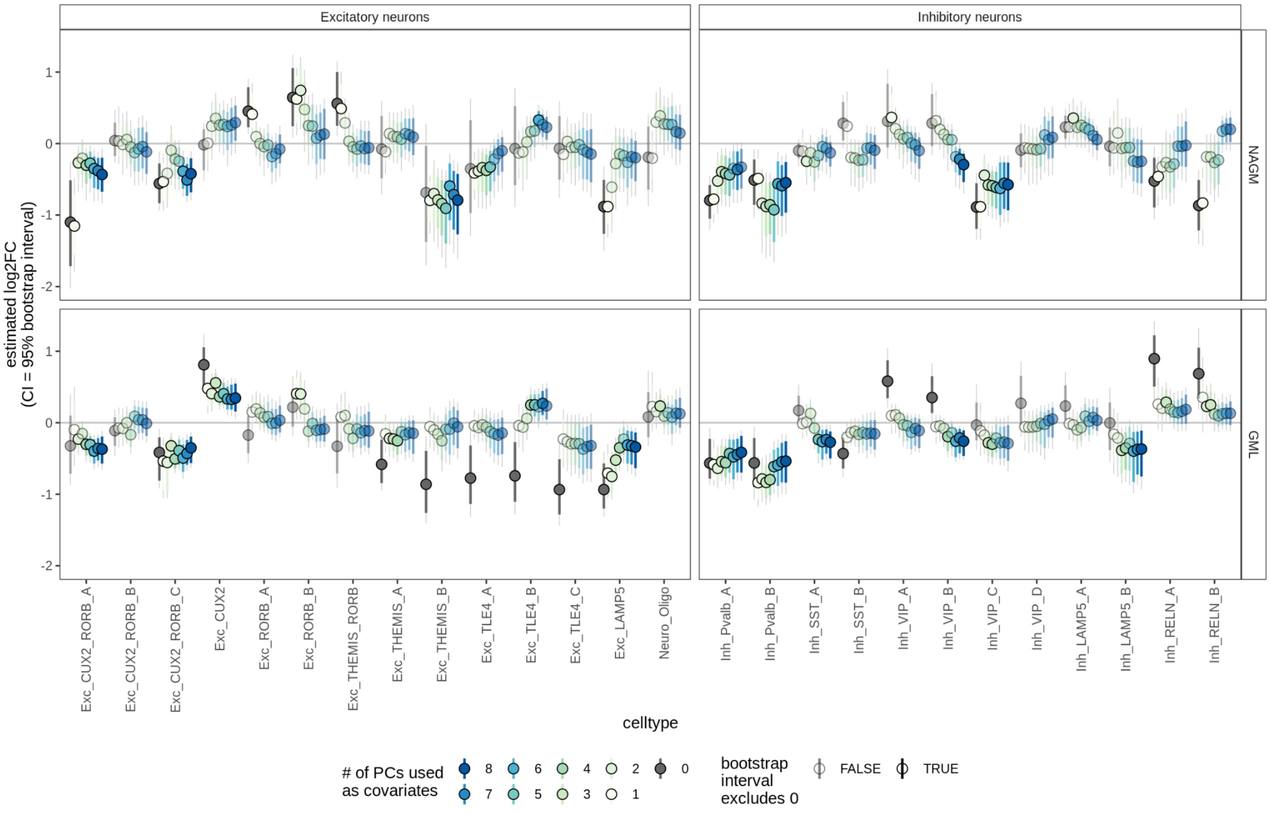


Non-neurons only

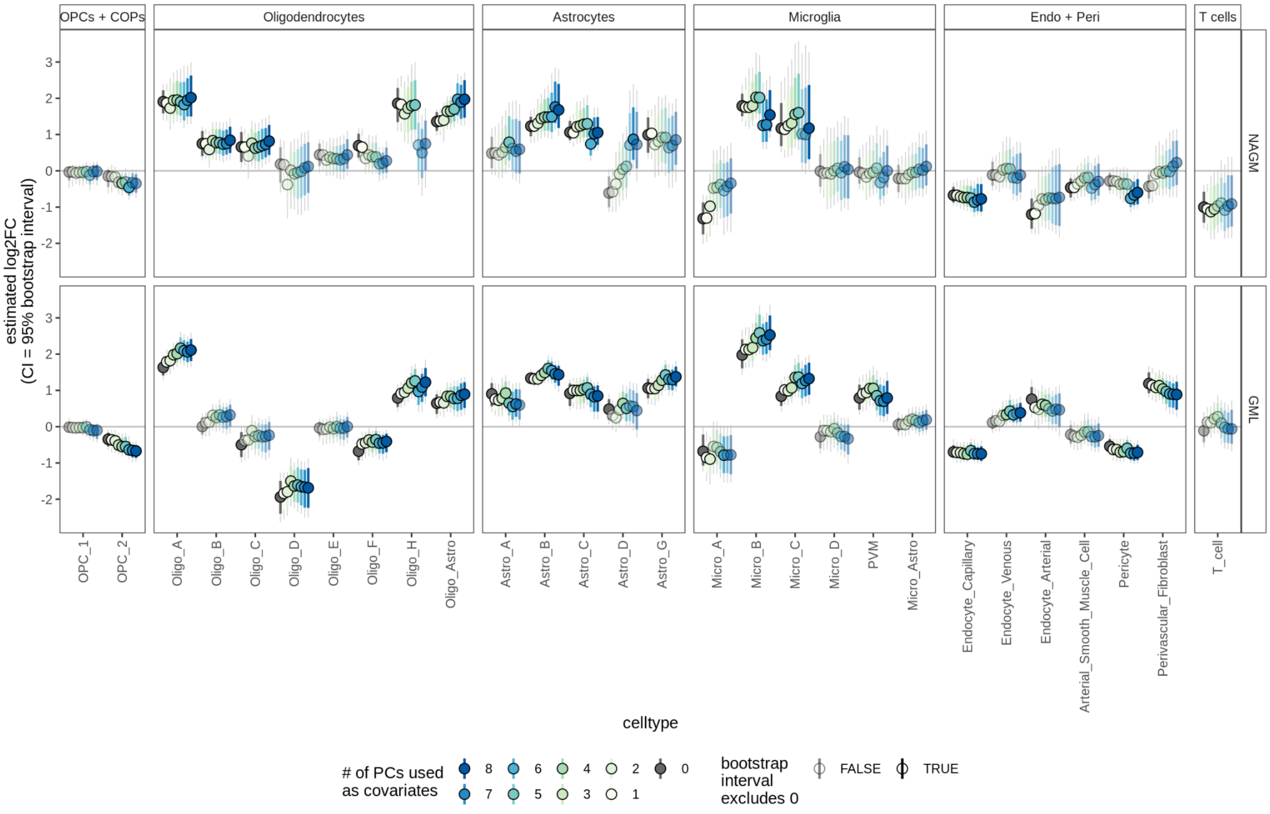

### Extended Data Files

Extended Data File 1: Sample-level metadata and QC metric summary

Extended Data File 2: Alternative QC and clustering pipeline

Extended Data File 3: Marker genes for all broad and fine cell types

Extended Data File 4: Differentially expressed genes for all celltypes, both WM and GM (fine and broad level)

Extended Data File 5: GSEA results for GM and WM MOFA factors

### Extended Data Tables

**Extended Data Table 1**

Results of chi-squared tests of independence between case-control status and four test metadata variables, within WM and GM. FDR is Benjamini-Hochberg corrected p-value.

| Matter | Reference variable | Test variable | Chi-squared statistic | Degrees of freedom | p-value | FDR | Is significant? (at 5%) |
|---|---|---|---|---|---|---|---|
| WM | Case-control | Age category | 3.95 | 2 | 1.40E-01 | 1.80E-01 | FALSE |
| WM | Case-control | Sex | 1.45 | 1 | 2.30E-01 | 2.30E-01 | FALSE |
| WM | Case-control | PMI category | 4.49 | 2 | 1.10E-01 | 1.80E-01 | FALSE |
| WM | Case-control | Sample source | 27.31 | 2 | 1.20E-06 | 4.70E-06 | TRUE |
| GM | Case-control | Age category | 9.65 | 2 | 8.00E-03 | 1.50E-02 | TRUE |
| GM | Case-control | Sex | 0.53 | 1 | 4.70E-01 | 4.70E-01 | FALSE |
| GM | Case-control | PMI category | 9.01 | 2 | 1.10E-02 | 1.50E-02 | TRUE |
| GM | Case-control | Sample source | 58.78 | 1 | 1.80E-14 | 7.10E-14 | TRUE |

### Supplementary note

#### Contribution of model elements to cell type abundance variability

To test whether lesion type and donor ID explain the cell type proportion across samples, we fit a series of nested models of increasing complexity for each cell type:

- null (*counts ~ 1*);
- covariates only full (*counts ~ sex + age_norm + pmi_cat*);
- fixed full (*counts ~ lesion_type + sex + age_norm + pmi_cat*); and
- full (*counts ~ lesion_type + sex + age_norm + pmi_cat + (1 | donor_id)*).

We use the R package *glmmTMB* (v1.1.2.2) to fit a negative binomial distribution for each model to the raw counts. We then use likelihood ratio tests (via the *anova* function) to test whether the more complex model increases the goodness of fit, i.e. whether adding this variable improves the model more than we would expect by chance. We adjusted the p-values across these tests for all cell types using the Benjamini-Hochberg procedure (i.e. across 3 * 50 p-values).

Note that we model raw counts rather than cell type proportions. An implicit assumption here is that there is no consistent bias in sample sizes between conditions or donors. This is a strong assumption, however this analysis is used only to make a weak conclusion, that donor variability is an important factor for some cell types.

If we model cell type proportions rather than raw counts, we introduce difficulties due composition (i.e. proportions summing to 1). Here, an increase in the proportion of one cell type results in a decrease in the proportion of all other cell types. Treating the data as compositional makes interpretation much more challenging, as differences in proportion could equally be caused by an increase in absolute abundance of one cell type, or a decrease in absolute abundance of a different cell type. In particular, it makes it difficult to make conclusions such as "abundance of cell type A varies considerably between MS patients", as this apparent effect may be mediated by another cell type. The true explanation could be "abundance of cell type B varies considerably between MS patients; this results in high patient-patient variability in the proportion of cell type A".

### Differential abundance of cell types in MS lesions and control samples

Differential abundance involves testing whether the abundance of a given group changes consistently between conditions; in our case, this indicates whether particular cell types are enriched or reduced in MS lesions vs controls. Such tests are subtle: to account for experimental variability, we would like to normalise by the total number of cells, and work with proportions. However, in the case where one cell type increases in abundance and all other cell types remain unchanged, the proportions of the other cell types will decrease, and vice versa.

Various methods have been developed to address this point. They principally use one or more cell types whose abundance is assumed to remain unchanged between all samples, and can therefore be used as a reference. ANCOM-BC is a method developed for microbiome data that uses a mixture model to computationally identify cell types which remain unchanged, and does not require users to specify a reference cell type [63]. An independent benchmarking study (of microbiome data) found ANCOM-BC to consistently be one of the best-performing currently available methods [81]. scCODA is a method developed specifically for single cell data, however it requires users to specify one unchanging cell type as reference[82]. As we do not know *a priori* which cell types are unaffected by MS, we used ANCOM-BC, and were unable to cross-check the results against scCODA.

### Differential expression analysis using generalised linear mixed models

Recent work has indicated that the primary limitation in differential expression analysis is sampling from a small number of individuals[67,83], and that pseudo-bulk approaches such as `muscat`[66] (that work at the level of the transcript totals across all cells of a given type in each sample) offer a good compromise between sensitivity and run time constraints. We therefore considered several pseudobulk approaches for analysing differential expression between our comparisons of interest. After analysis of the options, we chose a negative binomial model with a random effect, fit to count data with the glmmTMB package[27].

Initially, we ran muscat with `edgeR`[68], using the formula *counts ~ lesion_type + sex + age_scale + pmi_cat*. Here, *age_scale* is patient age, normalised to have SD = 0.5[56], and *pmi_cat* is post-mortem interval, split into three categories (under 1 hour, between 1 and 12 hours, and more than 12 hours). However, we found that for some genes there were substantial patient effects, i.e. genes where the donor ID was a much stronger determinant of expression than lesion type. Unfortunately `edgeR` does not at present allow random effects models.

We then investigated using `dream` from the `variancePartition` package[84], however we found that it returned extremely low numbers of results for cell types where the library sizes were low. In particular, this resulted in 2 significant results being reported for immune cells, compared to over 200 (biologically expected) genes reported for the same comparison with the negative binomial model. This may be due to `dream` operating on logCPM values, which do not explicitly model the variability of small counts. Using a mixed model also has the advantage of returning a value for each donor random effect (i.e. donor) in the study, which can be interesting for downstream analysis.

We therefore used a generalised linear mixed model, to allow both random effects and properly model count data. We used the `glmmTMB` function from the `glmmTMB` package[27], with a negative binomial model, and *donor_id* as a random effect. The formula for WM was *counts ~ lesion_type + sex + age_scale + pmi_cat + (1 | donor_id)*; the formula for GM was *counts ~ lesion_type + sex + age_scale + pmi_cat2 + (1 | donor_id)*, where *pmi_cat2* has only two categories (between 1 and 12 hours, and more than 12 hours). We included an offset of *log(lib.size) - log(1e6)*, so that the reported coefficients correspond to log counts per million (logCPM).

For a small number of genes, there were zero counts for the control condition. Where this occurs, `glmmTMB` does not fit properly (it operates at the log scale, and such a situation corresponds to the mean of one condition equalling log(0), i.e. minus infinity). Where an all-zeros condition occurred, we therefore added a count of 1 to the sample in the all-zeros condition with the largest library size. This is the smallest possible perturbation that avoids complete separation of the data.
